# Transient demographic dynamics of recovering fish populations shaped by past climate variability, harvest, and management

**DOI:** 10.1101/2023.03.22.533437

**Authors:** Daisuke Goto

**Affiliations:** Institute of Marine Research, P.O. Box 1870, Nordnes, 5817 Bergen, Norway; Swedish University of Agricultural Sciences, Department of Aquatic Resources, Drottningholm, Sweden

**Keywords:** age truncation, stock assessment, meta-analysis, demographic buffering, climate change, ecosystem-based management, matrix population model, life history trait, life table response experiment, hierarchical modeling

## Abstract

Large-scale commercial harvesting and climate-induced fluctuations in ocean properties shape the dynamics of marine populations as interdependent drivers at varied timescales. Selective removals of larger, older members of a population can distort its demographic structure, eroding resilience to fluctuations in habitat conditions and thus amplifying volatility in transient dynamics. Through the implementation of stricter management measures, many historically depleted fish stocks began showing signs of recovery in recent decades. But these interventions coincided with accelerated changes in the oceans triggered by increasingly warmer, more variable climates. Applying multilevel models to annual estimates of demographic metrics of 38 stocks comprising 11 species across seven ecoregions in the northeast Atlantic Ocean, this study explores how time-varying local and regional climates contributed to the transient dynamics of recovering populations exposed to variable fishing pressures moderated by management actions. Analyses reveal that progressive reductions in fishing pressure and shifting climate conditions nonlinearly shaped rebuilding patterns of the stocks through restorations of maternal demographic structure (reversing age truncation) and reproductive capacity. As the survival rate and demographic structure of reproductive fish improved, transient growth became less sensitive to variability in recruitment and juvenile survival and more to that in adult survival. As the biomass of reproductive fish rose, recruitment success also became increasingly regulated by density-dependent processes involving higher numbers of older fish. When reductions in fishing pressure were insufficient or delayed, however, stocks became further depleted, with more eroded demographic structures. Although warmer local climates in spawning seasons promoted recruitment success in some ecoregions, changing climates in recent decades began adversely affecting reproductive performances overall, amplifying sensitivities to recruitment variability. These shared patterns underscore the value of demographic transients in developing robust strategies for managing marine resources. Such strategies could form the foundation for effective applications of adaptive measures resilient to future environmental change.

## Introduction

Excessive human exploitation that persisted through the twentieth century depleted marine living resources worldwide, threatening biodiversity, food and nutrient security, and other ecosystem services that the oceans provide (Pauly, Watson & Alder 2005; Crowder *et al*. 2008; Worm *et al*. 2009). Long-lived, late-maturing vertebrate predators were disproportionately depleted by industrial fishing as target or incidental catch (Christensen *et al*. 2003; Juan-Jordá *et al*. 2022). As part of the efforts to halt and reverse the losses, national government and intergovernmental organizations began applying the precautionary principle to the management of marine fisheries (Hilborn *et al*. 2001; Punt 2006), following international agreements, including the Code of Conduct for Responsible Fisheries adopted by the Food and Agriculture Organization of the United Nations (FAO 1995; Melnychuk *et al*. 2021). Since the implementation of stricter management measures to regulate fishing operations in the early 1990s, an increasing number of historically overexploited fish stocks began showing signs of recovery (Hilborn *et al*. 2020; Juan-Jordá *et al*. 2022). These management interventions, however, coincided with accelerated changes in the oceans (Barange *et al*. 2018). Global climate is becoming increasingly warmer and more variable, modifying regional and local habitat conditions that support the biomass production of fish stocks, possibly shifting biologically sustainable levels of harvesting (Chassot *et al*. 2010; Free *et al*. 2019).

The dynamics of marine resource populations in many systems are shaped by drivers operating interactively at varied timescales, including high-frequency perturbations like fishing and low-frequency variability in ocean properties forced by climate (Rouyer *et al*. 2012; Goto *et al*. 2022b). When exposed to such persistent, nonstationary perturbations, near-term (transient) dynamics of an exploited population, which depend on both variable vital rates (survival, growth, and fecundity) and demographic (age and stage) structures, may deviate from asymptotic (equilibrium) dynamics (Hastings 2004; Koons *et al*. 2016). Past research instructs us that large-scale shifts in ecosystem productivity under alternative climate regimes can be propagated through the system to influence fluctuations in fisheries catches (Chassot *et al*. 2010; Cheung, Watson & Pauly 2013). Climate effects on marine species are often integrated over a series of oceanic, ecological, and biological processes over space and through time (Ottersen *et al*. 2001; Stenseth *et al*. 2003). Fluctuations in population structure and size can, for example, emerge with some time lag from age (or stage)-dependent demographic processes like birth and death (Jenouvrier *et al*. 2022) modulated by seasonal and interannual variations in weather and habitat conditions (Hutchings & Reynolds 2004; Lindegren *et al*. 2013). These transient fluctuations in the population could sustain over generations through delayed feedbacks (Hastings 2004).

In contrast, fishing operations and management actions that regulate the operations can directly modify short-term trends and variability in population structure and demographic rates, and thus the transient dynamics of exploited species (Gamelon, Sandercock & Sæther 2019). Large-scale commercial fishing often selectively removes larger, older members of the population, truncating age (and size) structure, modifying life history traits, reducing the capacity to absorb the effects of perturbations, and thus amplifying year-to-year variation in population size in variable environments (Hsieh *et al*. 2006; Rouyer *et al*. 2012). Fishing-induced age truncations (juvenescence) are widely reported for harvested marine populations (Hsieh *et al*. 2010; Rouyer *et al*. 2012), many of which suffer from reduced reproductive capacity (Hixon, Johnson & Sogard 2014). Climate-induced changes in the variability of ocean habitat conditions thus may indirectly promote or impede the recovery of depleted populations with weakened demographic resilience to perturbations (Lindegren *et al*. 2013; Harris *et al*. 2018).

In commercial fisheries a harvested population is commonly managed through regulation of fishing pressure based on its population status (Beddington, Agnew & Clark 2007). Because of high socioeconomic values, exploitation of marine resource populations is regularly monitored through scientific surveys and assessed based on population size estimates (Hilborn & Walters 1992). These population assessments form the scientific basis for the implementation of management measures like catch limits. When population size declines below a predefined threshold (a reference point set under the precautionary principle, ICES 2021; Goto *et al*. 2022a), for example, fishing pressure may be adjusted to mitigate the risk of overharvesting. This adaptive management cycle (human–natural system feedbacks through monitoring, assessment, and management action) thus can be considered as a natural experiment (Hilborn & Walters 1992; Jensen, Branch & Hilborn 2012), providing opportunities to gain insights into the transient dynamics of managed populations in variable environments. Understanding what drives these transients would help develop robust strategies to sustainably manage human exploitation of marine resource populations as we face increasingly uncertain conditions in the oceans.

This article reports a study that empirically explores how past fishing, management action, and climate variability influenced the transient dynamics of recovering fish populations in the northeast Atlantic Ocean (FAO Major Fishing Area 27, Fig. 1a). These populations experienced wide ranges of fishing pressure, climate, and management action in recent decades (Rouyer *et al*. 2011; Zimmermann & Werner 2019). Annual fishing pressures applied to many northeast Atlantic stocks began to be eased in the late 1990s to rebuild biomass that supports sustainable harvesting (Sparholt, Bertelsen & Lassen 2007; Lassen, Kelly & Sissenwine 2014). Following the reductions in fishing pressure, these intensively managed stocks experienced considerable changes in population size, demographic structure, and productivity (Hidalgo *et al*. 2012; Rouyer *et al*. 2012; Britten, Dowd & Worm 2016). During this period the regional climate also became increasingly warmer and more variable as greenhouse gas emissions rose, which was reflected especially in upper ocean heat content and transport (Delworth *et al*. 2016; Zanna *et al*. 2019). Warming patterns, however, highly varied among geographic subregions (ecoregions) owing to local air–sea feedbacks (Roemmich *et al*. 2015; Delworth *et al*. 2016), triggering a series of oceanic and ecological responses at variable rates (Beaugrand 2009; Delworth *et al*. 2016). To glean insights into the recovery patterns of historically depleted fish stocks, this study specifically asks how local and regional climate variability contributes to variations in the reproductive performance and transient growth of marine species exposed to variable fishing pressures moderated by management actions.

**Figure 1.**
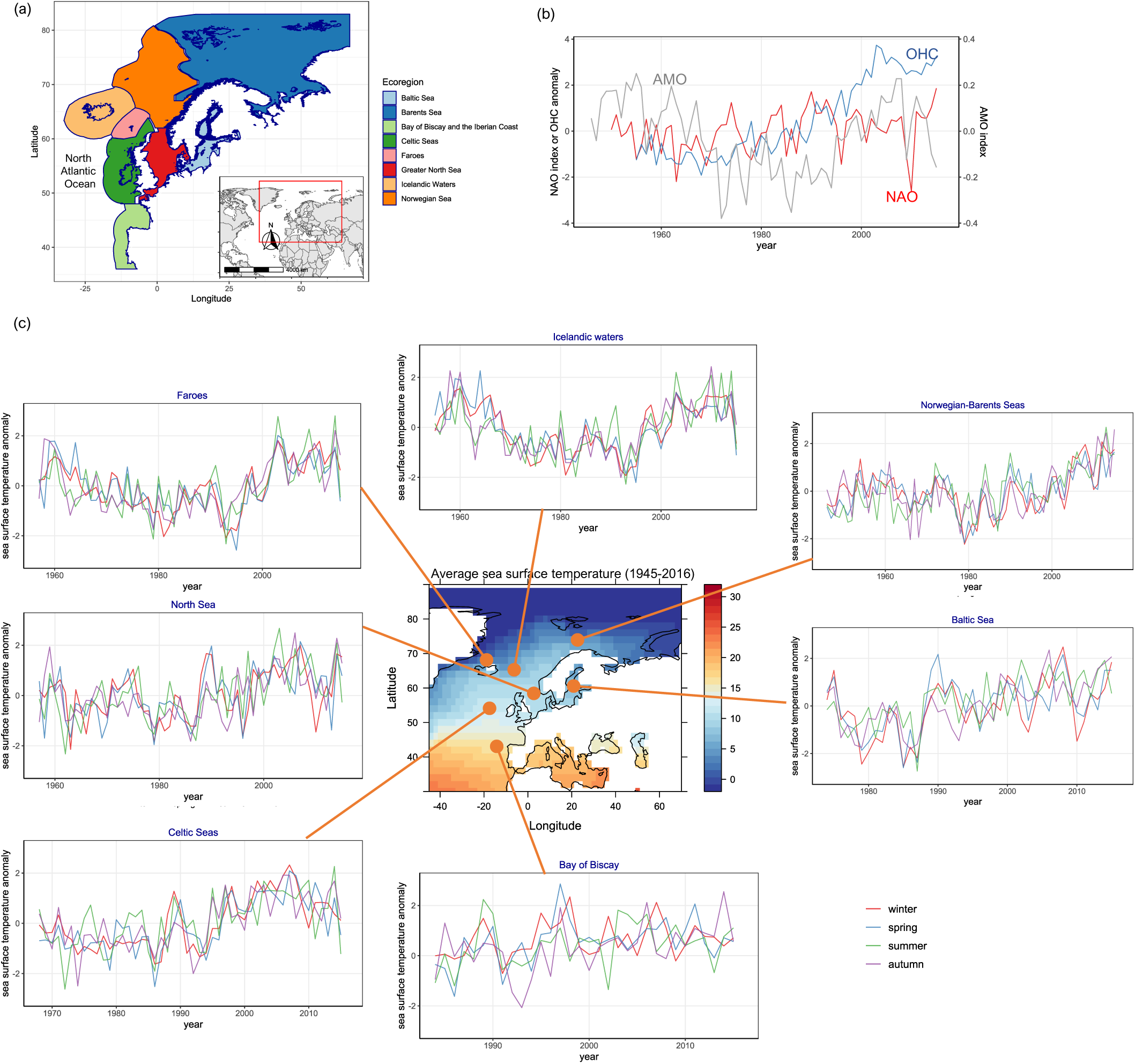
Study systems and climate conditions in the northeast Atlantic Ocean. (a) Geographic locations of the ecoregions, which are defined by the International Council of Exploration of the Sea (ICES): Norwegian and Barents Seas (62°0’N–82°0’N at 11°0’W–68°30’E), Greater North Sea (56°0’N at 2°55’E), Baltic Sea (53°35’N–65°54’N at 11°57’E–30°14’E), Faroes (60°0’N–63°0’N at 15°0’W–4°0’W), Icelandic waters (64°38’N at 18°55’W), the Bay of Biscay and the Iberian Coast (42°39’N at 8°53’W), and Celtic Seas (48°0’N–60°30’N at 18°0’W–1°21’W). (b) Regional climate indices (North Atlantic Oscillation (NAO) index, Atlantic Meridional Oscillation index, and upper (0–400m) ocean heat content (OHC) anomaly) during the study period (1946–2016). (c) Seasonal sea surface temperature anomalies in each ecoregion. The time series of temperature anomalies in each ecoregion are truncated to show only the years that the fish stock data and assessments included in the study are available.

## Materials and Methods

To investigate the transient dynamics of northeast Atlantic fish stocks at local and regional scales, a series of multilevel statistical models were applied to the time series of population and demographic metrics along with intrinsic and extrinsic covariates. In the following I describe (a) development of matrix population models to quantify transient population growth and elasticities to demographic rates through transient life table response experiments, (b) trend analyses to characterize the dynamics of reproductive fish biomass and age structure, transient growth, and fishing pressure, followed by analyses to test for direct fishing effects, and (c) evaluation of the relative contributions of time-varying intrinsic (reproductive fish biomass and age structure) and extrinsic (fishing pressure and climate) drivers to variations in recruitment success and in elasticity to recruitment success.

### Data

Time series (1946–2016) of annual estimates of age-specific information (mean body mass, abundance, maturity rate, and natural and fishing mortality rates) and management reference points (which define the management targets for fishing pressure and spawner biomass, as described below in *Management action for fisheries*) of marine fishes in the northeast Atlantic Ocean were extracted from the 2017 stock assessment reports by the International Council of Exploration of the Sea (ICES) (Table 1). In the assessment of each stock (defined here as a population in a management unit area defined by the ICES) the demographic rates and life history traits were estimated using data from commercial catch reporting and scientific monitoring surveys. Stock sizes and fishing mortality rates then were estimated annually using age-structured population models fitted to time series data of abundance indices (ICES 2017e; ICES 2017a; ICES 2017b; ICES 2017c; ICES 2017f; ICES 2017d). The reliability of the models and data had been evaluated by experts prior to assessments and extensively documented (e.g., ICES 2016). For analysis I excluded stocks with missing age-specific information, rejected assessments, or shorter time series (less than 15 years). The final dataset consists of 38 stocks comprising 11 species (cod *Gadus morhua*, haddock *Melanogrammus aeglefinus*, saithe *Pollachius virens*, herring *Clupea harengus*, plaice *Pleuronectes platessa*, turbot *Scophthalmus maximus*, sole *Solea solea*, whiting *Merlangius merlangus*, sprat *Sprattus sprattus*, megrim *Lepidorhombus spp*., and four-spot megrim *Lepidorhombus boscii*) across seven ecoregions in the northeast Atlantic Ocean defined by the ICES: the Norwegian and Barents Seas (combined as a single ecoregion for analysis), the Greater North Sea (hereafter the North Sea), the Baltic Sea, Faroes, Icelandic waters, the Bay of Biscay and the Iberian Coast (hereafter the Bay of Biscay), and the Celtic Seas (Table 1 and Fig. 1a). These stocks support some of the largest fisheries in the region, collectively yielding 1.9 to 2.6 million tonnes of commercial landings annually in the recent decades (1997–2016).

**Table 1.**
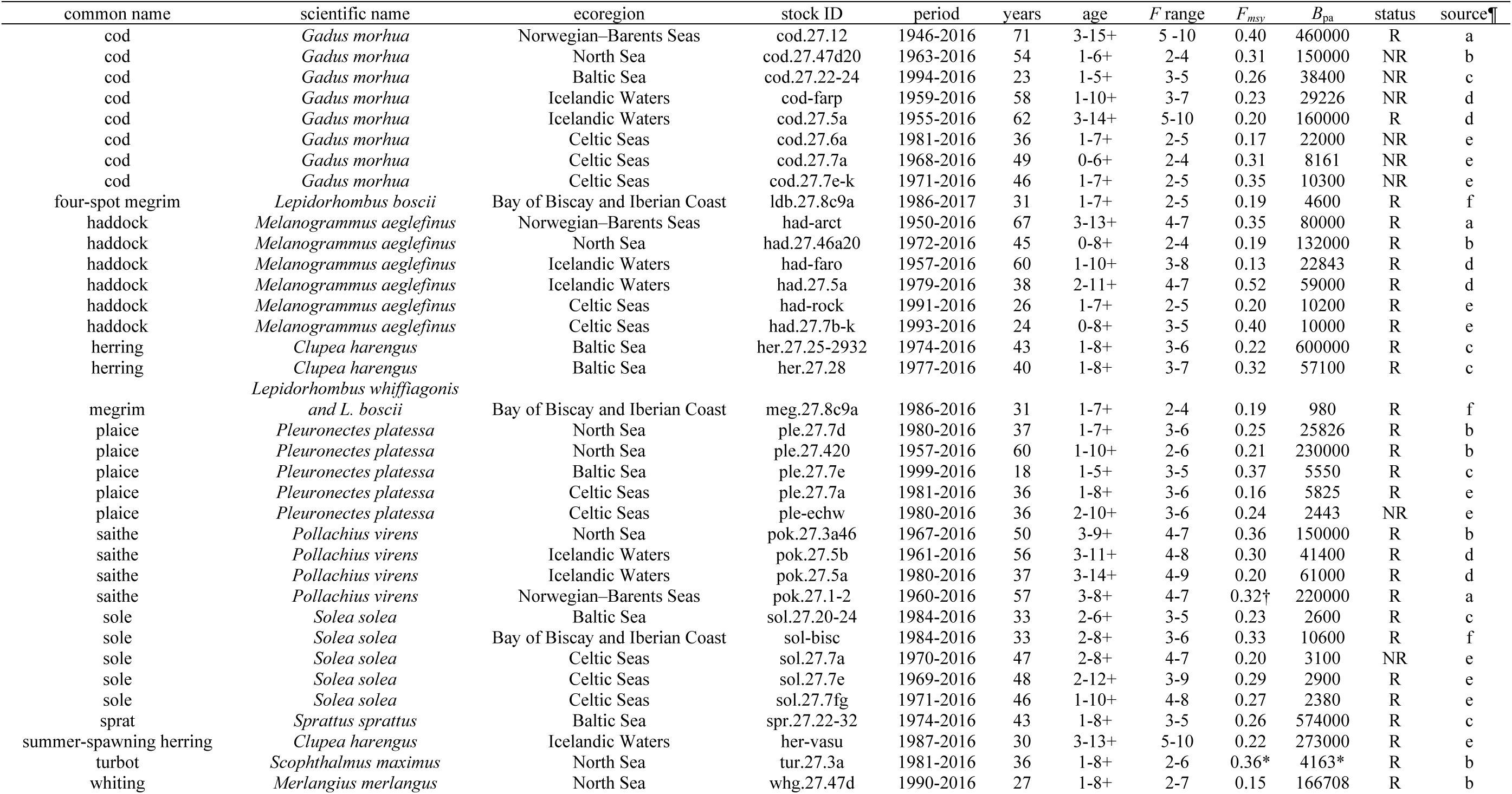
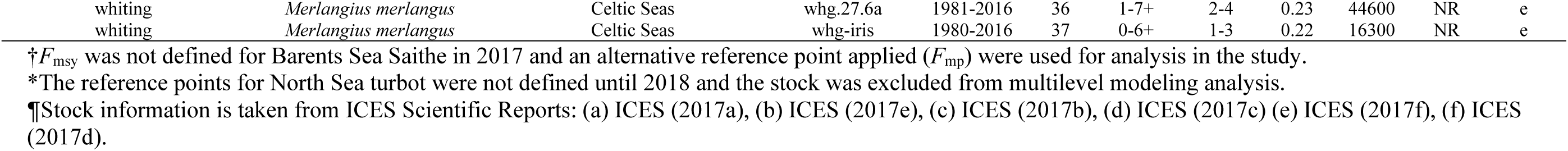
Stock-specific information and management measures used in the assessments of 38 fish stocks in the northeast Atlantic Ocean included in the study. Stock ID indicates a unique ID for each stock assigned by the International Council of Exploration of the Sea (ICES); period and years indicate the years and the total number of years that stock data are available; age indicates the age classes included in the assessments; *F* range indicates the age range used to compute mean fishing mortality rates for assessment; *F*_msy_ and *B*_pa_ indicate the management reference points set for each stock as target fishing mortality rates that produce maximum sustainable yields and biomasses that satisfy the precautionary approach; and status indicates recovery status assigned based on spawner biomass relative to *B*_pa_ for analysis (R: recovering and NR: non-recovering) in this study.

Climate indices and sea surface temperature (SST) anomalies were used as proxies that represent variations in overall regional (North Atlantic) and local (ecoregion) climate conditions (1946–2016), which are filtered through oceanic processes fish populations directly and indirectly experience over interannual and decadal timescales (Ottersen & Stenseth 2001; Stenseth *et al*. 2003). SST was chosen as a proxy for local habitat conditions because its reconstructed data can provide adequate spatial and temporal coverage for this study. Further, the climate data were used as covariates in multilevel models with early life-history processes as response variables (as described below in *Covariates of recruitment success* and *Covariates of elasticity to recruitment success*). Climate effects on these processes can, for example, be mediated by lower trophic level (like zooplankton) production responding to regional climate-driven SST anomalies (Beaugrand & Kirby 2018) and other ocean properties (like heat transport), which may in turn manifest as delayed effects in young fish survival (Hjermann *et al*. 2007).

Monthly time series of three regional climate indices were downloaded and averaged over winter months (when the atmosphere is most active in the region, Stenseth *et al*. 2003) for analysis (Fig. 1b): the North Atlantic Oscillation (NAO) and Atlantic Multidecadal Oscillation (AMO) indices (both computed from the Hadley Centre Sea Ice and Sea Surface Temperature data set [HadISST1, https://www.metoffice.gov.uk] and the extended reconstructed SST [ERSSTv5, NOAA Physical Sciences Laboratory; https://psl.noaa.gov] and averaged over the sources), and upper (0–700 m) ocean heat content (OHC) anomaly in the North Atlantic (NOAA National Centers for Environmental Information; https://www.ncei.noaa.gov). The NAO index reflects the dominant mode of atmospheric variability between the Azores and Iceland, linked with a range of large-scale oceanic features (such as surface temperature and heat content) in the North Atlantic that regulate biological and ecosystem processes (Stenseth *et al*. 2003). The AMO index, computed by averaging SSTs over the North Atlantic (25°–60°N at 7°–70°W) using the method by van Oldenborgh *et al*. (2009), also reflects large-scale climate variability that may influence the movement and migration of marine species (Faillettaz *et al*. 2019). OHC measures the amount of excess heat (from increasing greenhouse gases) trapped in the ocean, integrating basin-wide heat storage and transport (Zanna *et al*. 2019). Because of high signal-to-noise ratio, OHC reflects more consistent decadal trends (less influenced by natural variability) than other climate indices (Roemmich *et al*. 2015).

Reconstructed gridded monthly SST data were downloaded from and averaged over three sources to account for systematic uncertainties in each source: HadISST1, COBE Sea Surface Temperature SST (NOAA Physical Sciences Laboratory), and ERSSTv5. I first averaged the SST data over ecoregion and for each ecoregion over season; winter (December in the previous year–February, wSST_anom_), spring (March–May, spSST_anom_), summer (June–July, smSST_anom_), and autumn (September–November, aSST_anom_). I then computed SST anomalies by subtracting the mean across years (1946–2016) for each season and ecoregion (Fig. 1c).

### Transient population growth and elasticity

I used transient population growth rate as an integrated measure that captures time-varying demographic rates and age structures responding to non-stationary extrinsic factors (fishing pressure, management action, and climate) and estimated the growth rates for each stock using an age-structured matrix projection model (Caswell 2001). Leslie (state-transition) matrices (*A*) were first constructed with annual estimates of age-specific survival and reproductive rates.

Survival rates (*S*_a,y_) were estimated as *S*_a,y_ = exp(–*Z*_a,y_), where *Z*_a,y_ = *F*_a,y_ + *M*_a,y_, and *F*_a,y_ and *M*_a,y_ are fishing and natural (non-fishing like predation and starvation) mortality rates for *a*-year olds in year *y*. Egg production rates are often not estimated in the assessments of marine fish stocks, and here I estimated recruitment success (integrating over reproduction and early life-history stage survival) rate per spawner (reproductive fish) for *a*-year olds in year *y* (RS_a,y_) (Rouyer *et al*. 2011) as a proxy for per capita fecundity rate as

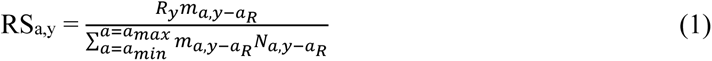

where *R*_y_ is the recruit number in year *y*, *m*_a,y-a_R and *N*_a,y-a_R are the maturity rate (fraction of juveniles becoming adults) and abundance of *a*-year olds in year *y*-*a*_R_ (spawning year), *a*_R_ is the age of recruits, which accounts for variations in age at recruitment (0–3 years old) among stocks reported in the assessments. *a*_min_ and *a*_max_ are the youngest and oldest ages in the stock. I then numerically estimated deterministic transient growth rates (*λ*_trans_) by projecting age-specific abundances using the annual state-transition matrices and the annual estimates of age-specific numbers taken from the assessments as

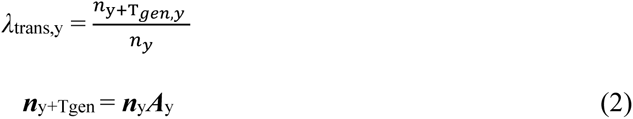

where ***n***_y_ and ***A***_y_ are a vector of initial age-specific numbers and an *a* × *a* state-transition matrix in year *y*, and *T*_gen_ is the generation time (years) in year *y*. To account for variations in life-history strategy among stocks, the transient growth rates of each stock were estimated over its generation time, which is defined as the mean age of parents of a cohort (Caswell 2001),

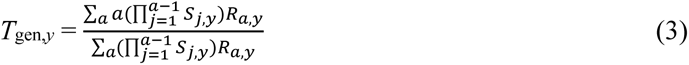

The transient growth rates here were thus defined in timescales of 2.6 to 6.7 years (Fig. S1), relevant for fisheries management.

To evaluate the contributions of demographic variations to transient growth I performed transient life table response experiments (LTRE) by numerically estimating transient elasticities (*e_y_*) to survival and fecundity rates through matrix projections (Koons *et al*. 2016); demographic parameters (fecundity or survival) of all ages were first perturbed by a small amount (1%), and stock numbers were projected to re-compute *λ*_trans,*y*_ (Durant *et al*. 2013). Elasticity (relative sensitivity), defined as the proportional change in *λ*_trans_ relative to a proportional change in a demographic rate (Caswell 2001), was computed as 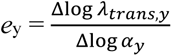, where *α*_y_ is the survival or fecundity rates in year *y* that are perturbed (Coulson *et al*. 2004). Age-specific elasticities were then summed for analysis; changes in elasticities to juvenile and adult survival rates were evaluated separately by summing age-specific elasticities weighted by maturity rates. To quantify how the transient dynamics deviate from expected long-term equilibriums (asymptotic dynamics assuming stable environments and age structures) across years I also computed year-specific asymptotic (geometric) growth rates and stable age distributions from the logarithms of the dominant (largest) eigenvalues and the right eigenvectors (respectively) of the annual state-transition matrices (Caswell 2001).

To evaluate how variations in demographic structure have contributed to the reproductive performance and recovery patterns of fish stocks, I used mean age and age diversity of spawners as indices for demographic structure. Mean age (SA_mean_) is the weighted (by abundance) average of spawner age as

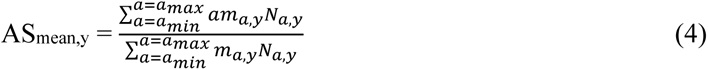

The Shannon diversity index was used as an indicator of age diversity (SA_diversity_) and computed 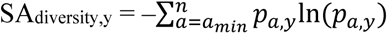, where *p*_a,y_ is the proportion of *a*-year-old spawner number in a given age group in year *y* (Marteinsdottir & Thorarinsson 1998).

### Management action for fisheries

Management action (here adjustment of fishing pressure) is based on perceived current stock status and harvest control rule set by the ICES (which are designed to achieve sustainable harvesting without compromising the reproductive capacity of the stock). The current status is assessed using scientific monitoring and catch data before the provision of annual catch advice (total allowable catch). Under the control rule, when the estimated SB remains above a fixed threshold (*B*_pa_) at the start of the year following the assessment, the catch limit is computed based on target exploitation rate (*F*_msy_). These two parameters (*B*_pa_ and *F*_msy_) are designed to prevent overharvesting by accounting for uncertainty in population and harvest dynamics (ICES 2019a). When the SB falls below *B*_pa_, exploitation rate is adjusted to *F*_msy_ scaled to the proportion of SB relative to *B*_pa_, thereby allowing the population to rebuild (adaptive management). For analysis here fishing pressure and SB relative to stock-specific reference points (*F̄*/*F*_msy_ and SB/*B*_pa_) were used as proxies to evaluate the influence of the management measures on the demographic metrics of fish stocks. Fishing pressure is defined as the instantaneous rate of fishing mortality (*F̄*), which is commonly estimated by averaging age-specific fishing mortality rates over dominant ages in fishery catch (Table 1).

### Data analysis

To explore how management-guided changes in fishing pressure and climate change influenced the vital rates, transient dynamics, and recovery patterns of the northeast Atlantic fish stocks, I applied multilevel (hierarchical) modeling to derived demographic metrics to account for nonindependence in data structure (Gelman & Hill 2006) with stock, ecoregion, and year as grouping factors (random intercepts). Data exploration including identification of collinearity (through visual inspection correlation plots and variance inflation factors) and outliers (>three standard deviations) was performed before analysis (Zuur *et al*. 2009). The data exploration revealed two groups of stocks with diverging (increasing and declining) patterns in stock status based on relative spawner biomass (SB/*B*_pa_). To explore correlations underlying the differences I applied generalized additive mixed effects models (GAMMs, Wood 2017) to these groups separately. In the following analyses the stocks were grouped based on stock status in the last 15 years of the time series (2002–2016); if spawner biomass is larger than 80% of a biological threshold (*B*_pa_, Costello *et al*. 2016) for at least 50% of the period (eight years) a stock is classified as ‘recovering’, otherwise ‘non-recovering’ (sensitivity analyses showed that the diverging patterns in SB/*B*_pa_ are robust to variations in the criteria used, Fig. S2 and S3). This classification resulted in 27 recovering and 10 non-recovering stocks (Table 1).

*Trends in populations, demographics, and fishing*. Trend analyses were first performed with year as a covariate to capture shared long-term historical patterns in relative spawner biomass, spawner demographic structure (mean age and age diversity), relative fishing pressure (*F̄*/*F*_msy_), and transient growth. The response variables were assumed to follow a Gamma distribution with continuous, positive expected values *μ* and a logarithmic link function as

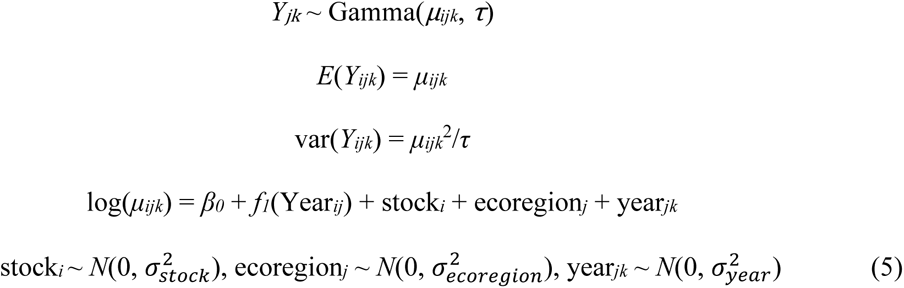

where *Y_jik_* is the response of stock *i* in ecoregion *j* in year *k*, *τ* is a Gamma distributed parameter, *β*_0_ is the intercept, and *ꬵ*_1_ is a smoother function (thin-plate cubic splines with five maximum degrees of freedom or knots) for year; *stock_i_*, *ecoregion_j_*, and *year_jk_* are normally distributed random intercepts with mean zero and standard deviation *σ* for stock, ecoregion, and year nested within ecoregion. The structure of random effects was selected based on likelihood ratio tests applied to ten structures fitted with a restricted maximum likelihood method (REML).

Abundance and demographic parameters in stock assessments are estimated with several known sources of uncertainty (including observation, process, and estimation errors, ICES 2019b), which may bias post-hoc analyses when used as data (Brooks & Deroba 2015). To address this issue, I performed sensitivity analyses by Monte Carlo simulations to illustrate how uncertainties in the estimates can propagate through matrix population and multilevel modeling analyses.

Because uncertainty in age-specific parameters was not consistently reported in the assessments, I first generated 1000 simulated datasets with Gaussian noise in recruit numbers and fishing mortality rates using the stock- and year-specific coefficients of variation derived from the assessments and propagated the estimation uncertainties through the computation of state-transition and transient population growth rates (see Fig. S4 for more detail). This step was then followed by multilevel model-fitting with transient population growth as a response variable and year as a covariate (as described above) applied to each simulated dataset.

Next, I assessed direct effects of fishing on spawner biomass and age structure by evaluating relationships between relative fishing pressure and relative biomass or age structure of spawners as

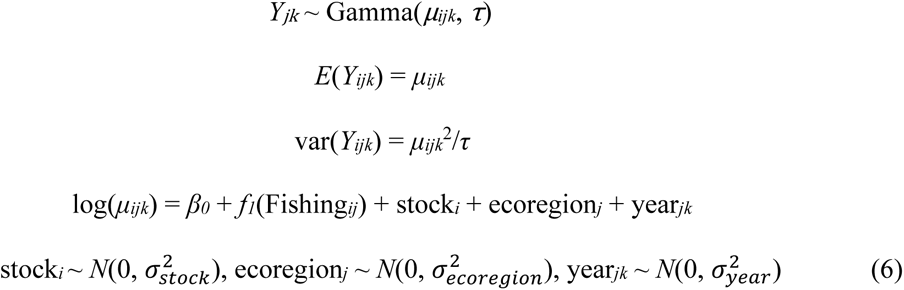

where *Y_jik_*,*τ*, *β*_0_, *σ_stock_*, *σ_ecoregion_* and *σ_year_* are the same as above. *ꬵ*_1_ is a smoother function for relative fishing pressure.

The contributions of fishing, management action, and climate to the transient dynamics of fish stocks with variable age structures were evaluated in two steps. I first tested for time-varying contributions of these extrinsic factors to variations in vital rates and then evaluated how these factors influenced the elasticities of transient population growth:

*Covariates of recruitment success*. Because natural mortality rates were assumed time-invariant in most of the assessments, variations in survival rates are in effect driven by fishing. Here, RS was thus selected as a demographic response variable (which integrates probabilities of spawning and early life history stage survival) for analysis. Because RS can vary with variations in spawner age structure and biomass (intrinsic factors), these intrinsic factors were also tested. To test for the time-varying contributions of the extrinsic and intrinsic factors, the following model was fitted to the data:

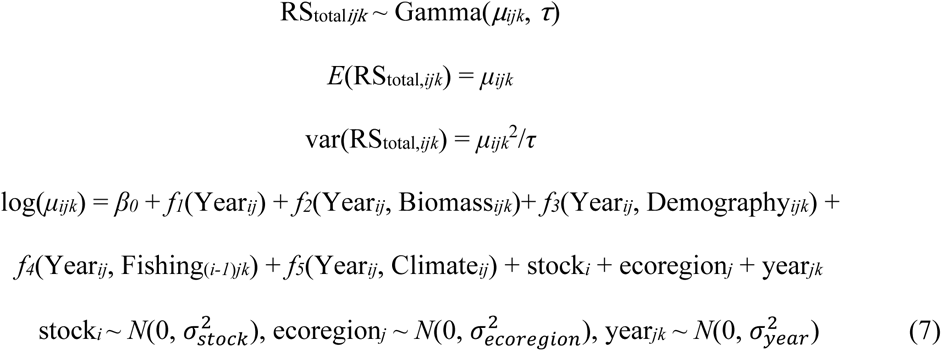

where RS_total,*ijk*_ is the total per capita recruitment success rate of stock *i* in ecoregion *j* in year *k*, *β*_0_ is the intercept, *ꬵ*_1_–*ꬵ*_5_ are smoother functions for covariates: year, year × density, year × demographics (spawner mean age and age diversity), year × climate, and year × fishing. *Climate*, *Biomass*, and *Demography* were lagged 0–3 years (depending on stock) to align with climate conditions and spawner status (biomass and age structure) in spawning year, whereas *Fishing* was lagged an additional year (spawning year–1) to reflect indirect effects of fishing on recruitment success that are not accounted for by variations in spawners. Regional climate covariates were also lagged an additional year (*Climate*_(_*i-1*_)_*j*) to reflect indirect effects through regulation of local climate and ecosystem processes. Time (year)-varying covariate smoothers were tested as tensor product construction (with 5 × 2 maximum knots using the *t2* function in the R package *mgcv*). For interpretability and model convergence one covariate of each type was tested at a time; one local or regional climate index (see above) for climate, SB or SB/*B*_pa_ for spawner density, *F̄* or *F̄*/*F*_msy_ for fishing pressure, and SA_mean_ or SA_diversity_ for demographics.

*Covariates of elasticity to recruitment success*. The transient dynamics of depleted fish stocks with age truncation often become recruitment-dependent and thus sensitive to variations in climate and habitat conditions. As the stocks recover (increases in abundance and age diversity), the transient dynamics are expected to become less sensitive to recruitment variability. To test for the time-varying contributions of the extrinsic and intrinsic factors to variations in elasticity to RS (*e*_rec_), the following model was fitted to the data, assuming *e*_rec_ follows a Beta distribution with expected values *μ* (0 < *μ* <1) and a logit link function as

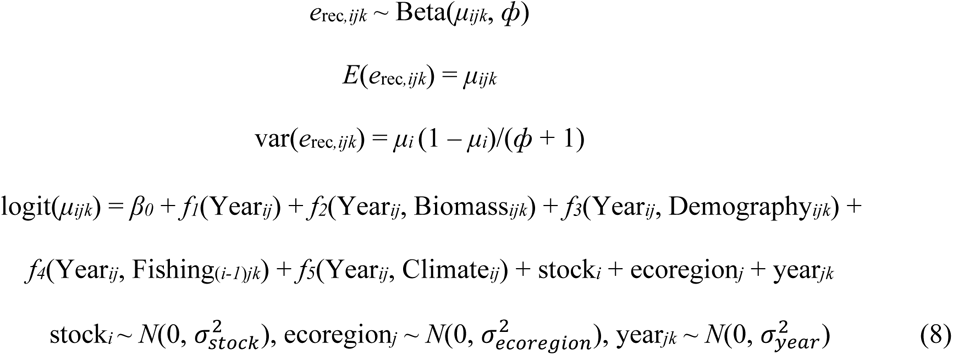

where *ф* is a parameter (> 0), and *β*_0_, *ꬵ*_1_–*ꬵ*_5_, *σ_stock_*, *σ_ecoregion_* and *σ_year_*, are the same as above (here tensor product construction for time-varying covariate smoothers was modeled using the *te* function in the R package *mgcv*).

All numeric covariates were standardized by subtracting the mean and dividing by the standard deviation (*z*-scores). In model selection intrinsic covariates were first added, followed by indirect fishing, and then climate. With the random effect structure selected (as described above, Table S1) the models were tested sequentially for covariates by fitting with a maximum likelihood method, followed by parameter (re)estimation with REML (Zuur *et al*. 2009) for the model with the lowest Akaike information criterion (AIC) score and highest model weight (*w*). AIC is computed as AIC = –2ln(*L*) + 2*k*, where *L* is the likelihood of the model and *k* is the number of parameters, and the model weight of model *i* (*w_i_*) is computed as

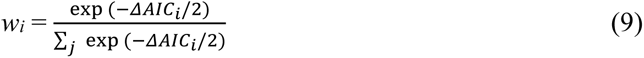

As diagnostics, model residuals were checked for homogeneity and regressed against covariates that were both included and not included in the selected models (Zuur *et al*. 2009). All analyses were performed in R (version 4.2.1, R Development Core Team 2022) with the R packages *gamm4* (v.0.2-6, Wood & Scheipl 2020) and *mgcv* (v.1.8-40, Wood 2022) for model-fitting, *mgcv* and *gratia* (v.0.7.3, Simpson & Singmann 2022) for model diagnostics, and *popbio* (v.2.7, Stubben, Milligan & Nantel 2020) and *popReconstruct* (1.0-7, Wheldon 2019) for computations of demographic parameters and projections of population dynamics.

Because fish populations in different ecoregions experience a wide range of climate conditions and often have adapted to local environments, fish–covariate patterns may deviate from shared patterns captured by the regional models. To test this hypothesis, ecoregion-scale analyses were also performed using the same model structures for RS_total_ and *e*_rec_ and the model selection steps as above except for random effects. Because random effect estimates become less stable when the number of groups is fewer than eight (Bolker 2015), the random effect of the ecoregion-scale models consisted of year only, with stock being a fixed effect, except for the Celtic Seas; the random effects of the Celtic Seas model consisted of year and stock.

## Results

Analyses were performed to address primarily (a) whether long-term changes in spawners (biomass and age structure) shared by intensely managed fish stocks in the northeast Atlantic reflect management-guided changes in fishing pressure, (b) how intrinsic (spawner biomass and age structure) and extrinsic (fishing and climate) drivers contribute to variations in recruitment success, and (c) how the intrinsic and extrinsic drivers influence the sensitivity of transient dynamics to recruitment variability. Overall, the diverging patterns in spawner biomass between the recovering and the non-recovering stocks also emerged in age structure and transient population dynamics, reflecting differential reductions in fishing pressures the stocks experienced. The differences in spawner biomass and age structure between the groups were further reflected in variations in recruitment success and elasticity to recruitment success, indicating divergence in the strength of density dependency and the contribution of older spawners. Although climate effects varied among ecoregions, most showed shifting responses in recruitment (from positive to negative) and in elasticity to recruitment (from negative to positive) to increasingly warmer than average local and regional climates.

Regionwide analyses showed that relative fishing pressures on the recovering stocks on average peaked in the early 1990s (*F̄*/*F*_msy_ = ∼2.2) and near-linearly declined since (∼2.1% per year; ΔAIC = 88.4), followed by increasing mean age (ΔAIC = 35.7), relative spawner biomass (ΔAIC = 17.2), and age diversity (ΔAIC = 21.1) (Fig. 2a). In contrast, relative fishing pressures on the non-recovering stocks kept rinsing until the late 1990s (to *F̄*/*F*_msy_ = ∼3.7 on average) before declining (∼3.0% per year, ΔAIC = 63.3), but their relative spawner biomass continued to decline (ΔAIC = 92.7 and 31.5; Fig. 2a). Although the mean spawner age of the non-recovering stocks began rising with declining relative fishing pressures, no change in age diversity was detected (Fig. 2a). Direct effects of selective fishing consistently emerged as negative correlations between relative fishing pressure and spawner biomass and age structure in both stock groups (ΔAIC **=** 15.1–165.0, Fig. 2b). Contrasting temporal patterns between relative fishing pressure and spawner features were more pronounced in some ecoregions captured in year-specific (nested within ecoregion) random effect (variance) estimates, especially the Norwegian–Barents Seas, which showed abrupt shifts in the early 1990s (Fig. S5).

**Figure 2.**
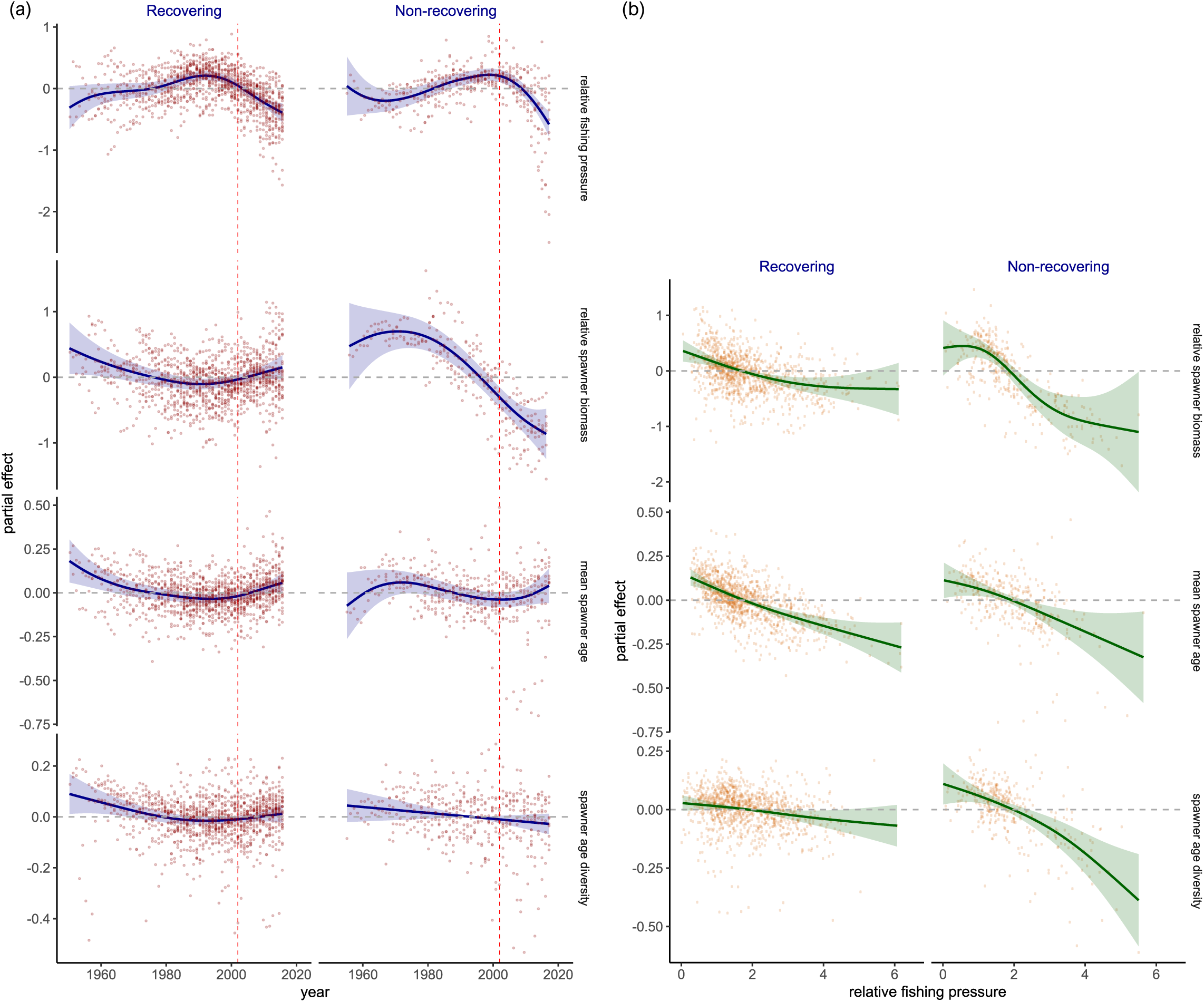
Regional patterns in fishing pressure and reproductive metrics of the recovering and non-recovering northeast Atlantic fish stocks during 1946–2016. (a) Temporal trends in relative fishing pressure, relative spawner biomass, mean spawner age, and spawner age diversity. Vertical dashed lines indicate the reference year (2002) used to define the recovery status of fish stocks based on relative spawner biomass in analysis. (b) Relationships between relative fishing pressure and relative spawner biomass and age structure metrics. Solid lines and ribbons indicate regional average trends in time (a) or relative fishing pressure (b) and 95% confidence intervals estimated by generalized additive mixed effects modeling with year (a) or relative fishing pressure (b) as a fixed effect, and ecoregion, year within ecoregion, and stock as random effects. Circles indicate partial residuals estimated from the models. Relative fishing pressure and relative spawner biomass are defined as estimated fishing mortality rate and spawner biomass relative to management reference points (*F*_msy_ and *B*_pa_) defined by the International Council of Exploration of the Sea (ICES).

The time series of transient population growth rates reflected the contrasting patterns between the recovering and the non-recovering stocks, and variations among ecoregions (Fig. 3 and S6). Sensitivity analyses showed that multilevel model parameter estimates in the trend analyses for both stock groups were robust to uncertainties in select abundance and demographic rates used as input (simulated parameter values < 2 standard deviations of the estimates, Fig. S7). The regional average growth rates of the recovering stocks linearly increased over the study period (ΔAIC = 15.0, Fig. 3). Although the relative spawner biomass of the non-recovering stocks continued to decline, their average growth rates began rising in the late 1990s (ΔAIC = 2.8, Fig. 3) when the relative fishing pressures began declining (Fig. 2a). These transient dynamics differed from asymptotic dynamics; the transient growth rates were lower than asymptotic growth rates in most stocks, and the deviations varied over years, with ∼50% (19) of the stocks showing increasingly less differences in the last decade (Fig. S8). The observed age distributions of many of these stocks also differed from theoretical stable age distributions and skewed toward younger ages (Fig. S9) owing to disproportionately higher annual mortality rates of older fish by fishing (Fig. S10).

**Figure 3.**
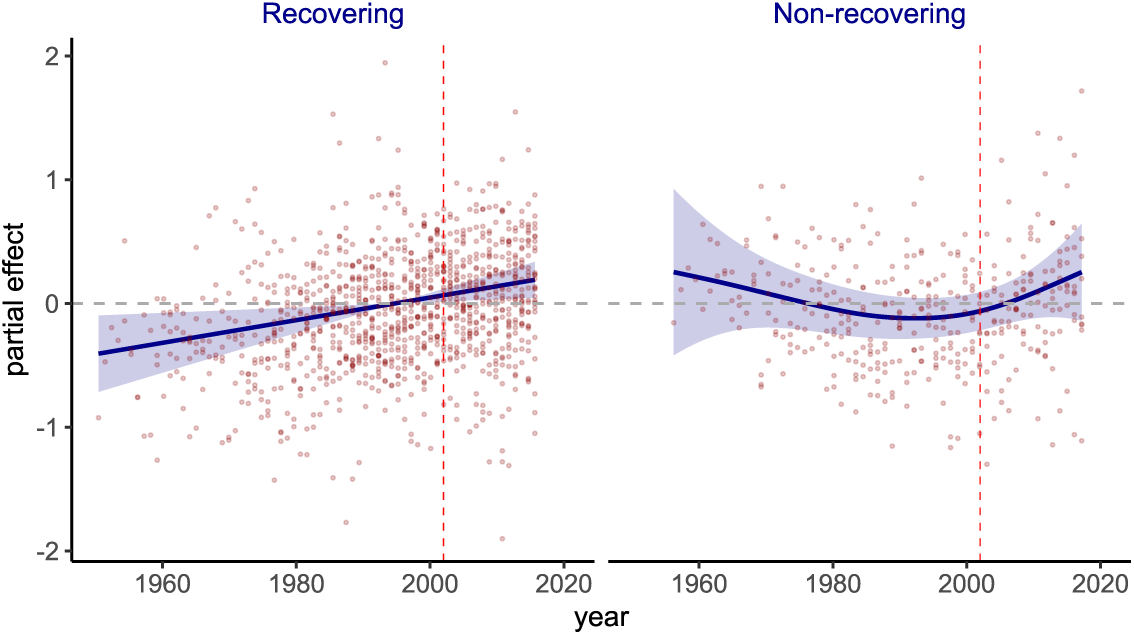
Regional trends in transient population growths of the recovering and non-recovering northeast Atlantic fish stocks during 1946–2016. Solid lines and ribbons indicate temporal trends and 95% confidence intervals estimated by generalized additive mixed effects modeling with year as a fixed effect, and ecoregion, year within ecoregion, and stock as random effects. Circles indicate partial residuals estimated from the models. Vertical dashed lines indicate the reference year (2002) used to define the recovery status of fish stocks based on relative spawner biomass in analysis.

Regionwide analyses detected declining trends in per capita recruitment success rate in both recovering and non-recovering stocks (ΔAIC = 2.1 and 2.7, Fig. 4a), with high among-stock variances (*σ*^2^*_stock_*= 0.31 and 0.48, Fig. 4f-h). Ecoregion-specific random intercepts also revealed generally higher year-to-year fluctuations in the non-recovering than in the recovering stocks (Fig. S6). After accounting for variations among stocks and ecoregions (and years within each ecoregion), the recruitment rates of both stock groups declined with increasing relative spawner biomass (Fig. 4c and Table 2), which was more supported by data than absolute biomass (Table S2). But the temporal patterns of density dependency differed between the groups; the density dependent effect in the recovering stocks increased as the biomass rose, whereas that in the non-recovering stocks emerged in the last two decades but declined over time (Fig. 4c). Likewise, the contribution of spawner age structure to recruitment variability differed between the groups; the relationship between recruitment success and mean age in the recovering stocks shifted from positive to negative in the early 1990s, whereas that in the non-recovering stocks remained largely positive (Fig. 4b). An indirect positive fishing effect on recruitment rate was detected in the recovering stocks until the late 1980s but gradually diminished since (Fig. 4d). The models for both stock groups showed weak support for climate effects (Table S2). Of the climate indices tested for the non-recovering stocks (whose model had slightly more support than the recovering stocks), the model with ocean heat content anomaly in the year of spawning showed shifting (positive to negative) climate effects in the recent decades (Fig. 4e).

**Figure 4.**
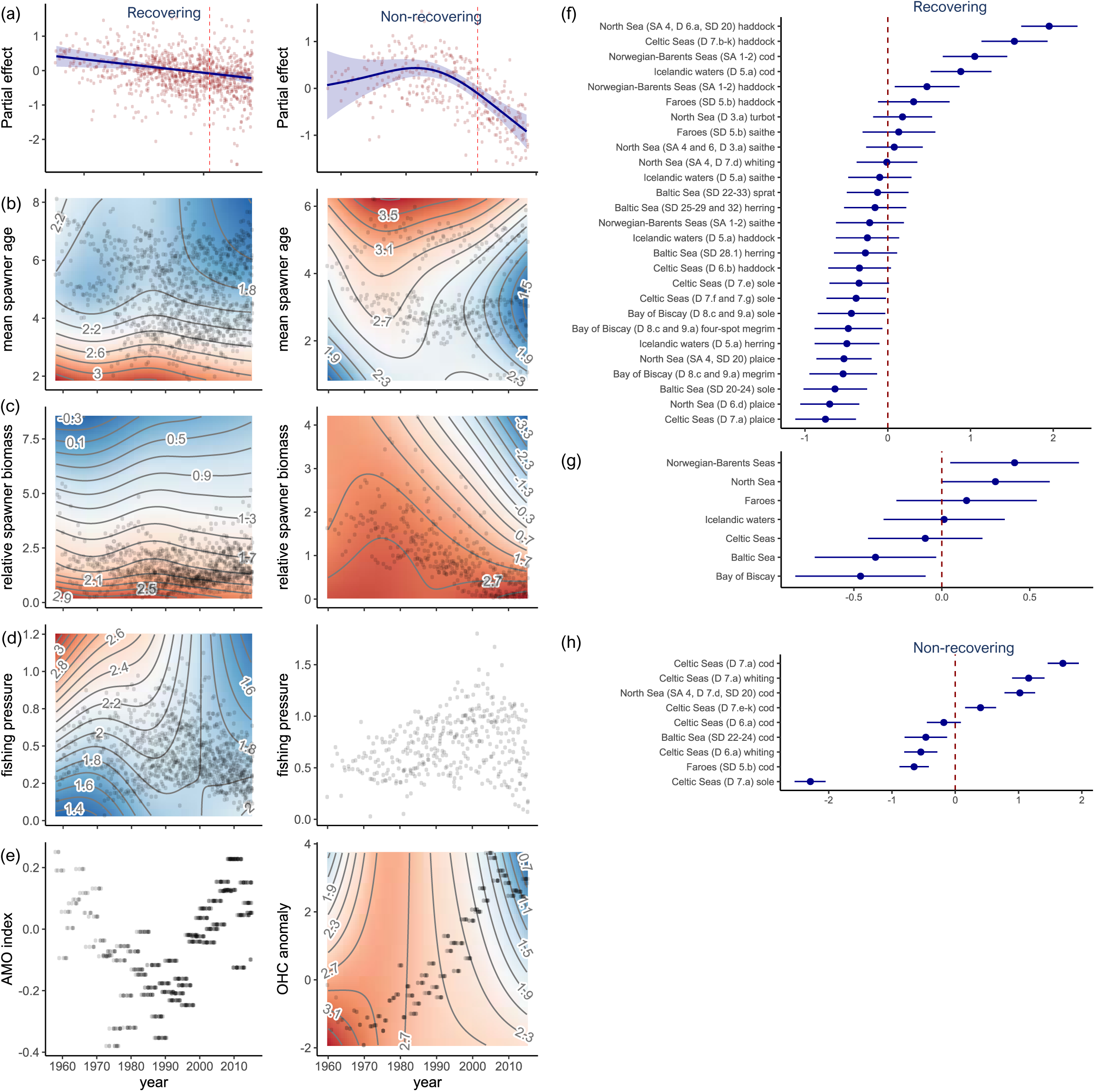
Regional trends in and covariates of per capita recruitment success (fecundity) of the recovering and non-recovering northeast Atlantic fish stocks during 1946–2016. (a) Temporal trends. Solid lines and ribbons indicate temporal trends and 95% confidence intervals estimated by generalized additive mixed effects modeling with year as a fixed effect, and ecoregion, year within ecoregion, and stock as random effects. Red circles indicate partial residuals estimated by the models. (b–e) Time-varying effects of spawner age structure, spawner biomass, fishing pressure. and climate condition. Covariates selected for each type in the models are indicated in the y-axis labels. Color gradients and contour lines indicate the partial effects of covariates estimated (as tensor interaction terms) in the models (only the covariate types that are supported by data are shown). Warmer colors indicate higher values, and the ranges of the partial effects differ among the covariates and are indicated as values on the contour lines. Gray circles indicate back-transformed input data. (f–h) Random effects estimates (±95% confidence intervals) for stock and ecoregion for the recovering and non-recovering stocks (ecoregion-specific estimates for the non-recovering stocks are negligible; random effect estimates for year nested within ecoregion are given in Fig. S6b).

**Table 2.**
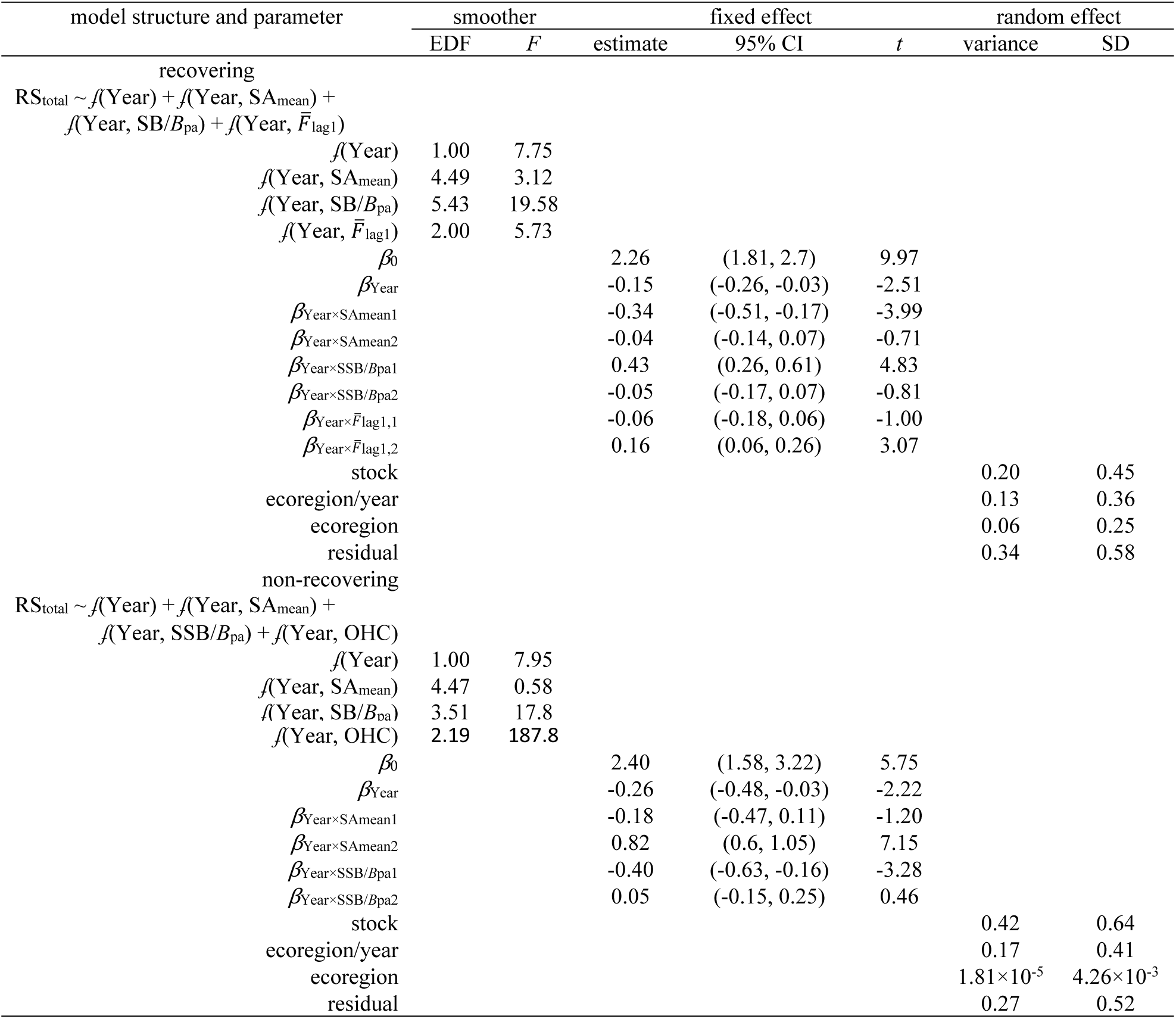
Parameter estimates and test statistics for the generalized additive mixed effects models selected for the northeast Atlantic fish stocks with per capita recruitment success (RS_total_) as a response variable (Table S2). EDF, *F*, CI, *t*, and SD indicate the effective degrees of freedom, F- statistic, confidence intervals, t-value, and standard deviation. All time (year)-dependent smoother terms for covariates are modeled as tensor product construction (using the *t2* function in the R package *mgcv*).

**Table 3.** Parameter estimates and test statistics for the generalized additive mixed effects models selected for the northeast Atlantic fish stocks with elasticity to recruitment success (*e*_RS_) as a response variable (Table S2). EDF, *F*, 95% CI, *t*, and SD indicate the effective degrees of freedom, F-statistic, confidence intervals, t-value, and standard deviation. All time (year)- dependent smoother terms for covariates are modeled as tensor product construction (using the *te* function in the R package *mgcv*).

| model structure and parameter | smoother |  | fixed effect |  |  | random effect |  |
| --- | --- | --- | --- | --- | --- | --- | --- |
|  | EDF | *F* | estimate | 95% CI | *t* | variance | SD |
| recovering |  |  |  |  |  |  |  |
| $e_{RS} \sim f(\text{Year}) + f(\text{Year}, SA_{\text{mean}}) + f(\text{Year}, SB/B_{\text{pa}})$ | | | | | | | |
| $+ f(\text{Year}, \bar{F}_{\text{lag1}}/F_{\text{msy}}) + f(\text{Year}, AMO_{\text{lag1}})$ | | | | | | | |
| $f(\text{Year})$ | 1.00 | 0.60 | | | | | |
| $f(\text{Year}, SA_{\text{mean}})$ | 3.51 | 1.61 | | | | | |
| $f(\text{Year}, SB/B_{\text{pa}})$ | 7.61 | 39.16 | | | | | |
| $f(\text{Year}, \bar{F}_{\text{lag1}}/F_{\text{msy}})$ | 5.06 | 3.71 | | | | | |
| $f(\text{Year}, AMO_{\text{lag1}})$ | 7.15 | 3.09 | | | | | |
| $\beta_0$ | | | -5.16 | (-5.25, -5.07) | -118.0 | | |
| stock |  |  |  |  |  | 0.02 | 0.16 |
| ecoregion/year |  |  |  |  |  | 0.01 | 0.11 |
| ecoregion |  |  |  |  |  | 0.01 | 0.08 |
| non-recovering |  |  |  |  |  |  |  |
| $e_{RS} \sim f(\text{Year}) + f(\text{Year}, SA_{\text{mean}}) + f(\text{Year}, SB/B_{\text{pa}})$ | | | | | | | |
| $+ f(\text{Year}, \bar{F}_{\text{lag1}}/F_{\text{msy}}) + f(\text{Year}, wSST_{\text{anom}})$ | | | | | | | |
| $f(\text{Year})$ | 1.00 | 2.10 | | | | | |
| $f(\text{Year}, SA_{\text{mean}})$ | 7.01 | 141.36 | | | | | |
| $f(\text{Year}, SB/B_{\text{pa}})$ | 6.65 | 4.88 | | | | | |
| $f(\text{Year}, \bar{F}_{\text{lag1}}/F_{\text{msy}})$ | 3.60 | 1.96 | | | | | |
| $f(\text{Year}, wSST_{\text{anom}})$ | 3.02 | 1.52 | | | | | |
| $\beta_0$ | | | -5.16 | (-5.36, -4.96) | -50.2 | | |
| stock |  |  |  |  |  | 0.09 | 0.31 |
| ecoregion/year |  |  |  |  |  | 0.008 | 0.088 |
| ecoregion | | | | | | $3.4 \times 10^{-7}$ | $5.8 \times 10^{-4}$ |

Ecoregion-scale analyses on recruitment success revealed patterns largely consistent with the regional analyses with some system-specific deviations (Fig. S11). In ecoregions with non-recovering stocks like the Baltic, Celtic, and North Seas the density-dependent effect on recruitment on average became weakened over time; in contrast, the effect in the Norwegian– Barents Seas (with only recovering stocks) became strengthened (Fig. S11c). Correlations with spawner age structure metrics were detected in all but the Bay of Biscay and Faroes (Fig. S11b); in the Baltic Sea a positive correlation with age diversity became detectable in the early 1990s when fishing pressures began declining (Fig. 2). The time-varying effect of spawner age was especially pronounced in the Norwegian–Barents Seas stocks, showing a shift from positive to negative in the 1990s (Fig. S11b). An indirect fishing effect was (weakly) supported in the North Sea stocks only (Table S3 and Fig. S11d). Climate anomalies (in the year of spawning) contributed to variability in recruitment success in all ecoregions except the North Sea but the indices with the most statistical support varied among ecoregions (Fig. 5 and Table S3) despite high correlations among climate indices (Fig. S12). The ways climate anomalies contributed to recruitment variability also differed (Fig. 5). In the Baltic Sea and Faroes correlations with climate anomalies remained largely positive during the past few decades (Fig. 5a,d). In contrast, the positive effect in the Norwegian–Barents Seas began diminishing in the recent decades (Fig. 5f). In the Bay of Biscay and Icelandic waters the climate effect shifted from positive to negative (Fig. 5b,e).

**Figure 5.**
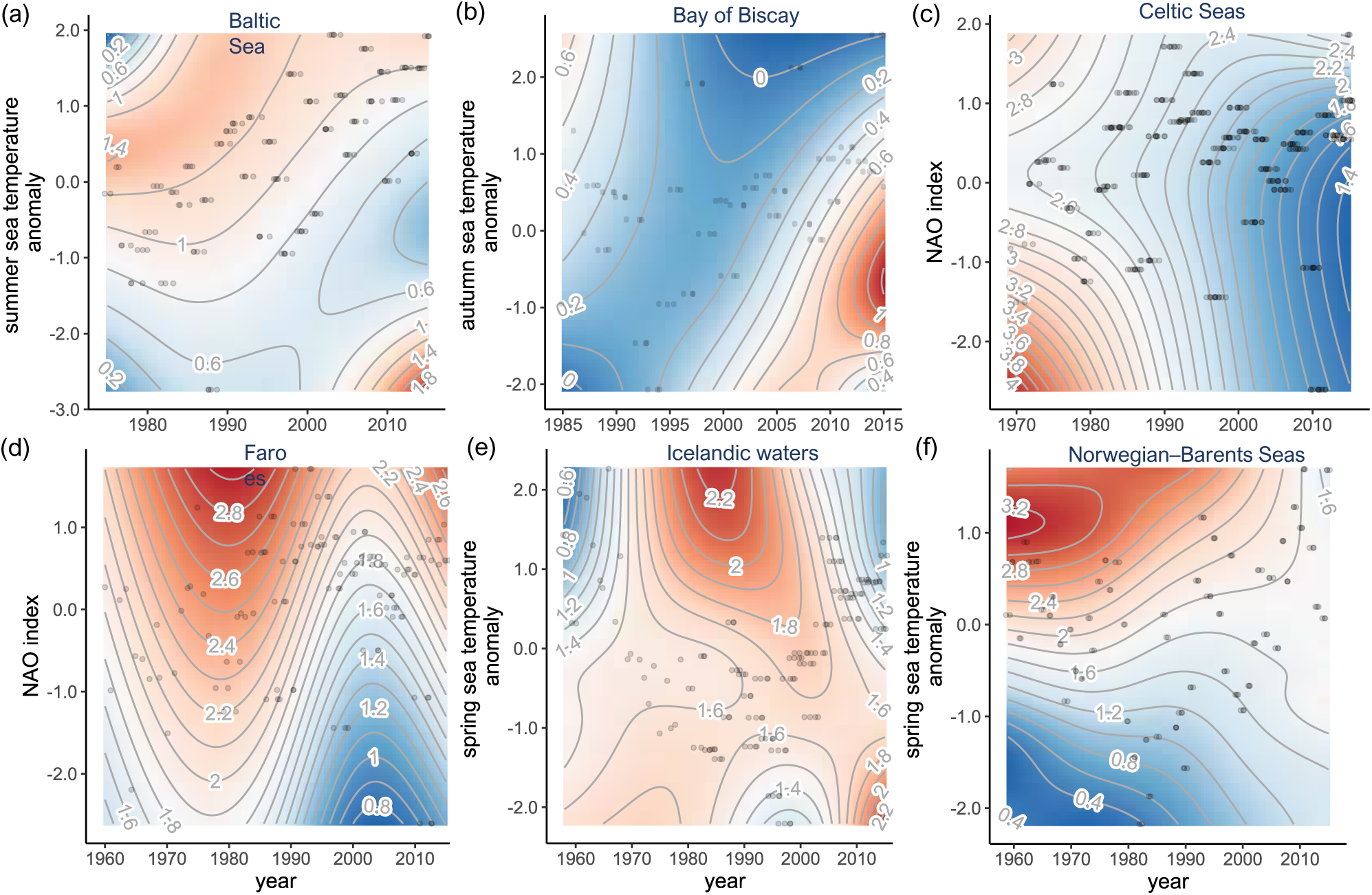
Ecoregion-scale climate effects on per capita recruitment success (fecundity) of northeast Atlantic fish stocks (the Baltic Sea, the Bay of Biscay, the Celtic Seas, Faroes, Icelandic waters, and the Norwegian–Barents Seas) during 1946–2016. Time-varying effects of climate condition are estimated by generalized additive mixed effects modeling with year (and stock for the Celtic Seas) as a random effect (see Fig. S11 for complete results of the models with the other covariates, spawner age structure, spawner biomass, and fishing pressure). Climate covariates selected in the models are indicated in the y-axis labels. Color gradients and contour lines indicate the partial effects of covariates estimated (as tensor interaction terms) in the models. Warmer colors indicate higher values, and the ranges of the partial effects differ among the covariates and are indicated as values on the contour lines. Gray circles indicate back- transformed input data.

Trend analyses revealed consistent temporal patterns in elasticities to vital rates for the recovering and non-recovering stocks. Both showed declining elasticities to (post-recruitment) juvenile survival rate (ΔAIC = 15.6 and 18.9) and increasing elasticities to adult survival rate over time (ΔAIC = 17.0 and 6.7) (Fig. S13). But the regionwide analyses detected a declining trend in elasticities to recruitment success rate in the recovering stocks only (ΔAIC = 3.6, Fig. 6a) with less among-stock variances than recruitment success (Fig. 6f-h). Ecoregion-specific variances in all the models fluctuated over years, but on average elasticities to adult survival showed higher deviations from the regional trends than the other vital rates (Fig. S13).

**Figure 6.**
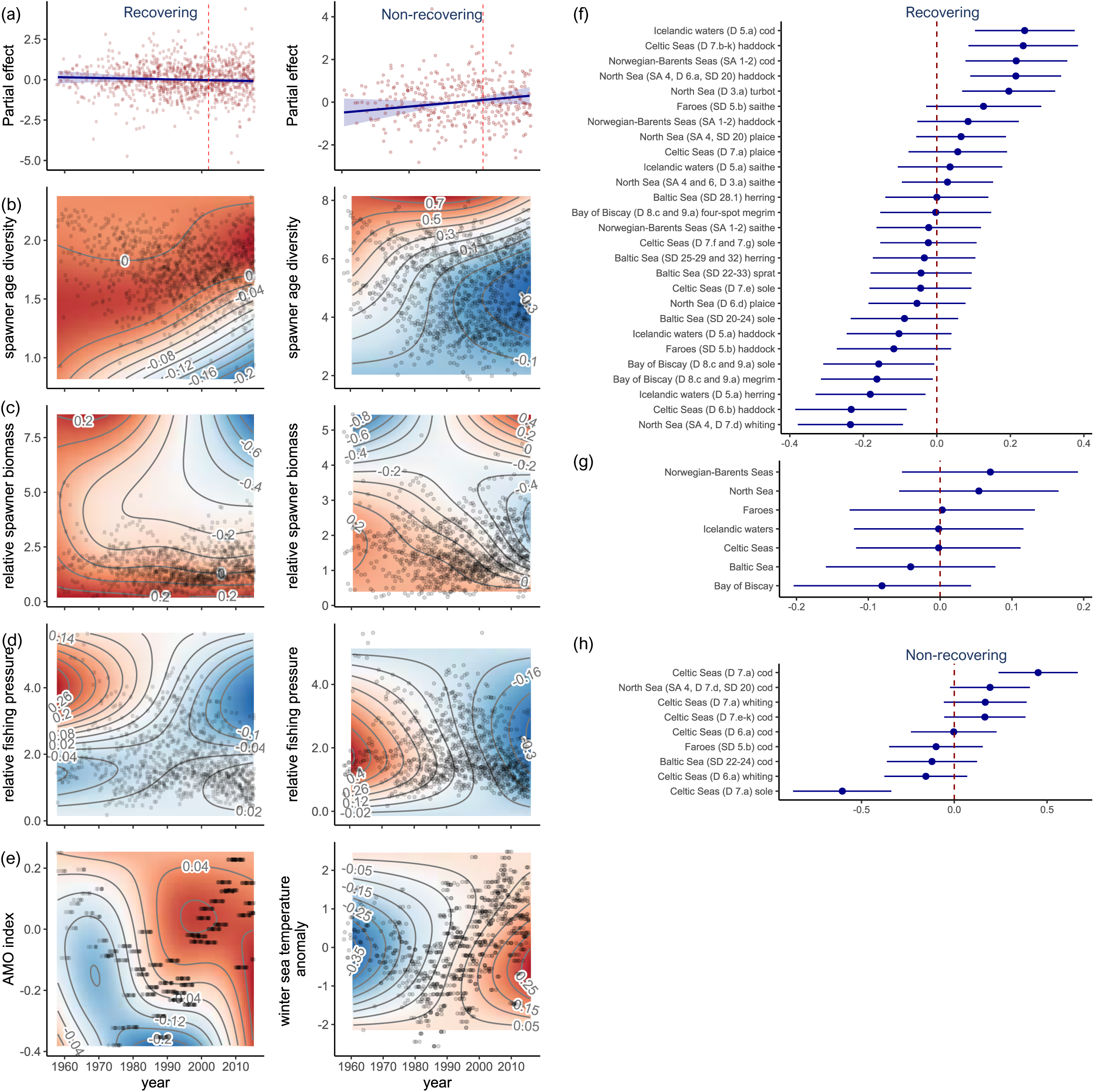
Regional trends in and covariates of elasticity to per capita recruitment success of the recovering and non-recovering northeast Atlantic fish stocks during 1946–2016. (a) Temporal trends. Solid lines and ribbons indicate temporal trends and 95% confidence intervals estimated by generalized additive mixed effects modeling with year as a fixed effect, and ecoregion, year within ecoregion, and stock as random effects. Circles indicate partial residuals estimated by the models. (b–e) Time-varying effects of spawner age structure, spawner biomass, fishing pressure, and climate condition. Covariates selected in the models are indicated in the y-axis labels. Color gradients and contour lines indicate the partial effects of covariates estimated (as tensor interaction terms) in the models. Warmer colors indicate higher values, and the ranges of the partial effects differ among the covariates and are indicated as values on the contour lines. Gray circles indicate back-transformed input data. (f–h) Random effects estimates (±95% confidence intervals) for stock and ecoregion for the recovering and non-recovering stocks (ecoregion- specific parameters for the non-recovering stocks are negligible; random effect estimates of year nested within ecoregion are given in Fig. S6c).

Regionwide analyses on elasticities to recruitment success largely reflected the time-varying responses in recruitment to intrinsic and extrinsic drivers. For the recovering stocks the model indicated that more diverse ages increasingly contributed to elasticities in more recent years, whereas the contributions of mean ages gradually increased in the non-recovering stocks (Fig. 6b). Elasticities to recruitment declined with higher spawner biomass in both stock groups, but the density-dependent effect in the non-recovering stocks diminished over time (Fig 6c). The model with an indirect fishing effect in both stock groups showed a declining contribution to variations in the elasticities after the reductions in fishing pressure (Fig. 6d). In contrast, the models for both stock groups indicated an increasingly more positive response to climate anomalies (the AMO index and winter sea temperature anomaly) over time (Fig. 6e).

Ecoregion-scale analyses on elasticities to recruitment success detected declining trends in all but the Bay of Biscay and the Norwegian–Barents Seas (Fig S14a), which showed little change in trends. The analyses also indicated spatial heterogeneity in the ways time-varying intrinsic and extrinsic drivers contributed to variations in the elasticities. For example, shifting contributions of spawner age structure were detected in most ecoregions, showing both diminishing (e.g., Faroes and the North Sea) and increasing (e.g., the Baltic Sea and Icelandic waters) trends (Fig. S14b). Negative correlations with spawner biomass were consistently detected except in the Baltic Sea (Fig. S14c). This maternal effect increased over time except in the Norwegian– Barents Seas, which showed the opposite trend (Fig. S14c). The models with an indirect fishing effect had statistical support for Faroes, and the Celtic, North, and Norwegian–Barents Seas; the fishing effect declined and/or shifted over time except the Norwegian–Barents Sea, which showed an increasing contribution of fishing since the 1990s (Fig. S14d). In the Baltic Sea and the Bay of Biscay a positive climate effect emerged in the 2000s (Fig. 7a,b), reflecting the temporal patterns of shifting local climate effects on recruitment success (Fig. 5a,b). In Faroes and Icelandic waters, where climate anomalies (the NAO index and spring sea temperature) had a positive effect on recruitment, the elasticities were inversely correlated with the indices, but this effect shifted to the opposite in the 1990s–2000s (Fig. 7c,d).

**Figure 7.**
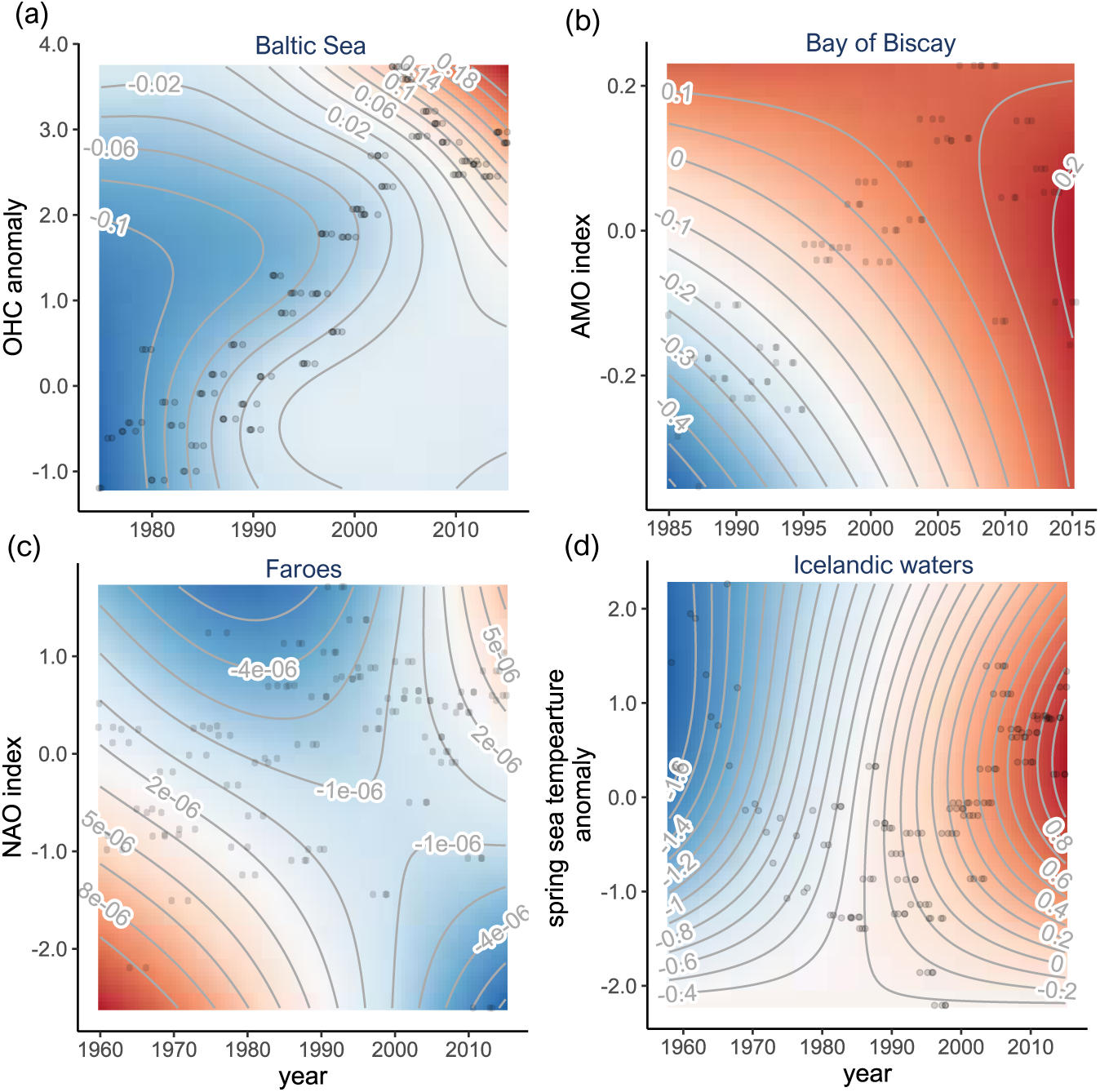
Ecoregion-scale climate effects on elasticity to per capita recruitment success of northeast Atlantic fish stocks (the Baltic Sea, the Bay of Biscay, Faroes, and Icelandic waters) during 1946–2016. Time-varying effects of climate condition are estimated by generalized additive mixed effects modeling with year (and stock for the Celtic Seas) as a random effect (see Fig. S14 for complete results of the models with the other covariates, spawner age structure, spawner biomass, and fishing pressure). Climate covariates selected in the models are indicated in the y-axis labels. Color gradients and contour lines indicate the partial effects of covariates estimated (as tensor interaction terms) in the models. Warmer colors indicate higher values, and the ranges of the partial effects differ among the covariates and are indicated as values on the contour lines. Gray circles indicate back-transformed input data.

## Discussion

Analyses on the transient dynamics of northeast Atlantic fish stocks reveal that management- aided reductions in fishing pressure and shifting climate conditions nonlinearly shaped rebuilding patterns of these historically depleted stocks through restorations of demographic structure (reversing age truncation) and reproductive capacity in recent decades. As the survival rates and demographic structures of reproductive fish improved with declining fishing pressures, transient population growth rates became less sensitive to variability in recruitment and juvenile survival and more to that in adult survival, allowing the stocks to rebuild biomass despite declining recruitment success. As the biomass of reproductive fish rebuilt, recruitment success rates also became increasingly regulated by density-dependent processes involving increasingly higher numbers of older reproductive fish. Although warmer than average climates promoted recruitment success throughout the early 2000s in some ecoregions, warming trends over the past decades began adversely affecting reproductive performance overall. Variations in regulation of fishing pressure among stocks also led to diverging trends in recoveries of the stocks exposed to a range of local climate conditions. Insufficient or delayed reductions in fishing pressure can further deplete a stock and erode reproductive fish demographic structure, thereby diminishing fecundity and thus recovery potential, while also remaining sensitive to climate variability.

Rebuilding biomass by accounting for demographic structure through management are thus not only key to sustainable use of marine resource populations but also the foundation for effective applications of adaptive measures resilient to future environmental change.

### Transient dynamics of recovering populations from intense fishing pressures

Fishing-induced changes in the demography and reproductive capacity of an exploited population can be reversible when management measures are effectively applied to regulate fishing pressure. This pattern is displayed especially in the Norwegian–Barents Seas, where fishing pressures were reduced by more than 60% (compared to the regional average of ∼50%) since the early 1990s, followed by sharp rises in the average age and biomass of reproductive fish. But when a population experiences long-term intensive harvesting, recovery time can become longer (Hutchings 2015) and more uncertain (Neubauer *et al*. 2013), with the population remaining in long transient states (Hastings *et al*. 2018). Although fishing pressures on many northeast Atlantic fish stocks precipitously declined through stricter management measures applied since the late 1990s, demographic restorations and stock recoveries on average progressed gradually. Reducing fishing pressures can allow more adults to survive and contribute to the production of new recruits but the restoration of the demographic structure takes place over generations and may take longer (a decade or more) for severely depleted stocks. Further, although the declining trends in reproductive fish biomass halted in the early 2000s in most ecoregions, reduction levels in fishing pressure varied among ecoregions and stocks and were reflected in recovery patterns. When reductions in fishing pressure were insufficient or delayed, the declining trends in reproductive capacity and deterioration of demographic structure in some stocks continued. Slow recoveries of severely depleted fish stocks are reportedly widespread (Hutchings 2000). Regionwide analyses on the non-recovering stocks in the northeast Atlantic reveal that sustaining high fishing pressures for even a few more years can result in vastly different recovery patterns, with further age truncations and reductions in reproductive capacity, which could further diminish fishery yields (Hilborn *et al*. 2020).

Removal of some biomass from a population by harvesting can promote its productivity (Neubauer *et al*. 2013) by attenuating density-dependent effects on early life stage processes (competition for food and space for example, Anderson *et al*. 2008). Intense fishing pressures before the implementation of stricter management measures–mediated through reproductive fish–may have led to higher recruitment rates in depleted stocks in the northeast Atlantic Ocean. Fishing not only truncates age structure but induces changes in life-history traits that may influence recovery (Neubauer *et al*. 2013; Hutchings & Kuparinen 2020). The generation times of these depleted northeast Atlantic stocks were shortened (by 14 to 50%), which is often associated with high year-to-year variability in recruitment (Minto, Myers & Blanchard 2008) and high average recruitment rates (Bjørkvoll *et al*. 2012). Higher growth rates of depleted (and age-truncated) fish populations also likely benefited during the early stages of recovery as more reproductive fish contributed to spawning events, thereby increasing the probability of recruitment success. These indirect effects, however, diminished over time following the reductions of fishing pressures. Despite the signs of recovery in reproductive fish, per capita recruitment success rates on average declined in the northeast Atlantic stocks, as also reported in other Atlantic ecoregions (Britten, Dowd & Worm 2016). But these apparent patterns were shaped by different processes among stocks. In the recovering stocks density dependency remained present in recruitment success throughout the past decades and its strength increased as reproductive fish biomass rose, likely slowing recoveries as fishing pressures declined. In contrast, in the non-recovering stocks density dependency in recruitment success emerged only after the reductions in fishing pressure but its strength continued to decline with reproductive fish biomass, suggesting contributions of other (extrinsic) drivers.

Density dependency in early life also perhaps explains discontinuous relationships (shifting from positive to negative) between reproductive fish age and per capita recruitment success in the recovering stocks (especially in the Norwegian–Barents Seas) following the reductions in fishing pressure. Higher mean age and age diversity in reproductive fish are expected to promote recruitment success (Rogers & Dougherty 2019). Age-dependent variations in spawning pattern can influence egg viability and larval survival; older spawners tend to produce more viable offspring (Hixon, Johnson & Sogard 2014). But the analyses on the northeast Atlantic fishes show that only the non-recovering (age-truncated) stocks kept positive relationships between reproductive fish age structure and recruitment success. In the recovering stocks, density dependency may have played a larger role than maternal effects, revealing a shift in compensatory processes. As adult survival improves and population density increases, pre-recruit survival may decline (owing to competition for food and other resources including cannibalism by adults), contributing to apparent increases in average ages of reproductive fish.

Short-term (year-to-year) fluctuations in recruitment can propagate through transient dynamics of a population when its (demographic) buffering capacity is compromised by age truncation (Anderson *et al*. 2008). Truncated age distributions in reproductive fish may produce volatile population behaviors that reflect recruitment pulses (Anderson *et al*. 2008). As adult survival improved and stock biomass rebuilt in many northeast Atlantic fish stocks, the transient growth became less sensitive to recruitment variability and more sensitive to adult survival. The regionwide analyses show that increases in transient growth rate and biomass during the past few decades resulted largely from improved survival of older fish. Greater sensitivities to adult survival also may make harvested populations more manageable. Nonlinear responses to age structure in reproductive fish is however also evident in sensitivity to recruitment variability.

Although the buffering effect of increasingly age-diverse reproductive fish emerged in many ecoregions especially in the early stages of recovery, this relationship also shifted as the recoveries further progressed (as seen in the Norwegian–Barents Seas fishes), likely responding to other intrinsic or extrinsic dynamics (such as catch limits being raised as stock biomass rebuilds).

### Climate regulation of demographic variations

Fish populations experiencing persistent selective depletions of older adults often become sensitive to variations in early life-history stage processes that reflect signals in environmental conditions like climate variability (Ottersen, Hjermann & Stenseth 2006; Rouyer *et al*. 2011). Warmer than average climate conditions in the northeast Atlantic Ocean increasingly contributed to observed patterns in fish recruitment success, reflecting shifting climate effects on recruitment processes. For example, stratification becomes intensified as climate warms, limiting nutrient movement to deeper waters and food accessibility for the early life of demersal and benthic species (Moore *et al*. 2018). At the region scale, however, the models captured only weak responses in recruitment to regional climate signals likely because of heterogeneous warming patterns among ecoregions, as also seen in other parts of the world (Barange *et al*. 2018).

Region-scale analyses may mask differential effects of local climate and oceanographic features (climate-induced shifts in ocean heat content and primary productivity for example) that marine species are adapted to and experience (Conover 1992).

Ecoregion-scale variability in recruitment success may better reflect how fish reproductive behavior responds to regional climate mediated by local habitat conditions, as hypothesized in past research (Myers 1998; Ottersen & Stenseth 2001). For stocks in high latitude systems like the Norwegian–Barents Seas and Icelandic waters, for example, climate conditions in winter– spring (December–May) that regulate phonologies like timings of spawning migration (Sundby & Nakken 2008) contributed to recruitment variability. Warming-induced reductions in sea ice extent in these ecoregions promoted ecosystem productivity in the late 1990s and early 2000s and may have benefited upper trophic animals like fish (Stenevik & Sundby 2007), likely through changes in growth rate and life stage duration (Houde 2016). When spawning times coincide with the period of high prey abundance, pre-recruit fish can grow more and experience less size-dependent mortalities (like predation), contributing to greater recruitment success (Peck, Huebert & Llopiz 2012). In contrast, in semi-enclosed systems like the Baltic Sea warming-induced changes in nearshore habitat conditions in summer (June–August) (lower salinity caused by higher precipitation and freshwater inflows for example, MacKenzie *et al*. 2007) may have played a greater role in regulating recruitment success through biophysical processes during the early years of life.

Demographic responses by fish populations to changing climate conditions also may vary discontinuously (Brander 2007). Local and regional climate conditions (both favorable and unfavorable) that early life-history stage fish experience can persist through time (Ottersen *et al*. 2001) and may emerge as delayed effects on transient population dynamics. In some northeast Atlantic ecoregions the ways that climate affects fish recruitment success shifted from positive to negative and may have slowed their recoveries. These shifts suggest that even in the ecoregions where warming climate promoted productivity in the early twentieth century, fish began to experience adverse (direct and indirect) effects. Mobile species can adjust their movement (deeper and poleward) to mitigate physiological stress from warming upper water columns (Pinsky *et al*. 2013; Engelhard, Righton & Pinnegar 2014). But asynchronous shifts in spatial distribution by spawning fish and seasonal events in lower trophic groups like spring plankton blooms (phenological mismatches) also could trigger a series of events unfavorable for early life fish (Asch & Erisman 2018), shrinking the probability of recruitment success.

Climate variability that regulates local ocean conditions may propagate through vital rates to shape transient population dynamics of marine species (Jenouvrier *et al*. 2018). Depleted populations in particular may have heightened sensitivities to adverse environmental conditions (Hidalgo *et al*. 2011; Rouyer *et al*. 2011), which are becoming increasingly more variable in recent decades (Rathore *et al*. 2020). Although the sensitivities of transient growth to recruitment variability in the northeast Atlantic fish stocks declined as they recovered, the analyses reveal that increasingly warmer than average climates began influencing how recruitment variability propagates through transient dynamics. For example, in high-latitude systems like Faroes, where fish stocks become less sensitive to recruitment variability under a warmer climate, this stabilizing effect gradually diminished (and shifted to the opposite) as the climate further warmed. In contrast, in the Baltic Sea, where fish stocks become more sensitive to recruitment variability under a warmer climate, this amplifying effect became strengthened as the climate further warmed. Climate effects on transient dynamics also may, however, depend on population size and demographic structure. In the Norwegian–Barents Seas, where fish stocks experienced higher reductions in fishing pressure early and thus higher biomass rebuilding rates than other stocks, the stocks responded to recruitment variability independently of climate conditions.

### Transients as an integrated measure of effects of fishing, management, and climate

Analysis of transient population dynamics can reveal how time-varying demographic structure and vital rates contribute to recovery patterns of managed resource populations when also exposed to low-frequency, nonstationary drivers like climate. Declined sensitivities to highly variable demographic parameters like recruitment, as displayed by intensely managed northeast Atlantic fish stocks in this study, can provide a buffer against rising environmental variability.

Increasing sensitivities to adult survival further suggest that the stocks also are expected to become more responsive to management actions through regulation of fishing pressure (enforcing positive feedback), and future management plans would benefit from maintaining healthy (age-diverse) reproductive populations of exploited species (Pinsky & Byler 2015). Combined with other management measures (like use of marine protected areas and fish size limits, White *et al*. 2013; Hixon, Johnson & Sogard 2014), restoring and maintaining the demographic structure of an exploited species thus may enhance resilience against destabilizing effects of environmental variability amplified by climate change.

Emerging non-negligible changes in demographic responses to rising climate variability nevertheless may influence the efficacy of fisheries management. Although historical climate conditions promoted the productivity of fish stocks in the northeast Atlantic Ocean, some stocks in this region and elsewhere in the Northern Hemisphere already are experiencing adverse effects of changing climate (Pershing *et al*. 2015; Free *et al*. 2019). As many biological responses to climate-driven changes in surface temperature and other ocean properties are often nonlinear, we would expect continued shifts in how climate influences marine resource populations and management (Britten, Dowd & Worm 2016). Accounting for these climate-induced changes thus may not only reduce uncertainties in assessment but also improve the performance of management measures (Rouyer et al. 2011; Bahri et al. 2021).

Exploited marine fish species are an integral part of an ecosystem and changes in their abundances and demographic structures may propagate through community and ecosystem dynamics (Jennings & Kaiser 1998), triggering a series of changes in ecologically connected species (Goto *et al*. 2022; Pérez-Rodríguez *et al*. 2022). As adult survival improves in historically overharvested populations, ecological processes including resource competition and predation also may indirectly modify vital rates and demographic structures of non-target species (van Gemert & Andersen 2018). Improving our understanding of how warming and more variable climates continue to reshape the transient dynamics (through changes in life history traits and demographics) would help develop management plans to mitigate the risk of overexploitation and safeguard ecosystem services against rising environmental variability.

## Acknowledgements

This work was partially supported by the Institute of Marine Research’s Reduced Uncertainty in Stock Assessment (REDUS) project.

## Data Availability Statement

The data that support the findings of this study are openly available in ICES Scientific Reports at https://ices-library.figshare.com/, reference number ICES CM 2017/ACOM:06, ICES CM 2017/ACOM:08, ICES CM 2017/ACOM:11, ICES CM 2017/ACOM:12, ICES CM 2017/ACOM:13, and ICES CM 2017/ACOM:21, and in the data archives of Met Office Hadley Centre, and NOAA Physical Sciences Laboratory and National Centers for Environmental Information at https://www.metoffice.gov.uk, https://psl.noaa.gov, and https://www.ncei.noaa.gov. Code and derived datasets (age-specific demographic parameter estimates) will be available upon publication.

