## Supporting Information for "Transient demographic dynamics of recovering fish populations shaped by past climate variability, harvest, and management"

**Table S1**. Results of model selection for random (intercept) effects (year, ecoregion, and stock) for recruitment success ($RS_{total}$), and elasticity to recruitment success ($e_{RS}$) as response variables for the recovering and non-recovering northeast Atlantic fish stocks. The models in bold indicate the best models for response variable and recovery status combinations. DF and $\chi^2$ and $\chi^2$ DF indicate the degrees of freedom in generalized additive mixed effects modeling, the Chi-square statistic and the degrees of freedom from likelihood ratio tests.

| random effect | DF | log-likelihood | $\chi^2$ | $\chi^2$ DF | $p(>\chi^2)$ |
|---|---|---|---|---|---|
| $RS_{total}$: recovering stocks | | | | | |
| none | 10 | -3829.4 | | | |
| year | 11 | -3811.2 | 36.322 | 1 | $1.67\times10^{-10}$ |
| year + ecoregion | 12 | -3605 | 412.534 | 1 | $< 2.20\times10^{-16}$ |
| ecoregion + stock | 12 | -3312.6 | 584.677 | 0 | |
| year + stock | 12 | -3304.1 | 16.968 | 0 | |
| year/ecoregion | 12 | -3640.9 | 0 | 0 | |
| ecoregion/year | 12 | -3564.1 | 153.704 | 0 | |
| **ecoregion/year + stock** | **13** | **-3258.1** | **612.009** | **1** | $\mathbf{< 2.20\times10^{-16}}$ |
| year + ecoregion + stock | 13 | -3303.4 | 0 | 0 | |
| year/ecoregion + stock | 13 | -3258.9 | 89.13 | 0 | |
| $RS_{total}$: non-recovering stocks | | | | | |
| none | 10 | -1281.5 | | | |
| year | 11 | -1276.6 | 9.8752 | 1 | 0.001675 |
| year + ecoregion | 12 | -1247.7 | 57.7703 | 1 | $2.95\times10^{-14}$ |
| ecoregion + stock | 12 | -1144.8 | 205.8769 | 0 | |
| year + stock | 12 | -1125.7 | 38.2662 | 0 | |
| year/ecoregion | 12 | -1269.5 | 0 | 0 | |
| ecoregion/year | 12 | -1244.7 | 49.5987 | 0 | |
| **ecoregion/year + stock** | **13** | **-1106.6** | **276.102** | **1** | $\mathbf{< 2.20\times10^{-16}}$ |
| year + ecoregion + stock | 13 | -1125.7 | 0 | 0 | |
| year/ecoregion + stock | 13 | -1105.5 | 40.2604 | 0 | |
| $e_{RS}$: recovering stocks | | | | | |
| none | 4 | 5242 | | | |
| year | 5 | 5251.1 | 18.138 | 1 | $2.55\times10^{-7}$ |
| year + ecoregion | 6 | 5293 | 83.859 | 1 | $< 2.20\times10^{-16}$ |
| ecoregion + stock | 6 | 5344.7 | 103.424 | 0 | |
| year + stock | 6 | 5354.6 | 19.719 | 0 | |
| year/ecoregion | 6 | 5294.8 | 0 | 0 | |
| ecoregion/year | 6 | 5312.8 | 35.987 | 0 | |
| **ecoregion/year + stock** | **7** | **5388.8** | **152.06** | **1** | $\mathbf{< 2.20\times10^{-16}}$ |
| year + ecoregion + stock | 7 | 5354.6 | 0 | 0 | |
| year/ecoregion + stock | 7 | 5389.9 | 70.523 | 0 | |
| $e_{RS}$: non-recovering stocks | | | | | |
| none | 10 | 1558.1 | | | |
| year | 11 | 1564.3 | 12.459 | 1 | 0.000416 |
| year + ecoregion | 12 | 1579 | 29.225 | 1 | $6.44\times10^{-8}$ |
| ecoregion + stock | 12 | 1607.4 | 56.947 | 0 | |
| year + stock | 12 | 1623.7 | 32.474 | 0 | |
| year/ecoregion | 12 | 1573.8 | 0 | 0 | |
| ecoregion/year | 12 | 1583.5 | 19.466 | 0 | |
| **ecoregion/year + stock** | **13** | **1630.8** | **94.592** | **1** | $\mathbf{< 2.20\times10^{-16}}$ |
| year + ecoregion + stock | 13 | 1623.7 | 0 | 0 | |
| year/ecoregion + stock | 13 | 1632.6 | 17.815 | 0 | |

**Table S2**. Results of model selection for covariates of recruitment success ($RS_{total}$) and elasticity to recruitment success ($e_{RS}$) as response variables for the recovering and non-recovering northeast Atlantic fish stocks. The models in bold indicate the best models for response variable and recovery status combinations. DF, AIC, ΔAIC, and *w* indicate the degrees of freedom, Akaike information criterion scores, differences in AIC scores from the best models, and model weights. All time (year)-dependent smoother terms for covariates are modeled as tensor product construction (using the *t2* function from the R package *mgcv* for $RS_{total}$ and the *te* function for $e_{RS}$).

| model | DF | AIC | ΔAIC | *w* | log-likelihood |
|---|---|---|---|---|---|
| $RS_{total}$: recovering stocks | | | | | |
| $RS_{total}$ ~ $f$(Year) | 1002 | 6353.2 | 165.5 | 0.00 | -3169.6 |
| $RS_{total}$ ~ $f$(Year) + $f$(Year, $SA_{mean}$) | 997 | 6307.2 | 119.5 | 0.00 | -3141.6 |
| $RS_{total}$ ~ $f$(Year) + $f$(Year, $SA_{diversity}$) | 997 | 6334.7 | 147.0 | 0.00 | -3155.3 |
| $RS_{total}$ ~ $f$(Year) + $f$(Year, $SA_{mean}$) + $f$(Year, SB) | 991 | 6247.5 | 59.8 | 0.00 | -3106.7 |
| $RS_{total}$ ~ $f$(Year) + $f$(Year, $SA_{mean}$) + $f$(Year, SB/$B_{pa}$) | 991 | 6190.4 | 2.7 | 0.08 | -3078.2 |
| **$RS_{total}$ ~ $f$(Year) + $f$(Year, $SA_{mean}$) + $f$(Year, SB/$B_{pa}$) + $f$(Year, $\bar{F}_{lag1}$)** | **986** | **6187.7** | **0.0** | **0.32** | **-3071.9** |
| $RS_{total}$ ~ $f$(Year) + $f$(Year, $SA_{mean}$) + $f$(Year, SB/$B_{pa}$) + $f$(Year, $\bar{F}_{lag1}/F_{msy}$) | 986 | 6192.7 | 5.0 | 0.03 | -3074.4 |
| $RS_{total}$ ~ $f$(Year) + $f$(Year, $SA_{mean}$) + $f$(Year, SB/$B_{pa}$) + $f$(Year, $wSST_{anom}$) | 981 | 6197.1 | 9.4 | 0.00 | -3071.6 |
| $RS_{total}$ ~ $f$(Year) + $f$(Year, $SA_{mean}$) + $f$(Year, SB/$B_{pa}$) + $f$(Year, $spSST_{anom}$) | 981 | 6192.1 | 4.4 | 0.04 | -3069.0 |
| $RS_{total}$ ~ $f$(Year) + $f$(Year, $SA_{mean}$) + $f$(Year, SB/$B_{pa}$) + $f$(Year, $smSST_{anom}$) | 981 | 6192.3 | 4.6 | 0.03 | -3069.2 |
| $RS_{total}$ ~ $f$(Year) + $f$(Year, $SA_{mean}$) + $f$(Year, SB/$B_{pa}$) + $f$(Year, $aSST_{anom}$) | 981 | 6195.5 | 7.8 | 0.01 | -3070.8 |
| $RS_{total}$ ~ $f$(Year) + $f$(Year, $SA_{mean}$) + $f$(Year, SB/$B_{pa}$) + $f$(Year, NAO) | 981 | 6190.2 | 2.5 | 0.09 | -3068.1 |
| $RS_{total}$ ~ $f$(Year) + $f$(Year, $SA_{mean}$) + $f$(Year, SB/$B_{pa}$) + $f$(Year, AMO) | 981 | 6195.2 | 7.5 | 0.01 | -3070.6 |
| $RS_{total}$ ~ $f$(Year) + $f$(Year, $SA_{mean}$) + $f$(Year, SB/$B_{pa}$) + $f$(Year, OHC) | 983 | 6190.9 | 3.2 | 0.06 | -3070.4 |
| $RS_{total}$ ~ $f$(Year) + $f$(Year, $SA_{mean}$) + $f$(Year, SB/$B_{pa}$) + $f$ Year, ($NAO_{lag1}$) | 981 | 6193.8 | 6.1 | 0.02 | -3069.9 |
| $RS_{total}$ ~ $f$(Year) + $f$(Year, $SA_{mean}$) + $f$(Year, SB/$B_{pa}$) + $f$(Year, $AMO_{lag1}$) | 981 | 6188.4 | 0.7 | 0.23 | -3067.2 |
| $RS_{total}$ ~ $f$(Year) + $f$(Year, $SA_{mean}$) + $f$(Year, SB/$B_{pa}$) + $f$(Year, $OHC_{lag1}$) | 983 | 6193.7 | 6.0 | 0.02 | -3071.9 |
| $RS_{total}$: non-recovering stocks | | | | | |
| $RS_{total}$ ~ $f$(Year) | 360 | 2655.1 | 66.8 | 0.00 | -1320.5 |
| $RS_{total}$ ~ $f$(Year) + $f$(Year, $SA_{mean}$) | 355 | 2631.8 | 43.5 | 0.00 | -1303.9 |
| $RS_{total}$ ~ $f$(Year) + $f$(Year, $SA_{diversity}$) | 355 | 2651.7 | 63.4 | 0.00 | -1313.9 |
| $RS_{total}$ ~ $f$(Year) + $f$(Year, $SA_{mean}$) + $f$(Year, SB) | 350 | 2612.5 | 24.2 | 0.00 | -1289.3 |
| $RS_{total}$ ~ $f$(Year) + $f$(Year, $SA_{mean}$) + $f$(Year, SB/$B_{pa}$) | 350 | 2588.8 | 0.5 | 0.35 | -1278.1 |
| $RS_{total}$ ~ $f$(Year) + $f$(Year, $SA_{mean}$) + $f$(Year, SB/$B_{pa}$) + $f$(Year, $\bar{F}_{lag1}$) | 345 | 2593.5 | 5.2 | 0.03 | -1274.7 |
| $RS_{total}$ ~ $f$(Year) + $f$(Year, $SA_{mean}$) + $f$(Year, SB/$B_{pa}$) + $f$(Year, $\bar{F}_{lag1}/F_{msy}$) | 346 | 2592.8 | 4.5 | 0.05 | -1277.4 |
| $RS_{total}$ ~ $f$(Year) + $f$(Year, $SA_{mean}$) + $f$(Year, SB/$B_{pa}$) + $f$(Year, $wSST_{anom}$) | 345 | 2594.1 | 5.8 | 0.02 | -1275.1 |
| $RS_{total}$ ~ $f$(Year) + $f$(Year, $SA_{mean}$) + $f$(Year, SB/$B_{pa}$) + $f$(Year, $spSST_{anom}$) | 345 | 2596.1 | 7.8 | 0.01 | -1276.1 |
| $RS_{total}$ ~ $f$(Year) + $f$(Year, $SA_{mean}$) + $f$(Year, SB/$B_{pa}$) + $f$(Year, $smSST_{anom}$) | 345 | 2595.9 | 7.6 | 0.01 | -1276.0 |
| $RS_{total}$ ~ $f$(Year) + $f$(Year, $SA_{mean}$) + $f$(Year, SB/$B_{pa}$) + $f$(Year, $aSST_{anom}$) | 345 | 2595.3 | 7.0 | 0.01 | -1275.7 |
| $RS_{total}$ ~ $f$(Year) + $f$(Year, $SA_{mean}$) + $f$(Year, SB/$B_{pa}$) + $f$(Year, NAO) | 345 | 2597.6 | 9.3 | 0.00 | -1276.8 |
| $RS_{total}$ ~ $f$(Year) + $f$(Year, $SA_{mean}$) + $f$(Year, SB/$B_{pa}$) + $f$(Year, AMO) | 345 | 2597.8 | 9.5 | 0.00 | -1276.9 |
| **$RS_{total}$ ~ $f$(Year) + $f$(Year, $SA_{mean}$) + $f$(Year, SB/$B_{pa}$) + $f$(Year, OHC)** | **347** | **2588.3** | **0.0** | **0.45** | **-1272.8** |
| $RS_{total}$ ~ $f$(Year) + $f$(Year, $SA_{mean}$) + $f$(Year, SB/$B_{pa}$) + $f$ Year, ($NAO_{lag1}$) | 345 | 2593.4 | 5.1 | 0.04 | -1274.7 |
| $RS_{total}$ ~ $f$(Year) + $f$(Year, $SA_{mean}$) + $f$(Year, SB/$B_{pa}$) + $f$(Year, $AMO_{lag1}$) | 345 | 2598.4 | 10.1 | 0.00 | -1277.2 |
| $RS_{total}$ ~ $f$(Year) + $f$(Year, $SA_{mean}$) + $f$(Year, SB/$B_{pa}$) + $f$(Year, $OHC_{lag1}$) | 345 | 2595.3 | 7.0 | 0.01 | -1275.7 |
| $e_{RS}$: recovering stocks | | | | | |
| $e_{RS}$ ~ $f$(Year) | 893 | -10276.5 | 93.1 | 0.00 | 5294.5 |
| $e_{RS}$ ~ $f$(Year) + $f$(Year, $SA_{mean}$) | 884 | -10287.7 | 81.8 | 0.00 | 5309.5 |
| $e_{RS}$ ~ $f$(Year) + $f$(Year, $SA_{diversity}$) | 887 | -10297.8 | 71.7 | 0.00 | 5312.5 |
| $e_{RS}$ ~ $f$(Year) + $f$(Year, $SA_{mean}$) + $f$(Year, SB) | 867 | -10338.1 | 31.4 | 0.00 | 5353.2 |
| $e_{RS}$ ~ $f$(Year) + $f$(Year, $SA_{mean}$) + $f$(Year, SB/$B_{pa}$) | 873 | -10352.2 | 17.3 | 0.00 | 5353.4 |
| $e_{RS}$ ~ $f$(Year) + $f$(Year, $SA_{mean}$) + $f$(Year, SB/$B_{pa}$) + $f$(Year, $\bar{F}_{lag1}$) | 872 | -10346.6 | 22.9 | 0.00 | 5355.2 |
| $e_{RS}$ ~ $f$(Year) + $f$(Year, $SA_{mean}$) + $f$(Year, SB/$B_{pa}$) + $f$(Year, $\bar{F}_{lag1}/F_{msy}$) | 870 | -10361.5 | 8.0 | 0.02 | 5362.7 |
| $e_{RS}$ ~ $f$(Year) + $f$(Year, $SA_{mean}$) + $f$(Year, SB/$B_{pa}$) + $f$(Year, $\bar{F}_{lag1}/F_{msy}$) + $f$(Year, $wSST_{anom}$) | 870 | -10361.5 | 8.0 | 0.02 | 5362.7 |
| $e_{RS}$ ~ $f$(Year) + $f$(Year, $SA_{mean}$) + $f$(Year, SB/$B_{pa}$) + $f$(Year, $\bar{F}_{lag1}/F_{msy}$) + $f$(Year, $spSST_{anom}$) | 878 | -10358.2 | 11.3 | 0.00 | 5354.9 |
| $e_{RS}$ ~ $f$(Year) + $f$(Year, $SA_{mean}$) + $f$(Year, SB/$B_{pa}$) + $f$(Year, $\bar{F}_{lag1}/F_{msy}$) + $f$(Year, $smSST_{anom}$) | 874 | -10352.5 | 17.0 | 0.00 | 5359.3 |
| $e_{RS}$ ~ $f$(Year) + $f$(Year, $SA_{mean}$) + $f$(Year, SB/$B_{pa}$) + $f$(Year, $\bar{F}_{lag1}/F_{msy}$) + $f$(Year, $aSST_{anom}$) | 878 | -10361.8 | 7.7 | 0.02 | 5357.2 |
| $e_{RS}$ ~ $f$(Year) + $f$(Year, $SA_{mean}$) + $f$(Year, SB/$B_{pa}$) + $f$(Year, $\bar{F}_{lag1}/F_{msy}$) + $f$(Year, NAO) | 871 | -10360.5 | 9.0 | 0.01 | 5362.9 |
| $e_{RS}$ ~ $f$(Year) + $f$(Year, $SA_{mean}$) + $f$(Year, SB/$B_{pa}$) + $f$(Year, $\bar{F}_{lag1}/F_{msy}$) + $f$(Year, AMO) | 870 | -10361.5 | 8.0 | 0.02 | 5362.7 |
| $e_{RS}$ ~ $f$(Year) + $f$(Year, $SA_{mean}$) + $f$(Year, SB/$B_{pa}$) + $f$(Year, $\bar{F}_{lag1}/F_{msy}$) + $f$(Year, OHC) | 873 | -10356 | 13.5 | 0.00 | 5361.1 |
| $e_{RS}$ ~ $f$(Year) + $f$(Year, $SA_{mean}$) + $f$(Year, SB/$B_{pa}$) + $f$(Year, $\bar{F}_{lag1}/F_{msy}$) + $f$(Year, $NAO_{lag1}$) | 868 | -10360.7 | 8.8 | 0.01 | 5366.6 |
| **$e_{RS}$ ~ $f$(Year) + $f$(Year, $SA_{mean}$) + $f$(Year, SB/$B_{pa}$) + $f$(Year, $\bar{F}_{lag1}/F_{msy}$) + $f$(Year, $AMO_{lag1}$)** | **884** | **-10369.5** | **0.0** | **0.90** | **5354.4** |

| | | | | | |
|---|---|---|---|---|---|
| $e_{RS} \sim f(\text{Year}) + f(\text{Year}, \text{SA}_{mean}) + f(\text{Year}, \text{SB}/B_{pa}) + f(\text{Year}, \bar{F}_{lag1}/F_{msy}) + f(\text{Year}, \text{OHC}_{lag1})$ | 869 | -10359.1 | 10.4 | 0.00 | 5362.9 |
| $e_{RS}$: non-recovering stocks | | | | | |
| $e_{RS} \sim f(\text{Year})$ | 315 | -3642.92 | 72.8 | 0.00 | 1877.8 |
| $e_{RS} \sim f(\text{Year}) + f(\text{Year}, \text{SA}_{mean})$ | 299 | -3690.02 | 25.7 | 0.00 | 1919.6 |
| $e_{RS} \sim f(\text{Year}) + f(\text{Year}, \text{SA}_{diversity})$ | 314 | -3639.29 | 76.4 | 0.00 | 1879.4 |
| $e_{RS} \sim f(\text{Year}) + f(\text{Year}, \text{SA}_{mean}) + f(\text{Year}, \text{SB})$ | 301 | -3694.86 | 20.9 | 0.00 | 1922.5 |
| $e_{RS} \sim f(\text{Year}) + f(\text{Year}, \text{SA}_{mean}) + f(\text{Year}, \text{SB}/B_{pa})$ | 303 | -3704.02 | 11.7 | 0.00 | 1923.2 |
| $e_{RS} \sim f(\text{Year}) + f(\text{Year}, \text{SA}_{mean}) + f(\text{Year}, \text{SB}/B_{pa}) + f(\text{Year}, \bar{F}_{lag1})$ | 301 | -3704.54 | 11.2 | 0.00 | 1926.3 |
| $e_{RS} \sim f(\text{Year}) + f(\text{Year}, \text{SA}_{mean}) + f(\text{Year}, \text{SB}/B_{pa}) + f(\text{Year}, \bar{F}_{lag1}/F_{msy})$ | 296 | -3707.35 | 8.4 | 0.01 | 1934.9 |
| **$e_{RS} \sim f(\text{Year}) + f(\text{Year}, \text{SA}_{mean}) + f(\text{Year}, \text{SB}/B_{pa}) + f(\text{Year}, \bar{F}_{lag1}/F_{msy}) + f(\text{Year}, \text{wSST}_{anom})$** | **300** | **-3715.73** | **0.0** | **0.90** | **1934.0** |
| $e_{RS} \sim f(\text{Year}) + f(\text{Year}, \text{SA}_{mean}) + f(\text{Year}, \text{SB}/B_{pa}) + f(\text{Year}, \bar{F}_{lag1}/F_{msy}) + f(\text{Year}, \text{spSST}_{anom})$ | 300 | -3708.26 | 7.5 | 0.02 | 1933.0 |
| $e_{RS} \sim f(\text{Year}) + f(\text{Year}, \text{SA}_{mean}) + f(\text{Year}, \text{SB}/B_{pa}) + f(\text{Year}, \bar{F}_{lag1}/F_{msy}) + f(\text{Year}, \text{smSST}_{anom})$ | 296 | -3707.42 | 8.3 | 0.01 | 1934.9 |
| $e_{RS} \sim f(\text{Year}) + f(\text{Year}, \text{SA}_{mean}) + f(\text{Year}, \text{SB}/B_{pa}) + f(\text{Year}, \bar{F}_{lag1}/F_{msy}) + f(\text{Year}, \text{aSST}_{anom})$ | 296 | -3707.14 | 8.6 | 0.01 | 1934.9 |
| $e_{RS} \sim f(\text{Year}) + f(\text{Year}, \text{SA}_{mean}) + f(\text{Year}, \text{SB}/B_{pa}) + f(\text{Year}, \bar{F}_{lag1}/F_{msy}) + f(\text{Year}, \text{NAO})$ | 296 | -3702.68 | 13.1 | 0.00 | 1934.8 |
| $e_{RS} \sim f(\text{Year}) + f(\text{Year}, \text{SA}_{mean}) + f(\text{Year}, \text{SB}/B_{pa}) + f(\text{Year}, \bar{F}_{lag1}/F_{msy}) + f(\text{Year}, \text{AMO})$ | 299 | -3695.08 | 20.6 | 0.00 | 1932.1 |
| $e_{RS} \sim f(\text{Year}) + f(\text{Year}, \text{SA}_{mean}) + f(\text{Year}, \text{SB}/B_{pa}) + f(\text{Year}, \bar{F}_{lag1}/F_{msy}) + f(\text{Year}, \text{OHC})$ | 295 | -3704.11 | 11.6 | 0.00 | 1936.7 |
| $e_{RS} \sim f(\text{Year}) + f(\text{Year}, \text{SA}_{mean}) + f(\text{Year}, \text{SB}/B_{pa}) + f(\text{Year}, \bar{F}_{lag1}/F_{msy}) + f(\text{Year}, \text{NAO}_{lag1})$ | 295 | -3702.29 | 13.4 | 0.00 | 1936.4 |
| $e_{RS} \sim f(\text{Year}) + f(\text{Year}, \text{SA}_{mean}) + f(\text{Year}, \text{SB}/B_{pa}) + f(\text{Year}, \bar{F}_{lag1}/F_{msy}) + f(\text{Year}, \text{AMO}_{lag1})$ | 296 | -3707.41 | 8.3 | 0.01 | 1934.9 |
| $e_{RS} \sim f(\text{Year}) + f(\text{Year}, \text{SA}_{mean}) + f(\text{Year}, \text{SB}/B_{pa}) + f(\text{Year}, \bar{F}_{lag1}/F_{msy}) + f(\text{Year}, \text{OHC}_{lag1})$ | 293 | -3707.57 | 8.2 | 0.02 | 1939.3 |

**Table S3**. Results of model selection for covariates of recruitment success ($RS_{total}$) as a response variable for each northeast Atlantic ecoregion in the study. The models in bold indicate the best models for response variable and recovery status combinations. DF, AIC, ΔAIC, and *w* indicate the degrees of freedom, Akaike information criterion scores, differences in AIC scores from the best models, and model weights. All time (year)-dependent smoother terms for covariates are modeled as tensor product construction (using the *t2* function from the R package *mgcv*).

| model | DF | AIC | ΔAIC | *w* | log likelihood |
|---|---|---|---|---|---|
| Baltic Sea | | | | | |
| $RS_{total}$ ~ stock + $f$(Year) | 163 | 813.8 | 17.6 | 0.00 | -397.9 |
| $RS_{total}$ ~ stock + $f$(Year) + $f$(Year, $SA_{mean}$) | 158 | 817.4 | 21.2 | 0.00 | -394.7 |
| $RS_{total}$ ~ stock + $f$(Year) + $f$(Year, $SA_{diversity}$) | 158 | 811.4 | 15.2 | 0.00 | -391.7 |
| $RS_{total}$ ~ stock + $f$(Year) + $f$(Year, $SA_{diversity}$) + $f$(Year, SB) | 152 | 817.1 | 20.9 | 0.00 | -389.5 |
| $RS_{total}$ ~ stock + $f$(Year) + $f$(Year, $SA_{diversity}$) + $f$(Year, SB/$B_{pa}$) | 152 | 799.4 | 3.2 | 0.12 | -380.7 |
| $RS_{total}$ ~ stock + $f$(Year) + $f$(Year, $SA_{diversity}$) + $f$(Year, SB/$B_{pa}$) + $f$(Year, $\bar{F}_{lag1}$) | 147 | 806.5 | 10.3 | 0.00 | -379.2 |
| $RS_{total}$ ~ stock + $f$(Year) + $f$(Year, $SA_{diversity}$) + $f$(Year, SB/$B_{pa}$) + $f$(Year, $\bar{F}_{lag1}/F_{msy}$) | 147 | 804.5 | 8.3 | 0.01 | -378.2 |
| $RS_{total}$ ~ stock + $f$(Year) + $f$(Year, $SA_{diversity}$) + $f$(Year, SB/$B_{pa}$) + $f$(Year, $wSST_{anom}$) | 147 | 806.4 | 10.2 | 0.00 | -379.2 |
| $RS_{total}$ ~ stock + $f$(Year) + $f$(Year, $SA_{diversity}$) + $f$(Year, SB/$B_{pa}$) + $f$(Year, $spSST_{anom}$) | 147 | 801.2 | 5.0 | 0.05 | -376.6 |
| **$RS_{total}$ ~ stock + $f$(Year) + $f$(Year, $SA_{diversity}$) + $f$(Year, SB/$B_{pa}$) + $f$(Year, $smSST_{anom}$)** | **148** | **796.2** | **0** | **0.58** | **-375.1** |
| $RS_{total}$ ~ stock + $f$(Year) + $f$(Year, $SA_{diversity}$) + $f$(Year, SB/$B_{pa}$) + $f$(Year, $aSST_{anom}$) | 149 | 800.0 | 3.8 | 0.09 | -378.0 |
| $RS_{total}$ ~ stock + $f$(Year) + $f$(Year, $SA_{diversity}$) + $f$(Year, SB/$B_{pa}$) + $f$(Year, NAO) | 147 | 805.2 | 9.0 | 0.01 | -378.6 |
| $RS_{total}$ ~ stock + $f$(Year) + $f$(Year, $SA_{diversity}$) + $f$(Year, SB/$B_{pa}$) + $f$(Year, AMO) | 148 | 807.4 | 11.2 | 0.00 | -380.7 |
| $RS_{total}$ ~ stock + $f$(Year) + $f$(Year, $SA_{diversity}$) + $f$(Year, SB/$B_{pa}$) + $f$(Year, OHC) | 149 | 800.1 | 3.9 | 0.08 | -378.1 |
| $RS_{total}$ ~ stock + $f$(Year) + $f$(Year, $SA_{diversity}$) + $f$(Year, SB/$B_{pa}$) + $f$(Year, $NAO_{lag1}$) | 147 | 805.5 | 9.3 | 0.01 | -378.7 |
| $RS_{total}$ ~ stock + $f$(Year) + $f$(Year, $SA_{diversity}$) + $f$(Year, SB/$B_{pa}$) + $f$(Year, $AMO_{lag1}$) | 148 | 801.5 | 5.3 | 0.04 | -377.8 |
| $RS_{total}$ ~ stock + $f$(Year) + $f$(Year, $SA_{diversity}$) + $f$(Year, SB/$B_{pa}$) + $f$(Year, $OHC_{lag1}$) | 149 | 804.1 | 7.9 | 0.01 | -380.1 |
| Bay of Biscay and Iberian Coast | | | | | |
| $RS_{total}$ ~ stock + $f$(Year) | 81 | 274.7 | 12.9 | 0.00 | -130.3 |
| $RS_{total}$ ~ stock + $f$(Year) + $f$(Year, $SA_{mean}$) | 76 | 274.7 | 12.9 | 0.00 | -125.3 |
| $RS_{total}$ ~ stock + $f$(Year) + $f$(Year, $SA_{diversity}$) | 76 | 277.0 | 15.2 | 0.00 | -126.5 |
| $RS_{total}$ ~ stock + $f$(Year) + $f$(Year, SB) | 76 | 267.8 | 6.0 | 0.04 | -121.9 |
| $RS_{total}$ ~ stock + $f$(Year) + $f$(Year, SB/$B_{pa}$) | 76 | 272.7 | 10.9 | 0.00 | -124.4 |
| $RS_{total}$ ~ stock + $f$(Year) + $f$(Year, SB) + $f$(Year, $\bar{F}_{lag1}$) | 71 | 277.4 | 15.6 | 0.00 | -121.7 |
| $RS_{total}$ ~ stock + $f$(Year) + $f$(Year, SB) + $f$(Year, $\bar{F}_{lag1}/F_{msy}$) | 71 | 273.7 | 11.9 | 0.00 | -119.9 |
| $RS_{total}$ ~ stock + $f$(Year) + $f$(Year, SB) + $f$(Year, $wSST_{anom}$) | 71 | 269.0 | 7.2 | 0.02 | -117.5 |
| $RS_{total}$ ~ stock + $f$(Year) + $f$(Year, SB) + $f$(Year, $spSST_{anom}$) | 71 | 270.3 | 8.5 | 0.01 | -118.1 |
| $RS_{total}$ ~ stock + $f$(Year) + $f$(Year, SB) + $f$(Year, $smSST_{anom}$) | 67 | 267.9 | 6.1 | 0.04 | -112.9 |
| **$RS_{total}$ ~ stock + $f$(Year) + $f$(Year, SB) + $f$(Year, $aSST_{anom}$)** | **71** | **261.8** | **0** | **0.76** | **-113.9** |
| $RS_{total}$ ~ stock + $f$(Year) + $f$(Year, SB) + $f$(Year, NAO) | 71 | 267.5 | 5.7 | 0.04 | -116.7 |
| $RS_{total}$ ~ stock + $f$(Year) + $f$(Year, SB) + $f$(Year, AMO) | 71 | 268.8 | 7.0 | 0.02 | -117.4 |
| $RS_{total}$ ~ stock + $f$(Year) + $f$(Year, SB) + $f$(Year, OHC) | 71 | 270.7 | 8.9 | 0.01 | -118.4 |
| $RS_{total}$ ~ stock + $f$(Year) + $f$(Year, SB) + $f$(Year, $NAO_{lag1}$) | 71 | 274.5 | 12.7 | 0.00 | -120.3 |
| $RS_{total}$ ~ stock + $f$(Year) + $f$(Year, SB) + $f$(Year, $AMO_{lag1}$) | 71 | 267.2 | 5.4 | 0.05 | -116.6 |
| $RS_{total}$ ~ stock + $f$(Year) + $f$(Year, SB) + $f$(Year, $OHC_{lag1}$) | 71 | 273.8 | 12.0 | 0.00 | -119.9 |
| Celtic Seas | | | | | |
| $RS_{total}$ ~ $f$(Year) | 401 | 2750.5 | 43.4 | 0.00 | -1369.2 |
| $RS_{total}$ ~ $f$(Year) + $f$(Year, $SA_{mean}$) | 396 | 2738.0 | 30.9 | 0.00 | -1358.0 |
| $RS_{total}$ ~ $f$(Year) + $f$(Year, $SA_{diversity}$) | 396 | 2751.1 | 44.0 | 0.00 | -1364.5 |
| $RS_{total}$ ~ $f$(Year) + $f$(Year, $SA_{mean}$) + $f$(Year, SB) | 391 | 2710.0 | 2.9 | 0.14 | -1339.0 |
| $RS_{total}$ ~ $f$(Year) + $f$(Year, $SA_{mean}$) + $f$(Year, SB/$B_{pa}$) | 391 | 2713.8 | 6.7 | 0.02 | -1340.9 |
| $RS_{total}$ ~ $f$(Year) + $f$(Year, $SA_{mean}$) + $f$(Year, SB) + $f$(Year, $\bar{F}_{lag1}$) | 386 | 2712.9 | 5.8 | 0.03 | -1335.4 |
| $RS_{total}$ ~ $f$(Year) + $f$(Year, $SA_{mean}$) + $f$(Year, SB) + $f$(Year, $\bar{F}_{lag1}/F_{msy}$) | 386 | 2716.1 | 9.0 | 0.01 | -1337.1 |
| $RS_{total}$ ~ $f$(Year) + $f$(Year, $SA_{mean}$) + $f$(Year, SB) + $f$(Year, $wSST_{anom}$) | 387 | 2716.4 | 9.3 | 0.01 | -1338.2 |
| $RS_{total}$ ~ $f$(Year) + $f$(Year, $SA_{mean}$) + $f$(Year, SB) + $f$(Year, $spSST_{anom}$) | 388 | 2715.2 | 8.1 | 0.01 | -1338.6 |
| $RS_{total}$ ~ $f$(Year) + $f$(Year, $SA_{mean}$) + $f$(Year, SB) + $f$(Year, $smSST_{anom}$) | 387 | 2717.0 | 9.9 | 0.00 | -1338.5 |
| $RS_{total}$ ~ $f$(Year) + $f$(Year, $SA_{mean}$) + $f$(Year, SB) + $f$(Year, $aSST_{anom}$) | 387 | 2717.4 | 10.3 | 0.00 | -1338.7 |
| **$RS_{total}$ ~ $f$(Year) + $f$(Year, $SA_{mean}$) + $f$(Year, SB) + $f$(Year, NAO)** | **386** | **2707.1** | **0** | **0.58** | **-1332.6** |
| $RS_{total}$ ~ $f$(Year) + $f$(Year, $SA_{mean}$) + $f$(Year, SB) + $f$(Year, AMO) | 387 | 2713.3 | 6.2 | 0.03 | -1336.7 |
| $RS_{total}$ ~ $f$(Year) + $f$(Year, $SA_{mean}$) + $f$(Year, SB) + $f$(Year, OHC) | 388 | 2711.9 | 4.8 | 0.05 | -1336.9 |
| $RS_{total}$ ~ $f$(Year) + $f$(Year, $SA_{mean}$) + $f$(Year, SB) + $f$(Year, $NAO_{lag1}$) | 386 | 2715.1 | 8.0 | 0.01 | -1336.6 |
| $RS_{total}$ ~ $f$(Year) + $f$(Year, $SA_{mean}$) + $f$(Year, SB) + $f$(Year, $AMO_{lag1}$) | 387 | 2715.9 | 8.8 | 0.01 | -1338.0 |
| $RS_{total}$ ~ $f$(Year) + $f$(Year, $SA_{mean}$) + $f$(Year, SB) + $f$(Year, $OHC_{lag1}$) | 388 | 2710.5 | 3.4 | 0.11 | -1336.2 |
| Faroes | | | | | |
| $RS_{total}$ ~ stock + $f$(Year) | 115 | 757.8 | 22.2 | 0.00 | -371.9 |
| $RS_{total}$ ~ stock + $f$(Year) + $f$(Year, $SA_{mean}$) | 110 | 760.3 | 24.7 | 0.00 | -368.1 |
| $RS_{total}$ ~ stock + $f$(Year) + $f$(Year, $SA_{diversity}$) | 110 | 767.7 | 32.1 | 0.00 | -371.9 |

| | | | | | |
|---|---|---|---|---|---|
| $RS_{total}$ ~ stock + *f*(Year) + *f*(Year, SB) | 110 | 750.6 | 15.0 | 0.00 | -363.3 |
| $RS_{total}$ ~ stock + *f*(Year) + *f*(Year, SB/$B_{pa}$) | 110 | 738.6 | 3.0 | 0.12 | -357.3 |
| $RS_{total}$ ~ stock + *f*(Year) + *f*(Year, SB/$B_{pa}$) + *f*(Year, $\bar{F}_{lag1}$) | 105 | 742.2 | 6.6 | 0.02 | -354.1 |
| $RS_{total}$ ~ stock + *f*(Year) + *f*(Year, SB/$B_{pa}$) + *f*(Year, $\bar{F}_{lag1}/F_{msy}$) | 105 | 747.6 | 12.0 | 0.00 | -356.8 |
| $RS_{total}$ ~ stock + *f*(Year) + *f*(Year, SB/$B_{pa}$) + *f*(Year, $wSST_{anom}$) | 105 | 742.6 | 7.0 | 0.02 | -354.3 |
| $RS_{total}$ ~ stock + *f*(Year) + *f*(Year, SB/$B_{pa}$) + *f*(Year, $spSST_{anom}$) | 105 | 741.7 | 6.1 | 0.03 | -353.9 |
| $RS_{total}$ ~ stock + *f*(Year) + *f*(Year, SB/$B_{pa}$) + *f*(Year, $smSST_{anom}$) | 105 | 743.6 | 8.0 | 0.01 | -354.8 |
| $RS_{total}$ ~ stock + *f*(Year) + *f*(Year, SB/$B_{pa}$) + *f*(Year, $aSST_{anom}$) | 105 | 745.6 | 10.0 | 0.00 | -355.8 |
| **$RS_{total}$ ~ stock + *f*(Year) + *f*(Year, SB/$B_{pa}$) + *f*(Year, NAO)** | **105** | **735.6** | **0** | **0.54** | **-350.8** |
| $RS_{total}$ ~ stock + *f*(Year) + *f*(Year, SB/$B_{pa}$) + *f*(Year, AMO) | 105 | 746.5 | 10.9 | 0.00 | -356.2 |
| $RS_{total}$ ~ stock + *f*(Year) + *f*(Year, SB/$B_{pa}$) + *f*(Year, OHC) | 106 | 742.0 | 6.4 | 0.02 | -355.0 |
| $RS_{total}$ ~ stock + *f*(Year) + *f*(Year, SB/$B_{pa}$) + *f*(Year, $NAO_{lag1}$) | 105 | 745.8 | 10.2 | 0.00 | -355.9 |
| $RS_{total}$ ~ stock + *f*(Year) + *f*(Year, SB/$B_{pa}$) + *f*(Year, $AMO_{lag1}$) | 105 | 737.5 | 1.9 | 0.21 | -351.8 |
| $RS_{total}$ ~ stock + *f*(Year) + *f*(Year, SB/$B_{pa}$) + *f*(Year, $OHC_{lag1}$) | 106 | 742.5 | 6.9 | 0.02 | -355.2 |
| Icelandic waters | | | | | |
| $RS_{total}$ ~ stock + *f*(Year) | 144 | 836.5 | 29.5 | 0.00 | -410.3 |
| $RS_{total}$ ~ stock + *f*(Year) + *f*(Year, $SA_{mean}$) | 139 | 827.1 | 20.1 | 0.00 | -400.5 |
| $RS_{total}$ ~ stock + *f*(Year) + *f*(Year, $SA_{diversity}$) | 139 | 839.0 | 32.0 | 0.00 | -406.5 |
| $RS_{total}$ ~ stock + *f*(Year) + *f*(Year, $SA_{mean}$) + *f*(Year, SB) | 134 | 820.1 | 13.1 | 0.00 | -392.1 |
| $RS_{total}$ ~ stock + *f*(Year) + *f*(Year, $SA_{mean}$) + *f*(Year, SB/$B_{pa}$) | 134 | 808.4 | 1.4 | 0.25 | -386.2 |
| $RS_{total}$ ~ stock + *f*(Year) + *f*(Year, $SA_{mean}$) + *f*(Year, SB/$B_{pa}$) + *f*(Year, $\bar{F}_{lag1}$) | 129 | 813.5 | 6.5 | 0.02 | -383.8 |
| $RS_{total}$ ~ stock + *f*(Year) + *f*(Year, $SA_{mean}$) + *f*(Year, SB/$B_{pa}$) + *f*(Year, $\bar{F}_{lag1}/F_{msy}$) | 129 | 813.1 | 6.1 | 0.02 | -383.6 |
| $RS_{total}$ ~ stock + *f*(Year) + *f*(Year, $SA_{mean}$) + *f*(Year, SB/$B_{pa}$) + *f*(Year, $wSST_{anom}$) | 129 | 817.6 | 10.6 | 0.00 | -385.8 |
| **$RS_{total}$ ~ stock + *f*(Year) + *f*(Year, $SA_{mean}$) + *f*(Year, SB/$B_{pa}$) + *f*(Year, $spSST_{anom}$)** | **129** | **807.0** | **0** | **0.50** | **-380.5** |
| $RS_{total}$ ~ stock + *f*(Year) + *f*(Year, $SA_{mean}$) + *f*(Year, SB/$B_{pa}$) + *f*(Year, $smSST_{anom}$) | 129 | 820.2 | 13.2 | 0.00 | -387.1 |
| $RS_{total}$ ~ stock + *f*(Year) + *f*(Year, $SA_{mean}$) + *f*(Year, SB/$B_{pa}$) + *f*(Year, $aSST_{anom}$) | 129 | 820.1 | 13.1 | 0.00 | -387.0 |
| $RS_{total}$ ~ stock + *f*(Year) + *f*(Year, $SA_{mean}$) + *f*(Year, SB/$B_{pa}$) + *f*(Year, NAO) | 129 | 816.6 | 9.6 | 0.00 | -385.3 |
| $RS_{total}$ ~ stock + *f*(Year) + *f*(Year, $SA_{mean}$) + *f*(Year, SB/$B_{pa}$) + *f*(Year, AMO) | 129 | 820.1 | 13.1 | 0.00 | -387.1 |
| $RS_{total}$ ~ stock + *f*(Year) + *f*(Year, $SA_{mean}$) + *f*(Year, SB/$B_{pa}$) + *f*(Year, OHC) | 129 | 819.7 | 12.7 | 0.00 | -386.9 |
| $RS_{total}$ ~ stock + *f*(Year) + *f*(Year, $SA_{mean}$) + *f*(Year, SB/$B_{pa}$) + *f*(Year, $NAO_{lag1}$) | 129 | 819.1 | 12.1 | 0.00 | -386.6 |
| $RS_{total}$ ~ stock + *f*(Year) + *f*(Year, $SA_{mean}$) + *f*(Year, SB/$B_{pa}$) + *f*(Year, $AMO_{lag1}$) | 129 | 808.9 | 1.9 | 0.19 | -381.4 |
| $RS_{total}$ ~ stock + *f*(Year) + *f*(Year, $SA_{mean}$) + *f*(Year, SB/$B_{pa}$) + *f*(Year, $OHC_{lag1}$) | 129 | 819.1 | 12.1 | 0.00 | -386.5 |
| North Sea | | | | | |
| $RS_{total}$ ~ stock + *f*(Year) | 282 | 2251.5 | 62.3 | 0.00 | -1114.8 |
| $RS_{total}$ ~ stock + *f*(Year) + *f*(Year, $SA_{mean}$) | 277 | 2217.0 | 27.8 | 0.00 | -1092.5 |
| $RS_{total}$ ~ stock + *f*(Year) + *f*(Year, $SA_{diversity}$) | 277 | 2237.3 | 48.1 | 0.00 | -1102.7 |
| $RS_{total}$ ~ stock + *f*(Year) + *f*(Year, $SA_{mean}$) + *f*(Year, SB) | 272 | 2192.7 | 3.5 | 0.09 | -1075.3 |
| $RS_{total}$ ~ stock + *f*(Year) + *f*(Year, $SA_{mean}$) + *f*(Year, SB/$B_{pa}$) | 272 | 2190.7 | 1.5 | 0.24 | -1074.4 |
| **$RS_{total}$ ~ stock + *f*(Year) + *f*(Year, $SA_{mean}$) + *f*(Year, SB/$B_{pa}$) + *f*(Year, $\bar{F}_{lag1}$)** | **267** | **2189.2** | **0** | **0.50** | **-1068.6** |
| $RS_{total}$ ~ stock + *f*(Year) + *f*(Year, $SA_{mean}$) + *f*(Year, SB/$B_{pa}$) + *f*(Year, $\bar{F}_{lag1}/F_{msy}$) | 267 | 2193.7 | 4.5 | 0.05 | -1070.8 |
| $RS_{total}$ ~ stock + *f*(Year) + *f*(Year, $SA_{mean}$) + *f*(Year, SB/$B_{pa}$) + *f*(Year, $wSST_{anom}$) | 267 | 2193.3 | 4.1 | 0.06 | -1070.7 |
| $RS_{total}$ ~ stock + *f*(Year) + *f*(Year, $SA_{mean}$) + *f*(Year, SB/$B_{pa}$) + *f*(Year, $spSST_{anom}$) | 267 | 2197.2 | 8.0 | 0.01 | -1072.6 |
| $RS_{total}$ ~ stock + *f*(Year) + *f*(Year, $SA_{mean}$) + *f*(Year, SB/$B_{pa}$) + *f*(Year, $smSST_{anom}$) | 267 | 2200.6 | 11.4 | 0.00 | -1074.3 |
| $RS_{total}$ ~ stock + *f*(Year) + *f*(Year, $SA_{mean}$) + *f*(Year, SB/$B_{pa}$) + *f*(Year, $aSST_{anom}$) | 267 | 2196.7 | 7.5 | 0.01 | -1072.4 |
| $RS_{total}$ ~ stock + *f*(Year) + *f*(Year, $SA_{mean}$) + *f*(Year, SB/$B_{pa}$) + *f*(Year, NAO) | 267 | 2198.6 | 9.4 | 0.00 | -1073.3 |
| $RS_{total}$ ~ stock + *f*(Year) + *f*(Year, $SA_{mean}$) + *f*(Year, SB/$B_{pa}$) + *f*(Year, AMO) | 267 | 2197.4 | 8.2 | 0.01 | -1072.7 |
| $RS_{total}$ ~ stock + *f*(Year) + *f*(Year, $SA_{mean}$) + *f*(Year, SB/$B_{pa}$) + *f*(Year, OHC) | 268 | 2196.4 | 7.2 | 0.01 | -1073.2 |
| $RS_{total}$ ~ stock + *f*(Year) + *f*(Year, $SA_{mean}$) + *f*(Year, SB/$B_{pa}$) + *f*(Year, $NAO_{lag1}$) | 267 | 2199.3 | 10.1 | 0.00 | -1073.6 |
| $RS_{total}$ ~ stock + *f*(Year) + *f*(Year, $SA_{mean}$) + *f*(Year, SB/$B_{pa}$) + *f*(Year, $AMO_{lag1}$) | 267 | 2198.8 | 9.6 | 0.00 | -1073.4 |
| $RS_{total}$ ~ stock + *f*(Year) + *f*(Year, $SA_{mean}$) + *f*(Year, SB/$B_{pa}$) + *f*(Year, $OHC_{lag1}$) | 268 | 2198.1 | 8.9 | 0.01 | -1074.1 |
| Norwegian–Barents Seas | | | | | |
| $RS_{total}$ ~ stock + *f*(Year) | 135 | 1208.0 | 81.9 | 0.00 | -597.0 |
| $RS_{total}$ ~ stock + *f*(Year) + *f*(Year, $SA_{mean}$) | 130 | 1180.7 | 54.6 | 0.00 | -578.3 |
| $RS_{total}$ ~ stock + *f*(Year) + *f*(Year, $SA_{diversity}$) | 130 | 1184.9 | 58.8 | 0.00 | -580.4 |
| $RS_{total}$ ~ stock + *f*(Year) + *f*(Year, $SA_{mean}$) + *f*(Year, SB) | 125 | 1128.5 | 2.4 | 0.12 | -547.2 |
| $RS_{total}$ ~ stock + *f*(Year) + *f*(Year, $SA_{mean}$) + *f*(Year, SB/$B_{pa}$) | 125 | 1151.0 | 24.9 | 0.00 | -558.5 |
| $RS_{total}$ ~ stock + *f*(Year) + *f*(Year, $SA_{mean}$) + *f*(Year, SB) + *f*(Year, $\bar{F}_{lag1}$) | 120 | 1129.7 | 3.6 | 0.07 | -542.8 |
| $RS_{total}$ ~ stock + *f*(Year) + *f*(Year, $SA_{mean}$) + *f*(Year, SB) + *f*(Year, $\bar{F}_{lag1}/F_{msy}$) | 120 | 1130.2 | 4.1 | 0.05 | -543.1 |
| $RS_{total}$ ~ stock + *f*(Year) + *f*(Year, $SA_{mean}$) + *f*(Year, SB) + *f*(Year, $wSST_{anom}$) | 120 | 1127.1 | 1.0 | 0.25 | -541.5 |
| **$RS_{total}$ ~ stock + *f*(Year) + *f*(Year, $SA_{mean}$) + *f*(Year, SB) + *f*(Year, $spSST_{anom}$)** | **120** | **1126.1** | **0** | **0.41** | **-541.0** |
| $RS_{total}$ ~ stock + *f*(Year) + *f*(Year, $SA_{mean}$) + *f*(Year, SB) + *f*(Year, $smSST_{anom}$) | 120 | 1132.7 | 6.6 | 0.02 | -544.3 |
| $RS_{total}$ ~ stock + *f*(Year) + *f*(Year, $SA_{mean}$) + *f*(Year, SB) + *f*(Year, $aSST_{anom}$) | 120 | 1130.5 | 4.4 | 0.05 | -543.3 |
| $RS_{total}$ ~ stock + *f*(Year) + *f*(Year, $SA_{mean}$) + *f*(Year, SB) + *f*(Year, NAO) | 120 | 1135.8 | 9.7 | 0.00 | -545.9 |
| $RS_{total}$ ~ stock + *f*(Year) + *f*(Year, $SA_{mean}$) + *f*(Year, SB) + *f*(Year, AMO) | 120 | 1138.4 | 12.3 | 0.00 | -547.2 |
| $RS_{total}$ ~ stock + *f*(Year) + *f*(Year, $SA_{mean}$) + *f*(Year, SB) + *f*(Year, OHC) | 120 | 1133.8 | 7.7 | 0.01 | -544.9 |
| $RS_{total}$ ~ stock + *f*(Year) + *f*(Year, $SA_{mean}$) + *f*(Year, SB) + *f*(Year, $NAO_{lag1}$) | 120 | 1134.4 | 8.3 | 0.01 | -545.2 |
| $RS_{total}$ ~ stock + *f*(Year) + *f*(Year, $SA_{mean}$) + *f*(Year, SB) + *f*(Year, $AMO_{lag1}$) | 120 | 1138.1 | 12.0 | 0.00 | -547.0 |
| $RS_{total}$ ~ stock + *f*(Year) + *f*(Year, $SA_{mean}$) + *f*(Year, SB) + *f*(Year, $OHC_{lag1}$) | 120 | 1135.1 | 9.0 | 0.00 | -545.6 |

**Table S4**. Parameter estimates and test statistics for the generalized additive mixed effects models with recruitment success ($RS_{total}$) as a response variable with covariates selected (Table S3) for each northeast Atlantic ecoregion in the study. EDF, *F*, CI, *t*, and SD indicate the effective degrees of freedom, F-statistic, confidence intervals, t-value, and standard deviation. All time (year)-dependent smoother terms for covariates are modeled as tensor product construction (using the *t2* function from the R package *mgcv*).

| Model structure and parameter | smoother | | fixed effect | | | random effect | |
|---|---|---|---|---|---|---|---|
| | EDF | F | estimate | 95% CI | t | variance | SD |
| Baltic Sea | | | | | | | |
| $RS_{total}$ ~ stock + $f$(Year) + $f$(Year, $SA_{diversity}$) + $f$(Year, SB/$B_{pa}$) + $f$(Year, $smSST_{anom}$) | | | | | | | |
| $f$(Year) | 1.00 | 14.87 | | | | | |
| $f$(Year, $SA_{diversity}$) | 3.19 | 0.73 | | | | | |
| $f$(Year, SB/$B_{pa}$) | 5.34 | 2.92 | | | | | |
| $f$(Year, $smSST_{anom}$) | 2.87 | 6.19 | | | | | |
| $\beta_{cod}$ | | | 1.77 | (1.25, 2.29) | 6.67 | | |
| $\beta_{herring}$ | | | 1.23 | (0.94, 1.51) | 8.44 | | |
| $\beta_{sole}$ | | | 1.05 | (0.74, 1.37) | 6.52 | | |
| $\beta_{sprat}$ | | | 1.71 | (1.50, 1.93) | 15.8 | | |
| $\beta_{Gulf\ of\ Riga\ herring}$ | | | 1.51 | (1.23, 1.79) | 10.6 | | |
| $\beta_{Year}$ | | | -0.33 | (-0.54, -0.12) | -3.05 | | |
| $\beta_{Year×SAdiversity1}$ | | | 0.13 | (-0.03, 0.29) | 1.60 | | |
| $\beta_{Year×SAdiversity2}$ | | | 0.06 | (-0.08, 0.2) | 0.83 | | |
| $\beta_{Year×SB/Bpa1}$ | | | -0.29 | (-0.45, -0.13) | -3.47 | | |
| $\beta_{Year×SB/Bpa2}$ | | | 0.00 | (-0.19, 0.18) | -0.05 | | |
| $\beta_{Year×smSSTanom}$ | | | 0.29 | (0.10, 0.48) | 3.01 | | |
| year | | | | | | 0.06 | 0.24 |
| residual | | | | | | 0.24 | 0.49 |
| Bay of Biscay and the Coast | | | | | | | |
| $RS_{total}$ ~ stock + $f$(Year) + $f$(Year, SB) + $f$(Year, $aSST_{anom}$) | | | | | | | |
| $f$(Year) | 1.00 | 2.03 | | | | | |
| $f$(Year, SB) | 5.56 | 3.37 | | | | | |
| $f$(Year, $fSST_{anom}$) | 3.12 | 2.95 | | | | | |
| $\beta_{four-spot\ megrim}$ | | | 1.33 | (1.1, 1.56) | 11.3 | | |
| $\beta_{megrim}$ | | | 0.42 | (-0.13, 0.97) | 1.50 | | |
| $\beta_{sole}$ | | | 2.06 | (1.43, 2.69) | 6.40 | | |
| $\beta_{Year}$ | | | -0.03 | (-0.16, 0.1) | -0.50 | | |
| $\beta_{Year×SB/Bpa1}$ | | | 0.75 | (0.29, 1.2) | 3.20 | | |
| $\beta_{Year×SB/Bpa2}$ | | | 0.04 | (-0.04, 0.12) | 0.88 | | |
| $\beta_{Year×aSSTanom1}$ | | | -0.11 | (-0.21, -0.02) | -2.40 | | |
| $\beta_{Year×aSSTanom2}$ | | | -0.16 | (-0.26, -0.07) | -3.29 | | |
| year | | | | | | 0.03 | 0.16 |
| residual | | | | | | 0.06 | 0.25 |
| Celtic Seas | | | | | | | |
| $RS_{total}$ ~ $f$(Year) + $f$(Year, $SA_{mean}$) + $f$(Year, SB) + $f$(Year, NAO) | | | | | | | |
| $f$(Year) | 1.00 | 3.64 | | | | | |
| $f$(Year, $SA_{mean}$) | 5.83 | 3.12 | | | | | |
| $f$(Year, SB) | 5.21 | 2.71 | | | | | |
| $f$(Year, NAO) | 6.04 | 1.29 | | | | | |
| $\beta_0$ | | | 2.47 | (1.85, 3.09) | 7.83 | | |
| $\beta_{Year}$ | | | -0.13 | (-0.35, 0.10.) | -1.11 | | |
| $\beta_{Year×SAmean1}$ | | | -0.40 | (-0.66, -0.13) | -2.92 | | |
| $\beta_{Year×SAmean2}$ | | | 0.05 | (-0.07, 0.17) | 0.84 | | |
| $\beta_{Year×SB1}$ | | | -0.28 | (-0.46, -0.11) | -3.16 | | |
| $\beta_{Year×SB2}$ | | | -0.07 | (-0.21, 0.08) | -0.87 | | |
| $\beta_{Year×NAO1}$ | | | -0.02 | (-0.18, 0.13) | -0.30 | | |
| $\beta_{Year×NAO2}$ | | | 0.10 | (-0.04, 0.25) | 1.38 | | |

| | | | | | | | |
|---|---|---|---|---|---|---|---|
| year | | | | | | 0.03 | 0.18 |
| stock | | | | | | 0.39 | 0.62 |
| residual | | | | | | 0.38 | 0.62 |
| Faroes | | | | | | | |
| $RS_{total}$ ~ stock + $f$(Year) + $f$(Year, SB/$B_{pa}$) + $f$(Year, NAO) | | | | | | | |
| $f$(Year) | 1.00 | 0.01 | | | | | |
| $f$(Year, SB/$B_{pa}$) | 7.51 | 2.49 | | | | | |
| $f$(Year, NAO) | 2.01 | 7.27 | | | | | |
| $\beta_{cod}$ | | | 1.84 | (1.83, 1.84) | 685.5 | | |
| $\beta_{haddock}$ | | | 2.01 | (2.00, 2.02) | 749.5 | | |
| $\beta_{saithe}$ | | | 1.95 | (1.95, 1.96) | 729.2 | | |
| $\beta_{Year}$ | | | -0.01 | (-0.01, 0.00) | -2.13 | | |
| $\beta_{Year \times SB/Bpa1}$ | | | 0.17 | (0.17, 0.18) | 63.7 | | |
| $\beta_{Year \times SB/Bpa2}$ | | | 0.32 | (0.32, 0.33) | 119.8 | | |
| $\beta_{Year \times NAO1}$ | | | -0.27 | (-0.28, -0.27) | -102.1 | | |
| $\beta_{Year \times NAO2}$ | | | 0.05 | (0.04, 0.06) | 18.7 | | |
| year | | | | | | 0.14 | 0.37 |
| residual | | | | | | 0.41 | 0.64 |
| Icelandic waters | | | | | | | |
| $RS_{total}$ ~ stock + $f$(Year) + $f$(Year, $SA_{mean}$) + $f$(Year, SB/$B_{pa}$) + $f$(Year, $spSST_{anom}$) | | | | | | | |
| $f$(Year) | 1.00 | 3.44 | | | | | |
| $f$(Year, $SA_{mean}$) | 7.04 | 0.40 | | | | | |
| $f$(Year, SB/$B_{pa}$) | 4.70 | 3.35 | | | | | |
| $f$(Year, $spSST_{anom}$) | 6.11 | 1.40 | | | | | |
| $\beta_{ccod}$ | | | 2.78 | (2.78, 2.78) | 1922.8 | | |
| $\beta_{chaddock}$ | | | 1.39 | (1.38, 1.39) | 958.7 | | |
| $\beta_{cherring}$ | | | 1.38 | (1.38, 1.38) | 955.0 | | |
| $\beta_{csaithe}$ | | | 1.76 | (1.76, 1.76) | 1218.0 | | |
| $\beta_{Year}$ | | | -0.22 | (-0.22, -0.22) | -152.1 | | |
| $\beta_{Year \times SAmean1}$ | | | -0.12 | (-0.12, -0.11) | -80.7 | | |
| $\beta_{Year \times SAmean2}$ | | | 0.00 | (0.00, 0.01) | 2.3 | | |
| $\beta_{Year \times SB/Bpa1}$ | | | 0.33 | (0.33, 0.34) | 230.3 | | |
| $\beta_{Year \times SB/Bpa2}$ | | | -0.12 | (-0.12, -0.12) | -84.2 | | |
| $\beta_{Year \times spSSTanom1}$ | | | 0.21 | (0.21, 0.22) | 147.1 | | |
| $\beta_{Year \times spSSTanom2}$ | | | 0.08 | (0.08, 0.08) | 56.1 | | |
| year | | | | | | 0.09 | 0.29 |
| residual | | | | | | 0.15 | 0.39 |
| North Sea | | | | | | | |
| $RS_{total}$ ~ stock + $f$(Year) + $f$(Year, $SA_{mean}$) + $f$(Year, SB/$B_{pa}$) + $f$(Year, $\bar{F}_{lag1}$) | | | | | | | |
| $f$(Year) | 1.00 | 46.16 | | | | | |
| $f$(Year, $SA_{mean}$) | 3.05 | 5.24 | | | | | |
| $f$(Year, SB/$B_{pa}$) | 2.00 | 23.88 | | | | | |
| $f$(Year, $\bar{F}_{lag1}$) | 2.00 | 8.16 | | | | | |
| $\beta_{plaice\ (D\ 6.d)}$ | | | 1.87 | (1.57, 2.18) | 11.99 | | |
| $\beta_{cod}$ | | | 2.95 | (2.71, 3.18) | 24.34 | | |
| $\beta_{haddock}$ | | | 4.73 | (4.49, 4.97) | 38.87 | | |
| $\beta_{plaice\ (SA\ 4,\ SD\ 20)}$ | | | 2.11 | (1.88, 2.34) | 18.13 | | |
| $\beta_{saithe}$ | | | 2.87 | (2.54, 3.2) | 17.06 | | |
| $\beta_{turbot}$ | | | 3.15 | (2.79, 3.51) | 17.11 | | |
| $\beta_{whiting}$ | | | 2.67 | (2.37, 2.97) | 17.56 | | |
| $\beta_{Year}$ | | | -0.40 | (-0.53, -0.27) | -5.98 | | |
| $\beta_{Year \times SAmean1}$ | | | 0.42 | (0.22, 0.63) | 4.01 | | |
| $\beta_{Year \times SAmean2}$ | | | 0.01 | (-0.15, 0.16) | 0.09 | | |
| $\beta_{Year \times SB/Bpa1}$ | | | -0.48 | (-0.62, -0.34) | -6.72 | | |
| $\beta_{Year \times SB/Bpa2}$ | | | 0.05 | (-0.07, 0.18) | 0.87 | | |
| $\beta_{Year \times \bar{F}lag1.1}$ | | | -0.04 | (-0.23, 0.15) | -0.39 | | |
| $\beta_{Year \times \bar{F}lag1.2}$ | | | 0.31 | (0.15, 0.47) | 3.75 | | |
| year | | | | | | 0.04 | 0.20 |
| residual | | | | | | 0.32 | 0.57 |

| | | | | | | | |
|---|---|---|---|---|---|---|---|
| Norwegian–Barents Seas<br>$RS_{total}$ ~ stock + $f$(Year) + $f$(Year, $SA_{mean}$) +<br>$f$(Year, SB) + $f$(Year, $spSST_{anom}$) | | | | | | | |
| $f$(Year) | 1.05 | 4.63 | | | | | |
| $f$(Year, $SA_{mean}$) | 7.27 | 2.37 | | | | | |
| $f$(Year, SB) | 6.41 | 7.86 | | | | | |
| $f$(Year, $spSST_{anom}$) | 2.65 | 16.49 | | | | | |
| $\beta_{cod}$ | | | 4.38 | (4.38, 4.38) | 3987.6 | | |
| $\beta_{haddock}$ | | | 2.41 | (2.41, 2.42) | 2196.5 | | |
| $\beta_{saithe}$ | | | 2.79 | (2.79, 2.79) | 2538.8 | | |
| $\beta_{Year}$ | | | 0.29 | (0.29, 0.29) | 265.7 | | |
| $\beta_{Year \times SAmean1}$ | | | 0.39 | (0.38, 0.39) | 351.4 | | |
| $\beta_{Year \times SAmean2}$ | | | -0.11 | (-0.12, -0.11) | -103.6 | | |
| $\beta_{Year \times SB1}$ | | | 2.25 | (2.25, 2.25) | 2047.3 | | |
| $\beta_{Year \times SB2}$ | | | 0.92 | (0.92, 0.92) | 838.0 | | |
| $\beta_{Year \times spSSTanom1}$ | | | -0.46 | (-0.46, -0.46) | -417.1 | | |
| $\beta_{Year \times spSSTanom2}$ | | | -0.35 | (-0.36, -0.35) | -321.5 | | |
| year | | | | | | 0.21 | 0.45 |
| residual | | | | | | 0.33 | 0.58 |

**Table S5**. Results of model selection for covariates of elasticity to recruitment success ($e_{RS}$) as a response variable for each northeast Atlantic ecoregion in the study. The models in bold indicate the best models for response variable and recovery status combinations. DF, AIC, ΔAIC, and *w* indicate the degrees of freedom, Akaike information criterion scores, differences in AIC scores from the best models, and model weights. All time (year)-dependent smoother terms for covariates are modeled as tensor product construction (using the *te* function from the R package *mgcv*).

| model | DF | AIC | ΔAIC | *w* | log-likelihood |
|---|---|---|---|---|---|
| Baltic Sea | | | | | |
| $e_{RS}$ ~ stock + *f*(Year) | 146 | -1725.6 | 13.1 | 0.00 | 892.8 |
| $e_{RS}$ ~ stock + *f*(Year) + *f*(Year, $SA_{mean}$) | 144 | -1728.5 | 10.2 | 0.00 | 895.7 |
| $e_{RS}$ ~ stock + *f*(Year) + *f*(Year, $SA_{diversity}$) | 143 | -1732.1 | 6.65 | 0.02 | 899.6 |
| $e_{RS}$ ~ stock + *f*(Year) + *f*(Year, $SA_{diversity}$) + *f*(Year, SB) | 141 | -1725.2 | 13.6 | 0.00 | 897.1 |
| $e_{RS}$ ~ stock + *f*(Year) + *f*(Year, $SA_{diversity}$) + *f*(Year, SB/$B_{pa}$) | 134 | -1728.6 | 10.1 | 0.00 | 908.4 |
| $e_{RS}$ ~ stock + *f*(Year) + *f*(Year, $SA_{diversity}$) + *f*(Year, $\bar{F}_{lag1}$) | 143 | -1732 | 6.67 | 0.02 | 899.6 |
| $e_{RS}$ ~ stock + *f*(Year) + *f*(Year, $SA_{diversity}$) + *f*(Year, $\bar{F}_{lag1}/F_{msy}$) | 143 | -1732.1 | 6.66 | 0.02 | 899.6 |
| $e_{RS}$ ~ stock + *f*(Year) + *f*(Year, $SA_{diversity}$) + *f*(Year, $wSST_{anom}$) | 141 | -1735 | 3.68 | 0.11 | 903.2 |
| $e_{RS}$ ~ stock + *f*(Year) + *f*(Year, $SA_{diversity}$) + *f*(Year, $spSST_{anom}$) | 137 | -1723.4 | 15.3 | 0.00 | 906.7 |
| $e_{RS}$ ~ stock + *f*(Year) + *f*(Year, $SA_{diversity}$) + *f*(Year, $smSST_{anom}$) | 144 | -1731.8 | 6.91 | 0.02 | 902.6 |
| $e_{RS}$ ~ stock + *f*(Year) + *f*(Year, $SA_{diversity}$) + *f*(Year, $aSST_{anom}$) | 141 | -1731.4 | 7.3 | 0.02 | 903.5 |
| $e_{RS}$ ~ stock + *f*(Year) + *f*(Year, $SA_{diversity}$) + *f*(Year, NAO) | 149 | -1730.1 | 8.65 | 0.01 | 894.7 |
| $e_{RS}$ ~ stock + *f*(Year) + *f*(Year, $SA_{diversity}$) + (Year, AMO) | 143 | -1731.2 | 7.5 | 0.02 | 899.2 |
| **$e_{RS}$ ~ stock + *f*(Year) + *f*(Year, $SA_{diversity}$) + *f*(Year, OHC)** | **139** | **-1738.7** | **0** | **0.68** | **907.0** |
| $e_{RS}$ ~ stock + *f*(Year) + *f*(Year, $SA_{diversity}$) + *f*(Year, $NAO_{lag1}$) | 142 | -1727.8 | 10.9 | 0.00 | 900.2 |
| $e_{RS}$ ~ stock + *f*(Year) + *f*(Year, $SA_{diversity}$) + *f*(Year, $AMO_{lag1}$) | 139 | -1732.6 | 6.09 | 0.03 | 906.2 |
| $e_{RS}$ ~ stock + *f*(Year) + *f*(Year, $SA_{diversity}$) + *f*(Year, $OHC_{lag1}$) | 143 | -1732 | 6.67 | 0.02 | 899.6 |
| Bay of Biscay and Iberian Coast | | | | | |
| $e_{RS}$ ~ stock + *f*(Year) | 76 | -942.89 | 14.5 | 0.00 | 488.5 |
| $e_{RS}$ ~ stock + *f*(Year) + *f*(Year, $SA_{mean}$) | 74 | -949.06 | 8.3 | 0.00 | 495.2 |
| $e_{RS}$ ~ stock + *f*(Year) + *f*(Year, $SA_{diversity}$) | 76 | -942.86 | 14.5 | 0.00 | 488.5 |
| $e_{RS}$ ~ stock + *f*(Year) + *f*(Year, $SA_{mean}$) + *f*(Year, SB) | 73 | -947.96 | 9.4 | 0.00 | 496.9 |
| $e_{RS}$ ~ stock + *f*(Year) + *f*(Year, $SA_{mean}$) + *f*(Year, SB/$B_{pa}$) | 76 | -952.88 | 4.48 | 0.03 | 494.5 |
| $e_{RS}$ ~ stock + *f*(Year) + *f*(Year, $SA_{mean}$) + *f*(Year, SB/$B_{pa}$) + *f*(Year, $\bar{F}_{lag1}$) | 74 | -952.01 | 5.35 | 0.02 | 495.9 |
| $e_{RS}$ ~ stock + *f*(Year) + *f*(Year, $SA_{mean}$) + *f*(Year, SB/$B_{pa}$) + *f*(Year, $\bar{F}_{lag1}/F_{msy}$) | 70 | -952.83 | 4.53 | 0.03 | 500.9 |
| $e_{RS}$ ~ stock + *f*(Year) + *f*(Year, $SA_{mean}$) + *f*(Year, SB/$B_{pa}$) + *f*(Year, $wSST_{anom}$) | 64 | -955.41 | 1.95 | 0.11 | 507.4 |
| $e_{RS}$ ~ stock + *f*(Year) + *f*(Year, $SA_{mean}$) + *f*(Year, SB/$B_{pa}$) + *f*(Year, $spSST_{anom}$) | 75 | -956.55 | 0.81 | 0.20 | 494.5 |
| $e_{RS}$ ~ stock + *f*(Year) + *f*(Year, $SA_{mean}$) + *f*(Year, SB/$B_{pa}$) + *f*(Year, $smSST_{anom}$) | 66 | -955.05 | 2.31 | 0.10 | 505.8 |
| $e_{RS}$ ~ stock + *f*(Year) + *f*(Year, $SA_{mean}$) + *f*(Year, SB/$B_{pa}$) + *f*(Year, $aSST_{anom}$) | 64 | -950.67 | 6.69 | 0.01 | 504.7 |
| $e_{RS}$ ~ stock + *f*(Year) + *f*(Year, $SA_{mean}$) + *f*(Year, SB/$B_{pa}$) + *f*(Year, NAO) | 74 | -953.2 | 4.16 | 0.04 | 495.7 |
| **$e_{RS}$ ~ stock + *f*(Year) + *f*(Year, $SA_{mean}$) + *f*(Year, SB/$B_{pa}$) + (Year, AMO)** | **77** | **-957.36** | **0** | **0.30** | **494.8** |
| $e_{RS}$ ~ stock + *f*(Year) + *f*(Year, $SA_{mean}$) + *f*(Year, SB/$B_{pa}$) + *f*(Year, OHC) | 69 | -955.7 | 1.66 | 0.13 | 502.7 |
| $e_{RS}$ ~ stock + *f*(Year) + *f*(Year, $SA_{mean}$) + *f*(Year, SB/$B_{pa}$) + *f*(Year, $NAO_{lag1}$) | 69 | -947.53 | 9.83 | 0.00 | 500.8 |
| $e_{RS}$ ~ stock + *f*(Year) + *f*(Year, $SA_{mean}$) + *f*(Year, SB/$B_{pa}$) + *f*(Year, $AMO_{lag1}$) | 68 | -949.49 | 7.87 | 0.01 | 502.3 |
| $e_{RS}$ ~ stock + *f*(Year) + *f*(Year, $SA_{mean}$) + *f*(Year, SB/$B_{pa}$) + *f*(Year, $OHC_{lag1}$) | 71 | -949.36 | 8 | 0.01 | 497.6 |
| Celtic Seas | | | | | |
| $e_{RS}$ ~ *f*(Year) | 388 | -3996.5 | 55.5 | 0.00 | 2024.9 |
| $e_{RS}$ ~ *f*(Year) + *f*(Year, $SA_{mean}$) | 382 | -4013.7 | 38.4 | 0.00 | 2037.1 |
| $e_{RS}$ ~ *f*(Year) + *f*(Year, $SA_{diversity}$) | 388 | -4007.8 | 44.2 | 0.00 | 2029.4 |
| $e_{RS}$ ~ *f*(Year) + *f*(Year, $SA_{mean}$) + *f*(Year, SB) | 378 | -4040.9 | 11.1 | 0.00 | 2057.3 |
| $e_{RS}$ ~ *f*(Year) + *f*(Year, $SA_{mean}$) + *f*(Year, SB/$B_{pa}$) | 381 | -4029.8 | 22.2 | 0.00 | 2049.1 |
| $e_{RS}$ ~ *f*(Year) + *f*(Year, $SA_{mean}$) + *f*(Year, SB/$B_{pa}$)+ *f*(Year, $\bar{F}_{lag1}$) | 373 | -4048.4 | 3.65 | 0.02 | 2067.9 |
| **$e_{RS}$ ~ *f*(Year) + *f*(Year, $SA_{mean}$) + *f*(Year, SB/$B_{pa}$) + *f*(Year, $\bar{F}_{lag1}/F_{msy}$)** | **373** | **-4052** | **0** | **0.13** | **2067.7** |
| $e_{RS}$ ~ *f*(Year) + *f*(Year, $SA_{mean}$) + *f*(Year, SB/$B_{pa}$) + *f*(Year, $wSST_{anom}$) | 374 | -4052 | 0.02 | 0.13 | 2066.3 |
| $e_{RS}$ ~ *f*(Year) + *f*(Year, $SA_{mean}$) + *f*(Year, SB/$B_{pa}$) + *f*(Year, $spSST_{anom}$) | 375 | -4048.9 | 3.13 | 0.03 | 2067.0 |
| $e_{RS}$ ~ *f*(Year) + *f*(Year, $SA_{mean}$) + *f*(Year, SB/$B_{pa}$) + *f*(Year, $smSST_{anom}$) | 374 | -4052 | 0.02 | 0.13 | 2066.3 |
| $e_{RS}$ ~ *f*(Year) + *f*(Year, $SA_{mean}$) + *f*(Year, SB/$B_{pa}$) + *f*(Year, $aSST_{anom}$) | 374 | -4052 | 0.04 | 0.12 | 2066.3 |
| $e_{RS}$ ~ *f*(Year) + *f*(Year, $SA_{mean}$) + *f*(Year, SB/$B_{pa}$) + *f*(Year, NAO) | 374 | -4052 | 0.02 | 0.13 | 2066.3 |
| $e_{RS}$ ~ *f*(Year) + *f*(Year, $SA_{mean}$) + *f*(Year, SB/$B_{pa}$) + (Year, AMO) | 375 | -4046.4 | 5.62 | 0.01 | 2065.8 |

| | | | | | |
|---|---|---|---|---|---|
| $e_{RS}$ ~ $f$(Year) + $f$(Year, $SA_{mean}$) + $f$(Year, $SB/B_{pa}$) + $f$(Year, OHC) | 374 | -4052 | 0.04 | 0.12 | 2066.3 |
| $e_{RS}$ ~ $f$(Year) + $f$(Year, $SA_{mean}$) + $f$(Year, $SB/B_{pa}$) + $f$(Year, $NAO_{lag1}$) | 376 | -4046.5 | 5.53 | 0.01 | 2065.2 |
| $e_{RS}$ ~ $f$(Year) + $f$(Year, $SA_{mean}$) + $f$(Year, $SB/B_{pa}$) + $f$(Year, $AMO_{lag1}$) | 374 | -4052 | 0.05 | 0.12 | 2066.3 |
| $e_{RS}$ ~ $f$(Year) + $f$(Year, $SA_{mean}$) + $f$(Year, $SB/B_{pa}$) + $f$(Year, $OHC_{lag1}$) | 375 | -4050.6 | 1.43 | 0.06 | 2066.1 |
| Faroes | | | | | |
| $e_{RS}$ ~ stock + $f$(Year) | 104 | -1375.6 | 73.6 | 0.00 | 729.8 |
| $e_{RS}$ ~ stock + $f$(Year) + $f$(Year, $SA_{mean}$) | 98 | -1393.4 | 55.8 | 0.00 | 745.0 |
| $e_{RS}$ ~ stock + $f$(Year) + $f$(Year, $SA_{diversity}$) | 101 | -1379.3 | 70 | 0.00 | 735.6 |
| $e_{RS}$ ~ stock + $f$(Year) + $f$(Year, $SA_{mean}$) + $f$(Year, SB) | 87 | -1410.6 | 38.6 | 0.00 | 766.7 |
| $e_{RS}$ ~ stock + $f$(Year) + $f$(Year, $SA_{mean}$) + $f$(Year, $SB/B_{pa}$) | 84 | -1424.7 | 24.5 | 0.00 | 776.6 |
| $e_{RS}$ ~ stock + $f$(Year) + $f$(Year, $SA_{mean}$) + $f$(Year, SB) + $f$(Year, $\bar{F}_{lag1}$) | 105 | -1224 | 225 | 0.00 | 656.6 |
| $e_{RS}$ ~ stock + $f$(Year) + $f$(Year, $SA_{mean}$) + $f$(Year, SB) + $f$(Year, $\bar{F}_{lag1}/F_{msy}$) | 70 | -1444.7 | 4.53 | 0.08 | 801.7 |
| $e_{RS}$ ~ stock + $f$(Year) + $f$(Year, $SA_{mean}$) + $f$(Year, SB) + $f$(Year, $\bar{F}_{lag1}/F_{msy}$) + $f$(Year, $wSST_{anom}$) | 132 | -831.9 | 617 | 0.00 | 428.8 |
| $e_{RS}$ ~ stock + $f$(Year) + $f$(Year, $SA_{mean}$) + $f$(Year, SB) + $f$(Year, $\bar{F}_{lag1}/F_{msy}$) + $f$(Year, $spSST_{anom}$) | 131 | -830.72 | 619 | 0.00 | 429.2 |
| $e_{RS}$ ~ stock + $f$(Year) + $f$(Year, $SA_{mean}$) + $f$(Year, SB) + $f$(Year, $\bar{F}_{lag1}/F_{msy}$) + $f$(Year, $smSST_{anom}$) | 131 | -909.96 | 539 | 0.00 | 469.2 |
| $e_{RS}$ ~ stock + $f$(Year) + $f$(Year, $SA_{mean}$) + $f$(Year, SB) + $f$(Year, $\bar{F}_{lag1}/F_{msy}$) + $f$(Year, $aSST_{anom}$) | 126 | -998.96 | 450 | 0.00 | 522.5 |
| **$e_{RS}$ ~ stock + $f$(Year) + $f$(Year, $SA_{mean}$) + $f$(Year, SB)** + $f$(Year, $\bar{F}_{lag1}/F_{msy}$) **+ $f$(Year, NAO)** | **70** | **-1449.3** | **0** | **0.75** | **805.0** |
| $e_{RS}$ ~ stock + $f$(Year) + $f$(Year, $SA_{mean}$) + $f$(Year, SB) + $f$(Year, $\bar{F}_{lag1}/F_{msy}$) + (Year, AMO) | 72 | -1440.4 | 8.9 | 0.01 | 799.1 |
| $e_{RS}$ ~ stock + $f$(Year) + $f$(Year, $SA_{mean}$) + $f$(Year, SB) + $f$(Year, $\bar{F}_{lag1}/F_{msy}$) + $f$(Year, OHC) | 72 | -1444 | 5.22 | 0.06 | 799.4 |
| $e_{RS}$ ~ stock + $f$(Year) + $f$(Year, $SA_{mean}$) + $f$(Year, SB) + $f$(Year, $\bar{F}_{lag1}/F_{msy}$) + $f$(Year, $NAO_{lag1}$) | 96 | -1167.7 | 282 | 0.00 | 638.1 |
| $e_{RS}$ ~ stock + $f$(Year) + $f$(Year, $SA_{mean}$) + $f$(Year, SB) + $f$(Year, $\bar{F}_{lag1}/F_{msy}$) + $f$(Year, $AMO_{lag1}$) | 133 | -757.8 | 691 | 0.00 | 391.8 |
| $e_{RS}$ ~ stock + $f$(Year) + $f$(Year, $SA_{mean}$) + $f$(Year, SB) + $f$(Year, $\bar{F}_{lag1}/F_{msy}$) + $f$(Year, $OHC_{lag1}$) | 69 | -1445.3 | 3.94 | 0.10 | 803.8 |
| Icelandic waters | | | | | |
| $e_{RS}$ ~ stock + $f$(Year) | 125 | -1503 | 33.2 | 0.00 | 781.4 |
| $e_{RS}$ ~ stock + $f$(Year) + $f$(Year, $SA_{mean}$) | 117 | -1507.1 | 29.1 | 0.00 | 794.1 |
| $e_{RS}$ ~ stock + $f$(Year) + $f$(Year, $SA_{diversity}$) | 125 | -1503 | 33.2 | 0.00 | 781.4 |
| $e_{RS}$ ~ stock + $f$(Year) + $f$(Year, $SA_{mean}$) + $f$(Year, SB) | 110 | -1511.4 | 24.8 | 0.00 | 804.2 |
| $e_{RS}$ ~ stock + $f$(Year) + $f$(Year, $SA_{mean}$) + $f$(Year, SB) + $f$(Year, $SB/B_{pa}$) | 108 | -1524.6 | 11.6 | 0.00 | 812.6 |
| $e_{RS}$ ~ stock + $f$(Year) + $f$(Year, $SA_{mean}$) + $f$(Year, SB) + $f$(Year, $\bar{F}_{lag1}$) | 106 | -1519.6 | 16.6 | 0.00 | 817.1 |
| $e_{RS}$ ~ stock + $f$(Year) + $f$(Year, $SA_{mean}$) + $f$(Year, SB) + $f$(Year, $\bar{F}_{lag1}/F_{msy}$) | 105 | -1521.4 | 14.8 | 0.00 | 817.9 |
| $e_{RS}$ ~ stock + $f$(Year) + $f$(Year, $SA_{mean}$) + $f$(Year, SB) + $f$(Year, $wSST_{anom}$) | 102 | -1523.2 | 13 | 0.00 | 821.0 |
| **$e_{RS}$ ~ stock + $f$(Year)** + $f$(Year, $SA_{mean}$) + $f$(Year, SB) **+ $f$(Year, $spSST_{anom}$)** | **106** | **-1536.2** | **0** | **0.96** | **821.7** |
| $e_{RS}$ ~ stock + $f$(Year) + $f$(Year, $SA_{mean}$) + $f$(Year, SB) + $f$(Year, $smSST_{anom}$) | 109 | -1524.2 | 12 | 0.00 | 813.4 |
| $e_{RS}$ ~ stock + $f$(Year) + $f$(Year, $SA_{mean}$) + $f$(Year, SB) + $f$(Year, $aSST_{anom}$) | 109 | -1524.3 | 11.9 | 0.00 | 813.3 |
| $e_{RS}$ ~ stock + $f$(Year) + $f$(Year, $SA_{mean}$) + $f$(Year, SB) + $f$(Year, NAO) | 109 | -1524 | 12.2 | 0.00 | 813.4 |
| $e_{RS}$ ~ stock + $f$(Year) + $f$(Year, $SA_{mean}$) + $f$(Year, SB) + (Year, AMO) | 109 | -1519.9 | 16.3 | 0.00 | 813.7 |
| $e_{RS}$ ~ stock + $f$(Year) + $f$(Year, $SA_{mean}$) + $f$(Year, SB) + $f$(Year, OHC) | 107 | -1525.2 | 11 | 0.00 | 815.4 |
| $e_{RS}$ ~ stock + $f$(Year) + $f$(Year, $SA_{mean}$) + $f$(Year, SB) + $f$(Year, $NAO_{lag1}$) | 109 | -1524.3 | 11.9 | 0.00 | 812.8 |
| $e_{RS}$ ~ stock + $f$(Year) + $f$(Year, $SA_{mean}$) + $f$(Year, SB) + $f$(Year, $AMO_{lag1}$) | 106 | -1528.7 | 7.47 | 0.02 | 819.1 |
| $e_{RS}$ ~ stock + $f$(Year) + $f$(Year, $SA_{mean}$) + $f$(Year, SB) + $f$(Year, $OHC_{lag1}$) | 107 | -1517.5 | 18.7 | 0.00 | 814.6 |
| North Sea | | | | | |
| $e_{RS}$ ~ stock + $f$(Year) | 284 | -2941.3 | 65.5 | 0.00 | 1481.7 |
| $e_{RS}$ ~ stock + $f$(Year) + $f$(Year, $SA_{mean}$) | 280 | -2956.2 | 50.6 | 0.00 | 1495.3 |
| $e_{RS}$ ~ stock + $f$(Year) + $f$(Year, $SA_{diversity}$) | 281 | -2952.4 | 54.4 | 0.00 | 1491.6 |
| $e_{RS}$ ~ stock + $f$(Year) + $f$(Year, $SA_{mean}$) + $f$(Year, SB) | 278 | -2998.7 | 8.09 | 0.02 | 1517.4 |
| $e_{RS}$ ~ stock + $f$(Year) + $f$(Year, $SA_{mean}$) + $f$(Year, $SB/B_{pa}$) | 264 | -2972.9 | 33.8 | 0.00 | 1525.2 |
| $e_{RS}$ ~ stock + $f$(Year) + $f$(Year, $SA_{mean}$) + $f$(Year, $SB/B_{pa}$) + $f$(Year, $\bar{F}_{lag1}$) | 277 | -2995.9 | 10.9 | 0.00 | 1518.7 |
| **$e_{RS}$ ~ stock + $f$(Year) + $f$(Year, $SA_{mean}$) + $f$(Year, $SB/B_{pa}$) + $f$(Year, $\bar{F}_{lag1}/F_{msy}$)** | **270** | **-3006.8** | **0** | **0.87** | **1532.3** |
| $e_{RS}$ ~ stock + $f$(Year) + $f$(Year, $SA_{mean}$) + $f$(Year, $SB/B_{pa}$) + $f$(Year, $wSST_{anom}$) | 274 | -2998.9 | 7.88 | 0.02 | 1523.5 |
| $e_{RS}$ ~ stock + $f$(Year) + $f$(Year, $SA_{mean}$) + $f$(Year, $SB/B_{pa}$) + $f$(Year, $spSST_{anom}$) | 273 | -2999.1 | 7.69 | 0.02 | 1525.1 |
| $e_{RS}$ ~ stock + $f$(Year) + $f$(Year, $SA_{mean}$) + $f$(Year, $SB/B_{pa}$) + $f$(Year, $smSST_{anom}$) | 274 | -2998.8 | 7.95 | 0.02 | 1523.5 |
| $e_{RS}$ ~ stock + $f$(Year) + $f$(Year, $SA_{mean}$) + $f$(Year, $SB/B_{pa}$) + $f$(Year, $aSST_{anom}$) | 272 | -2996.5 | 10.3 | 0.01 | 1525.5 |
| $e_{RS}$ ~ stock + $f$(Year) + $f$(Year, $SA_{mean}$) + $f$(Year, $SB/B_{pa}$) + $f$(Year, NAO) | 269 | -2995.6 | 11.2 | 0.00 | 1530.4 |
| $e_{RS}$ ~ stock + $f$(Year) + $f$(Year, $SA_{mean}$) + $f$(Year, $SB/B_{pa}$) + (Year, AMO) | 271 | -2996.5 | 10.2 | 0.01 | 1527.5 |
| $e_{RS}$ ~ stock + $f$(Year) + $f$(Year, $SA_{mean}$) + $f$(Year, $SB/B_{pa}$) + $f$(Year, OHC) | 274 | -2999 | 7.77 | 0.02 | 1523.4 |
| $e_{RS}$ ~ stock + $f$(Year) + $f$(Year, $SA_{mean}$) + $f$(Year, $SB/B_{pa}$) + $f$(Year, $NAO_{lag1}$) | 269 | -2995.5 | 11.2 | 0.00 | 1529.9 |
| $e_{RS}$ ~ stock + $f$(Year) + $f$(Year, $SA_{mean}$) + $f$(Year, $SB/B_{pa}$) + $f$(Year, $AMO_{lag1}$) | 274 | -2999.1 | 7.72 | 0.02 | 1523.4 |
| $e_{RS}$ ~ stock + $f$(Year) + $f$(Year, $SA_{mean}$) + $f$(Year, $SB/B_{pa}$) + $f$(Year, $OHC_{lag1}$) | 272 | -2995.8 | 11 | 0.00 | 1524.7 |
| Norwegian–Barents Seas | | | | | |

| | | | | | |
|---|---|---|---|---|---|
| $e_{RS}$ ~ stock + *f*(Year) | 122 | -1475.3 | 64.8 | 0.00 | 776.4 |
| $e_{RS}$ ~ stock + *f*(Year) + *f*(Year, $SA_{mean}$) | 119 | -1484.2 | 55.9 | 0.00 | 786.3 |
| $e_{RS}$ ~ stock + *f*(Year) + *f*(Year, $SA_{diversity}$) | 114 | -1479 | 61.1 | 0.00 | 787.7 |
| $e_{RS}$ ~ stock + *f*(Year) + *f*(Year, $SA_{mean}$) + *f*(Year, SB) | 105 | -1521.4 | 18.6 | 0.00 | 820.5 |
| $e_{RS}$ ~ stock + *f*(Year) + *f*(Year, $SA_{mean}$) + *f*(Year, SB/$B_{pa}$) | 114 | -1500.7 | 39.4 | 0.00 | 798.2 |
| $e_{RS}$ ~ stock + *f*(Year) + *f*(Year, $SA_{mean}$) + *f*(Year, SB) + *f*(Year, $\bar{F}_{lag1}$) | 100 | -1535.6 | 4.51 | 0.09 | 831.8 |
| **$e_{RS}$ ~ stock + *f*(Year) + *f*(Year, $SA_{mean}$) + *f*(Year, SB) + *f*(Year, $\bar{F}_{lag1}/F_{msy}$)** | **103** | **-1540.1** | **0** | **0.84** | **829.3** |
| $e_{RS}$ ~ stock + *f*(Year) + *f*(Year, $SA_{mean}$) + *f*(Year, SB) + *f*(Year, $wSST_{anom}$) | 105 | -1525.4 | 14.7 | 0.00 | 825.3 |
| $e_{RS}$ ~ stock + *f*(Year) + *f*(Year, $SA_{mean}$) + *f*(Year, SB) + *f*(Year, $spSST_{anom}$) | 105 | -1516.1 | 24 | 0.00 | 825.9 |
| $e_{RS}$ ~ stock + *f*(Year) + *f*(Year, $SA_{mean}$) + *f*(Year, SB) + *f*(Year, $smSST_{anom}$) | 105 | -1531 | 9.11 | 0.01 | 825.8 |
| $e_{RS}$ ~ stock + *f*(Year) + *f*(Year, $SA_{mean}$) + *f*(Year, SB) + *f*(Year, $aSST_{anom}$) | 105 | -1531.6 | 8.5 | 0.01 | 825.5 |
| $e_{RS}$ ~ stock + *f*(Year) + *f*(Year, $SA_{mean}$) + *f*(Year, SB) + *f*(Year, NAO) | 105 | -1532.6 | 7.54 | 0.02 | 825.4 |
| $e_{RS}$ ~ stock + *f*(Year) + *f*(Year, $SA_{mean}$) + *f*(Year, SB) + (Year, AMO) | 105 | -1532.3 | 7.8 | 0.02 | 825.1 |
| $e_{RS}$ ~ stock + *f*(Year) + *f*(Year, $SA_{mean}$) + *f*(Year, SB) + *f*(Year, OHC) | 106 | -1527 | 13.1 | 0.00 | 826.7 |
| $e_{RS}$ ~ stock + *f*(Year) + *f*(Year, $SA_{mean}$) + *f*(Year, SB) + *f*(Year, $NAO_{lag1}$) | 103 | -1514.8 | 25.3 | 0.00 | 827.5 |
| $e_{RS}$ ~ stock + *f*(Year) + *f*(Year, $SA_{mean}$) + *f*(Year, SB) + *f*(Year, $AMO_{lag1}$) | 105 | -1532.3 | 7.77 | 0.02 | 825.1 |
| $e_{RS}$ ~ stock + *f*(Year) + *f*(Year, $SA_{mean}$) + *f*(Year, SB) + *f*(Year, $OHC_{lag1}$) | 104 | -1523.3 | 16.8 | 0.00 | 826.8 |

**Table S6**. Parameter estimates and test statistics for the generalized additive mixed effects models with elasticity to recruitment success ($e_{RS}$) as a response variable with covariates selected (Table S5) for each northeast Atlantic ecoregion in the study. EDF, *F*, 95% CI, *t*, and SD indicate the effective degrees of freedom, F-statistic, confidence intervals, t-value, and standard deviation. All time (year)-dependent smoother terms for covariates are modeled as tensor product construction (using the te function in the R package *mgcv*).

| model structure and parameter | smoother | | fixed effect | | | random effect | |
|---|---|---|---|---|---|---|---|
| | EDF | F | estimate | 95% CI | t | variance | SD |
| Baltic Sea | | | | | | | |
| $e_{RS}$ ~ stock + $f$(Year) + $f$(Year, $SA_{diversity}$) + $f$(Year, OHC) | | | | | | | |
| $f$(Year) | 2.31 | 1.42 | | | | | |
| $f$(Year, $SA_{diversity}$) | 2.04 | 0.60 | | | | | |
| $f$(Year, OHC) | | | | | | | |
| $\beta_{cod}$ | 2.00 | 0.35 | | | | | |
| $\beta_{herring\ (SD\ 25\text{-}29\ and\ 32)}$ | | | -5.45 | (-5.56, -5.54) | -74.6 | | |
| $\beta_{sole}$ | | | -5.23 | (-5.34, -5.32) | -98.4 | | |
| $\beta_{sprat}$ | | | -5.26 | (-5.35, -5.33) | -93.8 | | |
| year | | | | | | 0.02 | 0.15 |
| Bay of Biscay and the Coast | | | | | | | |
| $e_{RS}$ ~ stock + $f$(Year) + $f$(Year, $SA_{mean}$) + $f$(Year, SB/$B_{pa}$) + $f$(Year, AMO) | | | | | | | |
| $f$(Year) | 1.00 | 5.11 | | | | | |
| $f$(Year, $SA_{mean}$) | 0.91 | 0.33 | | | | | |
| $f$(Year, SB/$B_{pa}$) | 2.63 | 1.98 | | | | | |
| $f$(Year, AMO) | 1.69 | 2.47 | | | | | |
| $\beta_{four\text{-}spot\ megrim}$ | | | -5.19 | (-5.27, -5.11) | -126.0 | | |
| $\beta_{megrim}$ | | | -5.44 | (-7.47, -3.4) | -130.0 | | |
| $\beta_{sole}$ | | | -5.40 | (-9.4, -1.4) | -138.0 | | |
| year | | | | | | 0.01 | 0.07 |
| Celtic Seas | | | | | | | |
| $e_{RS}$ ~ $f$(Year) + $f$(Year, $SA_{mean}$) + $f$(Year, SB/$B_{pa}$) + $f$(Year, $\bar{F}_{lag1}/F_{msy}$) | | | | | | | |
| $f$(Year) | 1.00 | 0.02 | | | | | |
| $f$(Year, $SA_{mean}$) | 3.29 | 6.99 | | | | | |
| $f$(Year, SB/$B_{pa}$) | 5.65 | 7.39 | | | | | |
| $f$(Year, $\bar{F}_{lag1}/F_{msy}$) | 6.00 | 2.61 | | | | | |
| $\beta_0$ | | | -5.16 | (-5.24, -5.07) | -104.0 | | |
| $\sigma_{27.6.a.\_whiting}$ | | | -0.15 | (-0.29, -0.01) | | | |
| $\sigma_{6.d.\_cod}$ | | | -0.08 | (-0.24, 0.09) | | | |
| $\sigma_{7.a.\_whiting}$ | | | 0.05 | (-0.08, 0.19) | | | |
| $\sigma_{7.e.\_sole}$ | | | 0.04 | (-0.12, 0.2) | | | |
| $\sigma_{7.f.g.\_sole}$ | | | 0.00 | (-0.15, 0.15) | | | |
| $\sigma_{cod}$ | | | 0.00 | (-0.13, 0.14) | | | |
| $\sigma_{haddock}$ | | | 0.18 | (0.01, 0.35) | | | |
| $\sigma_{Irish\ Sea\_cod}$ | | | 0.20 | (0.07, 0.33) | | | |
| $\sigma_{Irish\ Sea\_plaice}$ | | | 0.12 | (-0.01, 0.25) | | | |
| $\sigma_{Irish\ Sea\_sole}$ | | | -0.27 | (-0.43, -0.11) | | | |
| $\sigma_{Rockall\_haddock}$ | | | -0.10 | (-0.25, 0.05) | | | |
| stock | | | | | | 0.02 | 0.16 |
| year | | | | | | 0.00 | 0.05 |
| Faroes | | | | | | | |
| $e_{RS}$ ~ stock + $f$(Year) + $f$(Year, $SA_{mean}$) + $f$(Year, SB) + $f$(Year, $\bar{F}_{lag1}/F_{msy}$) + $f$(Year, NAO) | | | | | | | |
| $f$(Year) | 1.00 | 8.20 | | | | | |
| $f$(Year, $SA_{mean}$) | 1.84 | 1.22 | | | | | |
| $f$(Year, SB) | 2.05 | 6.71 | | | | | |
| $f$(Year, $\bar{F}_{lag1}/F_{msy}$) | 1.00 | 0.32 | | | | | |
| $f$(Year, NAO) | 0.00 | 0.00 | | | | | |

| | | | | | | | |
|---|---|---|---|---|---|---|---|
| $\beta_{\text{cod}}$ | | | -5.1602 | (-5.25, -5.07) | -110.0 | | |
| $\beta_{\text{haddock}}$ | | | -5.27 | (-5.37, -5.17) | -103.5 | | |
| $\beta_{\text{saithe}}$ | | | -4.99 | (-5.15, -4.82) | -60.7 | | |
| year | | | | | | 0.02 | 0.14 |
| Icelandic waters | | | | | | | |
| $e_{\text{RS}}$ ~ stock + $f$(Year) + $f$(Year, $\text{SA}_{\text{mean}}$) + $f$(Year, SB) + $f$(Year, $\text{spSST}_{\text{anom}}$) | | | | | | | |
| $f$(Year) | 1.00 | 9.45 | | | | | |
| $f$(Year, $\text{SA}_{\text{mean}}$) | 4.62 | 2.55 | | | | | |
| $f$(Year, SB) | 6.53 | 3.32 | | | | | |
| $f$(Year, $\text{spSST}_{\text{anom}}$) | 2.87 | 7.94 | | | | | |
| $\beta_{\text{cod}}$ | | | -4.95 | (-5.4, -5.26) | -136.0 | | |
| $\beta_{\text{haddock}}$ | | | -5.33 | (-5.44, -5.24) | -99.7 | | |
| $\beta_{\text{herring}}$ | | | -5.34 | (-5.22, -5.02) | -107.0 | | |
| $\beta_{\text{saithe}}$ | | | -5.12 | (-0.1, 0.1) | -99.9 | | |
| year | | | | | | 0.02 | 0.13 |
| North Sea | | | | | | | |
| $e_{\text{RS}}$ ~ stock + $f$(Year) + $f$(Year, $\text{SA}_{\text{mean}}$) + $f$(Year, SB/$B_{\text{pa}}$) + $f$(Year, $\bar{F}_{\text{lag1}}/F_{\text{msy}}$) | | | | | | | |
| $f$(Year) | 1.00 | 10.37 | | | | | |
| $f$(Year, $\text{SA}_{\text{mean}}$) | 0.87 | 0.14 | | | | | |
| $f$(Year, SB/$B_{\text{pa}}$) | 4.97 | 4.09 | | | | | |
| $f$(Year, $\bar{F}_{\text{lag1}}/F_{\text{msy}}$) | 0.00 | 0.00 | | | | | |
| $\beta_{\text{plaice (D 6.d)}}$ | | | -5.1847 | (-5.33, -5.04) | -71.5 | | |
| $\beta_{\text{cod}}$ | | | -5.15 | (-5.23, -5.08) | -136.0 | | |
| $\beta_{\text{haddock}}$ | | | -4.92 | (-5, -4.85) | -126.8 | | |
| $\beta_{\text{plaice (SA 4, SD 20)}}$ | | | -4.88 | (-4.97, -4.79) | -107.9 | | |
| $\beta_{\text{saithe}}$ | | | -5.05 | (-5.14, -4.97) | -116.3 | | |
| $\beta_{\text{turbot}}$ | | | -5.11 | (-5.27, -4.94) | -60.5 | | |
| $\beta_{\text{whiting}}$ | | | -5.24 | (-5.38, -5.11) | -78.4 | | |
| year | | | | | | 0.00 | 0.06 |
| Norwegian–Barents Seas | | | | | | | |
| $e_{\text{RS}}$ ~ stock + $f$(Year) + $f$(Year, $\text{SA}_{\text{mean}}$) + $f$(Year, SB) + $f$(Year, $\bar{F}_{\text{lag1}}/F_{\text{msy}}$) | | | | | | | |
| $f$(Year) | 1.00 | 0.47 | | | | | |
| $f$(Year, $\text{SA}_{\text{mean}}$) | 4.82 | 11.26 | | | | | |
| $f$(Year, SB) | 8.83 | 8.53 | | | | | |
| $f$(Year, $\bar{F}_{\text{lag1}}/F_{\text{msy}}$) | 3.25 | 4.32 | | | | | |
| $\beta_{\text{cod}}$ | | | -4.85 | (-4.95, -4.75) | -95.6 | | |
| $\beta_{\text{haddock}}$ | | | -5.13 | (-5.22, -5.05) | -116.7 | | |
| $\beta_{\text{saithe}}$ | | | -4.93 | (-5.02, -4.84) | -108.6 | | |
| year | | | | | | 0.03 | 0.16 |

**Figure S1**. Temporal variations in generation time estimates of 38 northeast Atlantic fish stocks during 1946–2016.

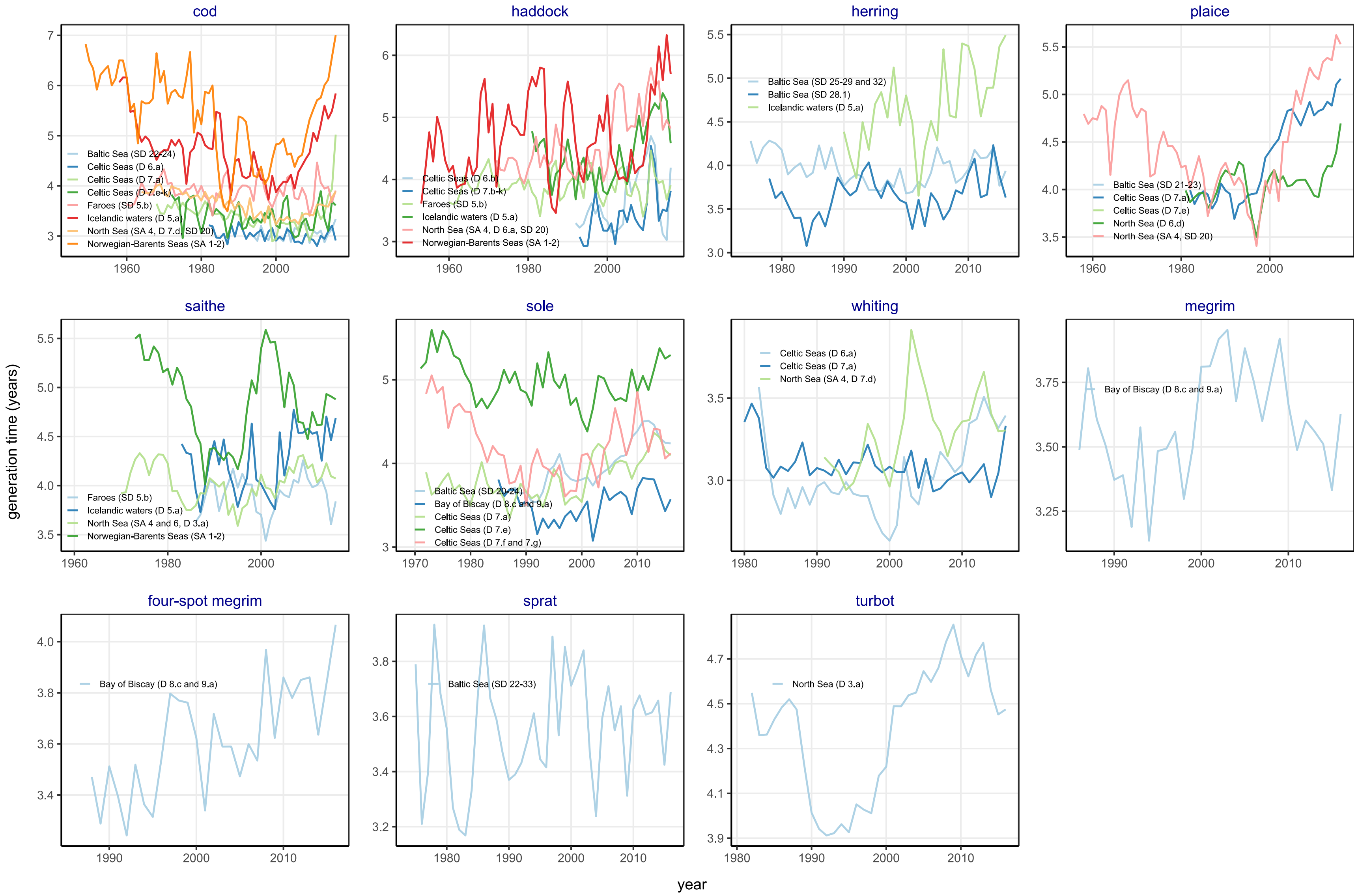

**Figure S2**. Sensitivity analysis for the classification of 38 northeast Atlantic fish stocks based on recovery status. A threshold for recovery status, relative spawner biomass, SB/$B_{pa}$, is tested for the range between 0.6 and 1.1. Lines indicate smoothed trends (with 95% confidence intervals) estimated using generalized additive mixed effects models with year as a fixed effect and stock, ecoregion, and year nested in ecoregion as random effects. Vertical dashed lines indicate the reference year (2002) used to define the recovery status of fish stocks.

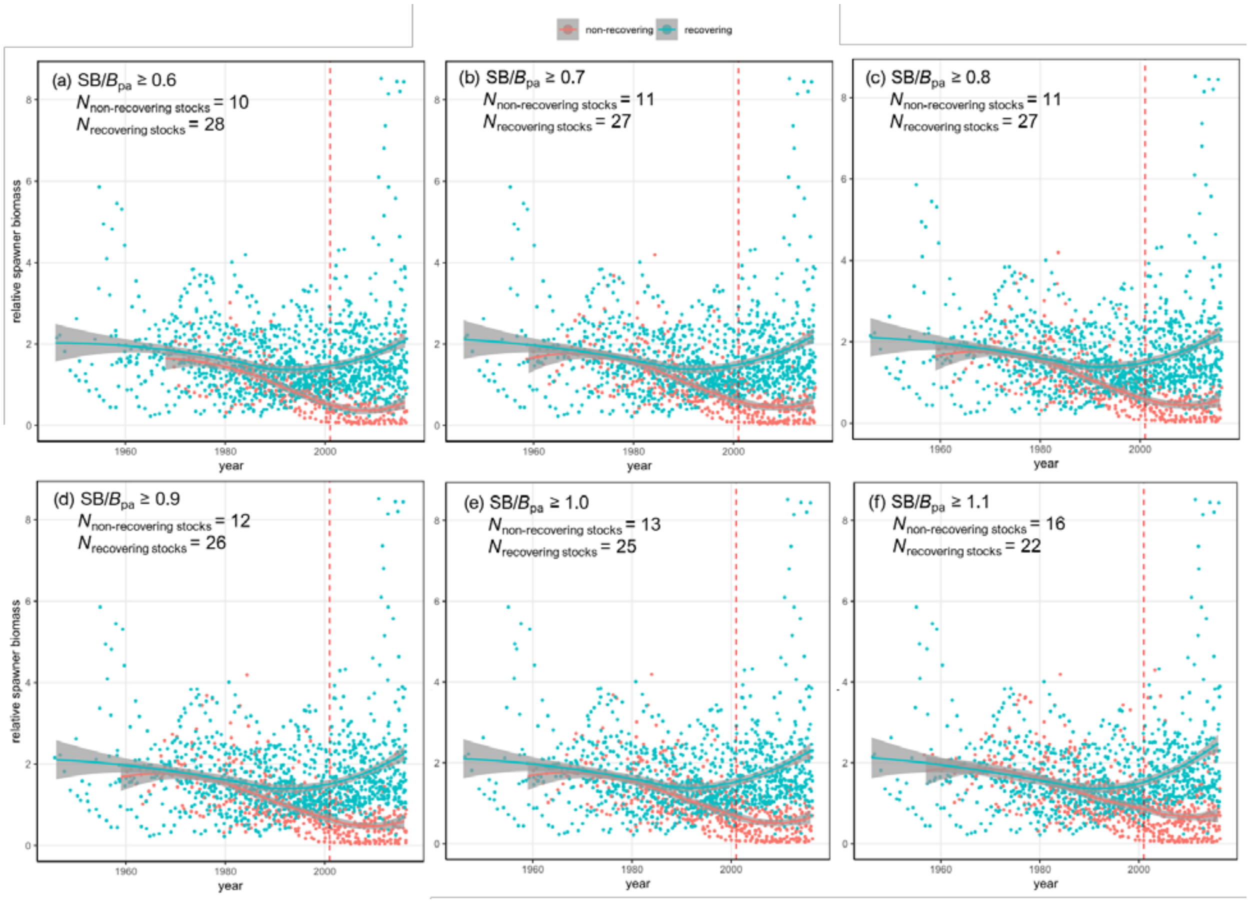

**Figure S3**. Sensitivity analysis for the classification of 38 northeast Atlantic fish stocks based on the frequency of meeting the recovery threshold (SB/$B_{pa}$, = 0.8). The minimum frequency (in proportion) of a stock meeting or exceeding a threshold for recovery (relative spawner biomass, SB/$B_{pa}$, ≥ 0.8) is tested for the range between 50% and 100% during the last 15 years of the time series of data (2002–2016). Lines indicate smoothed trends (with 95% confidence intervals) estimated using generalized additive mixed effects models with year as a fixed effect and stock, ecoregion, and year nested in ecoregion as random effects. Vertical dashed lines indicate the reference year (2002) used to define the recovery status of fish stocks.

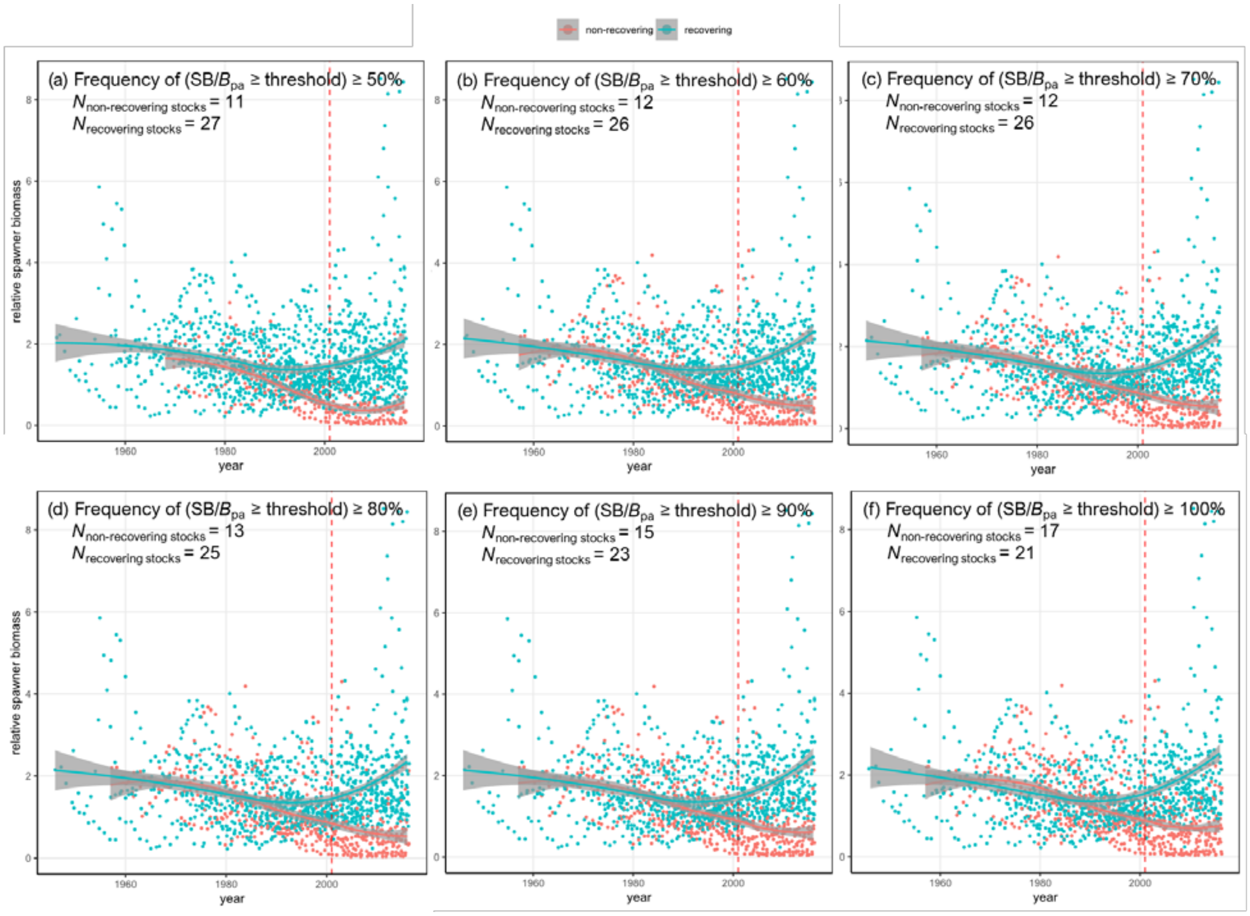

**Figure S4**. Time series of the transient population growths of 36 northeast Atlantic fish stocks during 1946–2016. Red lines indicate transient growths computed from age-specific abundance and demographic rate estimates taken from the International Council for the Exploration of the Sea (ICES) 2017 stock assessment reports. Blue lines indicate transient growths computed from 1000 simulated datasets with Gaussian noise added to recruit numbers and fishing mortality rates ($N(0, \sigma^2_{stock,year})$, where $\sigma^2_{stock,year}$ is the stock- and year-specific standard deviation). The Gaussian noise is generated with stock- and year-specific coefficients of variation (CVs) in these parameters reported in the ICES reports, assuming CVs in mean fishing mortality rates represent those in age-specific rates (for stocks with standard deviations (SDs) being reported, CVs are computed using means and SDs). The CVs range from 3.3% to 75.1% for recruit numbers and from 2.4% to 46.1% for fishing mortality rates. For stocks with no uncertainty information being reported, CVs averaged across stocks and years were used.

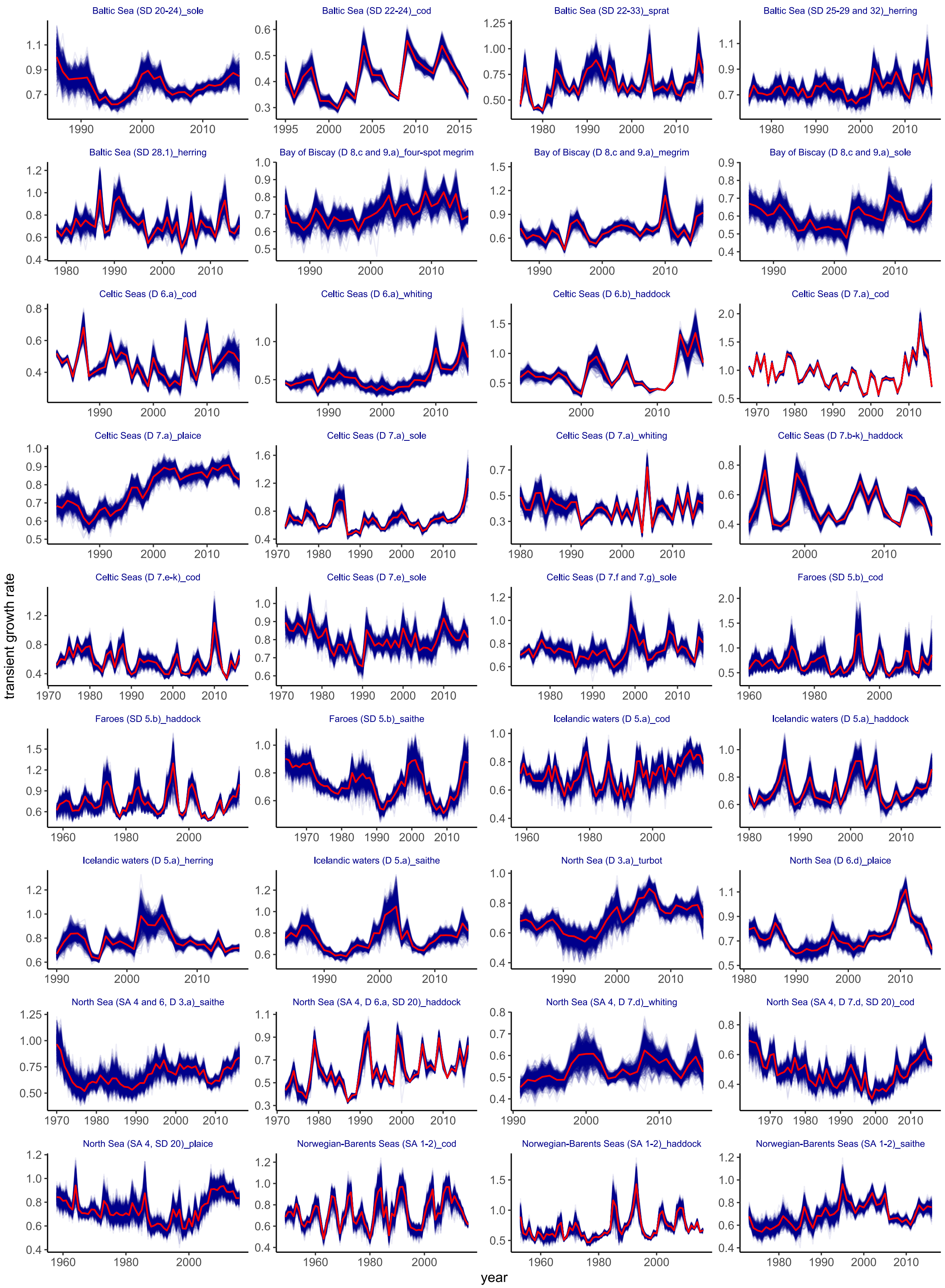
Baltic Sea (SD 20-24)_sole
Baltic Sea (SD 22-24)_cod
Baltic Sea (SD 22-33)_sprat
Baltic Sea (SD 25-29 and 32)_herring
Baltic Sea (SD 28.1)_herring
Bay of Biscay (D 8.c and 9.a)_four-spot megrim
Bay of Biscay (D 8.c and 9.a)_megrim
Bay of Biscay (D 8.c and 9.a)_sole
Celtic Seas (D 6.a)_cod
Celtic Seas (D 6.a)_whiting
Celtic Seas (D 6.b)_haddock
Celtic Seas (D 7.a)_cod
Celtic Seas (D 7.a)_plaice
Celtic Seas (D 7.a)_sole
Celtic Seas (D 7.a)_whiting
Celtic Seas (D 7.b-k)_haddock
Celtic Seas (D 7.e-k)_cod
Celtic Seas (D 7.e)_sole
Celtic Seas (D 7.f and 7.g)_sole
Faroes (SD 5.b)_cod
Faroes (SD 5.b)_haddock
Faroes (SD 5.b)_saithe
Icelandic waters (D 5.a)_cod
Icelandic waters (D 5.a)_haddock
Icelandic waters (D 5.a)_herring
Icelandic waters (D 5.a)_saithe
North Sea (D 3.a)_turbot
North Sea (D 6.d)_plaice
North Sea (SA 4 and 6, D 3.a)_saithe
North Sea (SA 4, D 6.a, SD 20)_haddock
North Sea (SA 4, D 7.d)_whiting
North Sea (SA 4, D 7.d, SD 20)_cod
North Sea (SA 4, SD 20)_plaice
Norwegian-Barents Seas (SA 1-2)_cod
Norwegian-Barents Seas (SA 1-2)_haddock
Norwegian-Barents Seas (SA 1-2)_saithe
transient growth rate
year

**Figure S5**. Trends in ecoregion-scale deviations in (a) relative fishing pressure, (b) relative spawner biomass, (c) mean spawner age, and (d) spawner age diversity of the recovering and non-recovering northeast Atlantic fish stocks during1946–2016. The deviations are estimated by generalized additive mixed effects modeling with year as a fixed effect and ecoregion, year within ecoregion, and stock as random intercepts.

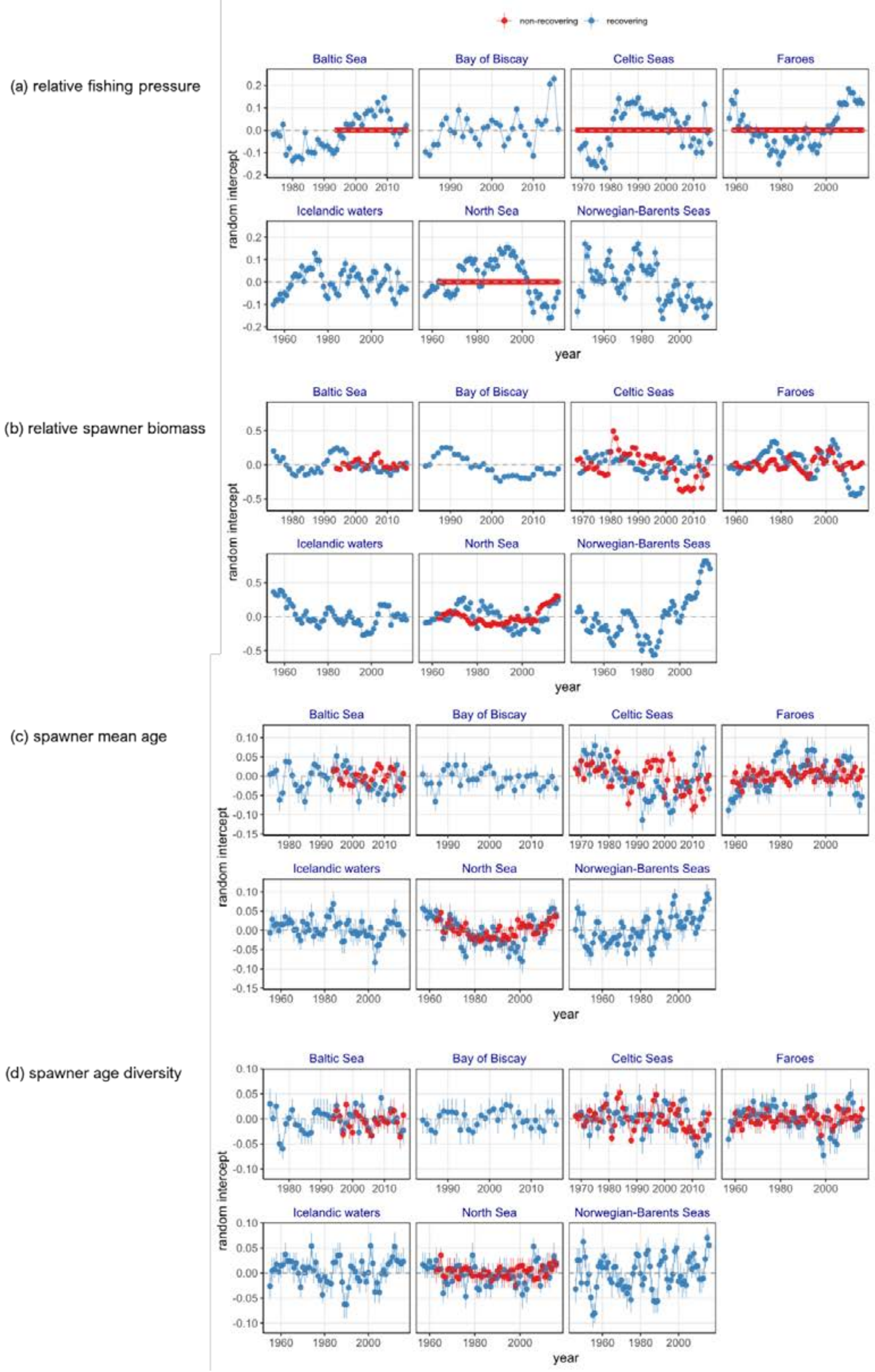

**Figure S6**. Trends in ecoregion-scale deviations in (a) transient population growth, (b) recruitment success, and (c) elasticity to recruitment success of the recovering and non-recovering northeast Atlantic fish stocks during1946–2016. The deviations are estimated by generalized additive mixed effects modeling with year as a fixed effect and ecoregion, year within ecoregion, and stock as random intercepts.

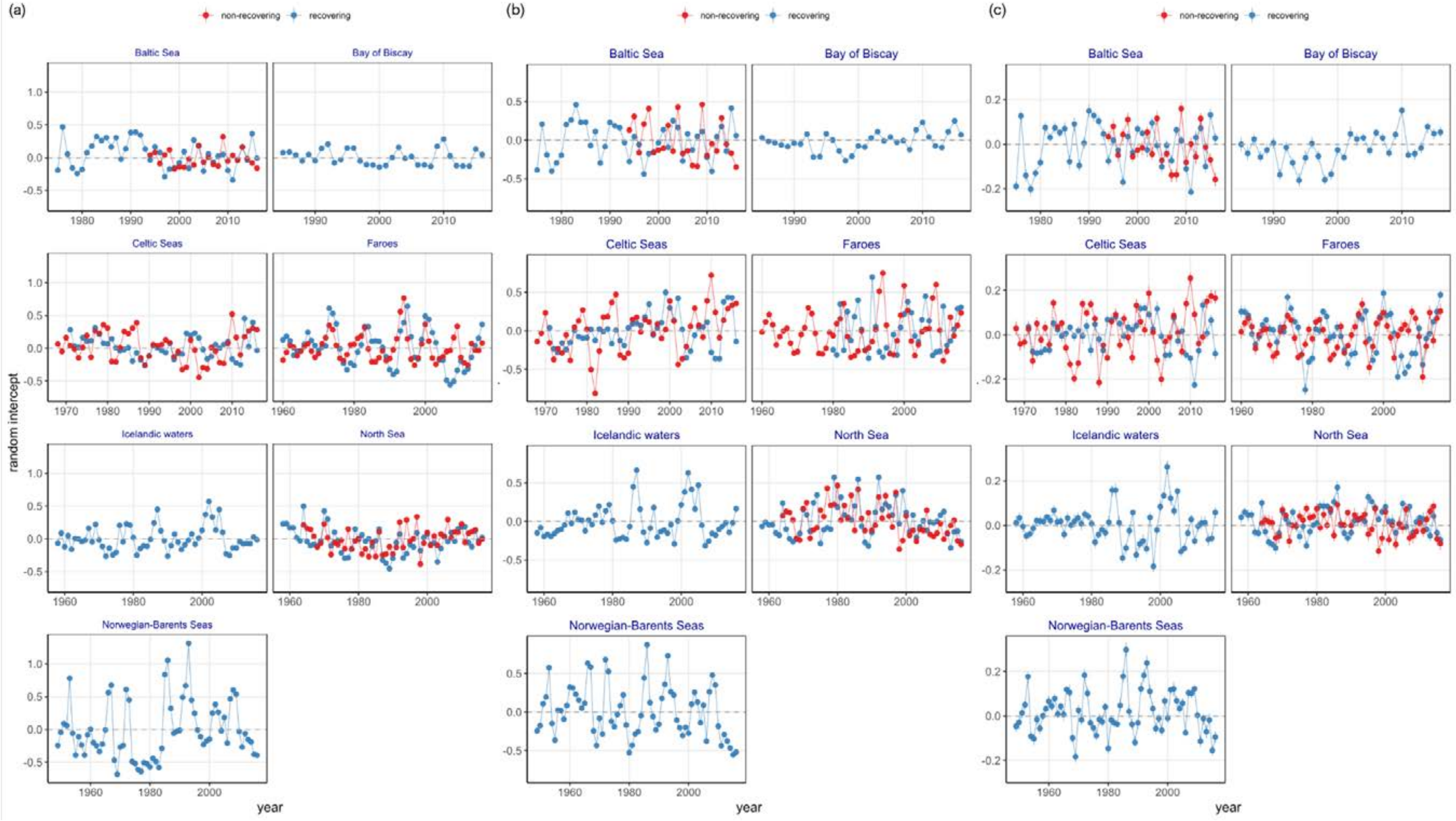

**Figure S7**. Parameter estimates (slopes and intercepts) from generalized additive mixed models with transient population growth as a response variable and year as a covariate. The models are fitted to each of 1000 simulated datasets generated with Gaussian noise added to recruit numbers and fishing mortality rate estimates taken from the International Council for the Exploration of the Sea (ICES) 2017 stock assessment reports prior to the computation of transient growths. Vertical dashed lines indicate medians. Red circles indicate parameter estimates (±95% confidence intervals) from the models fitted to the original abundance and demographic rate estimates taken from the reports.

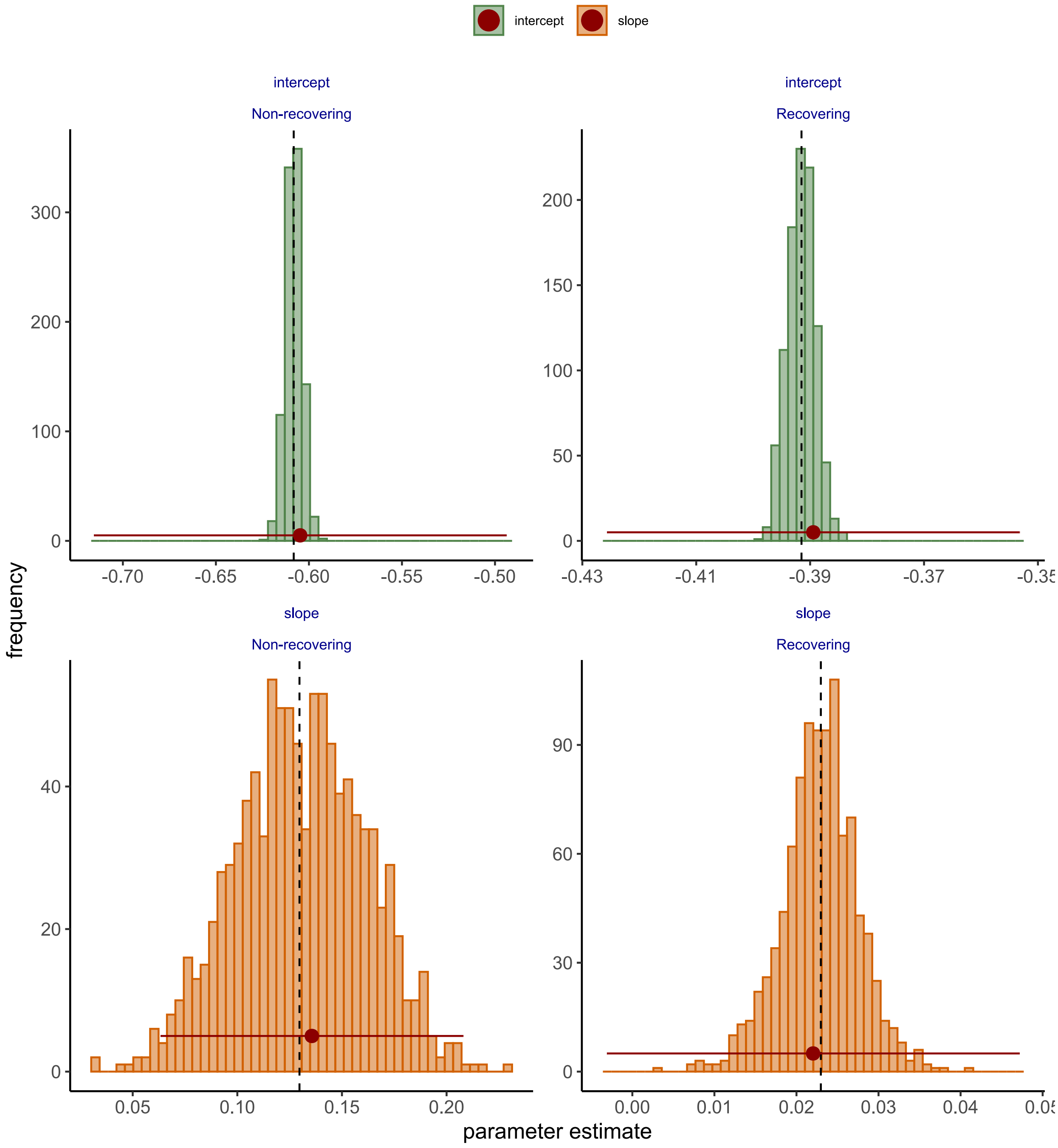

**Figure S8**. Trends in estimates of transient population growth relative to asymptotic population growth of 38 northeast Atlantic fish stocks during 1946–2016. Dashed grey horizontal lines indicate that transient growth is equal to asymptotic growth. SA indicates subarea; SD indicates subdivision; D indicates division within the northeast Atlantic ecoregions defined by the International Council for the Exploration of the Sea (ICES).

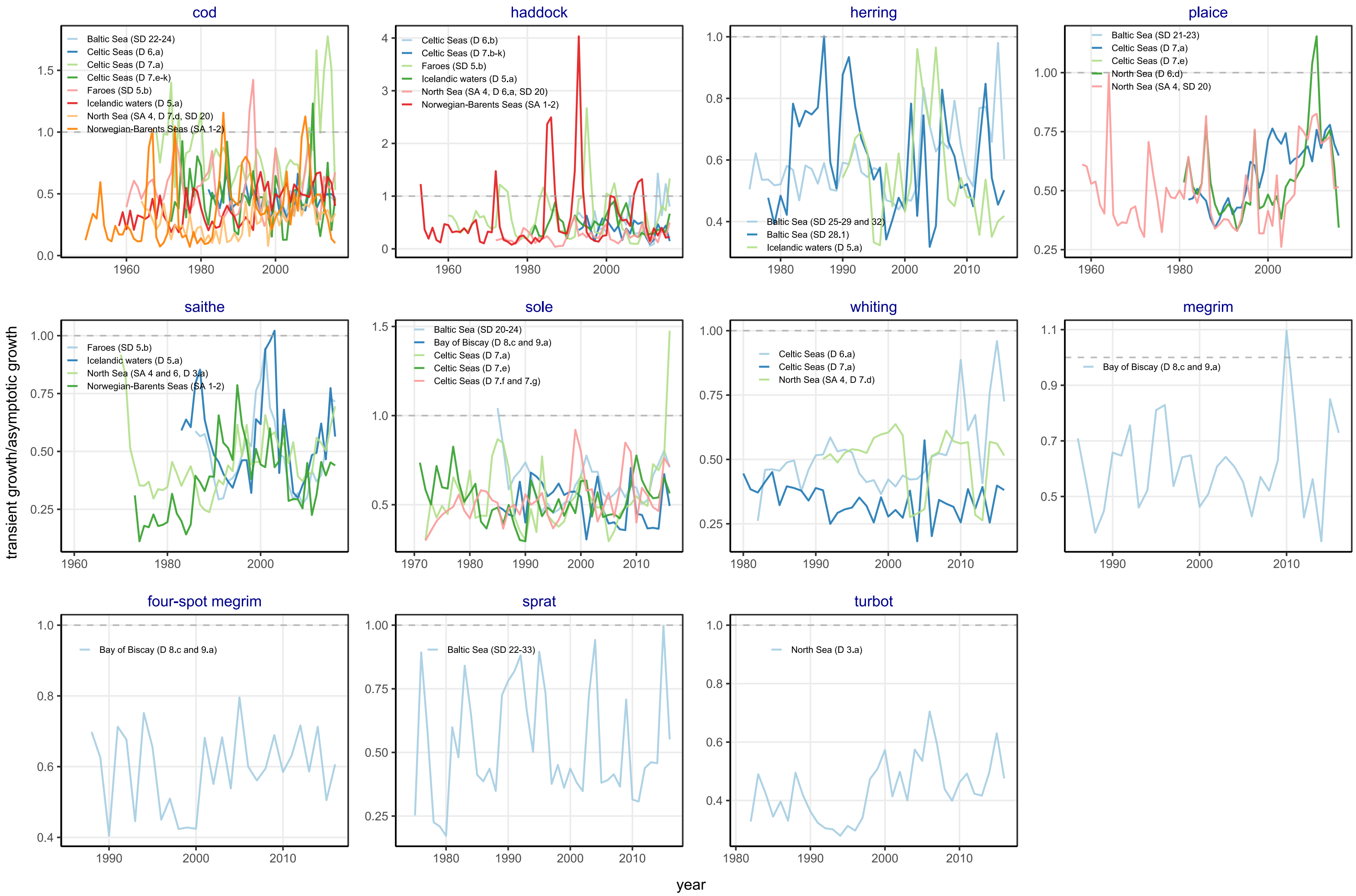

**Figure S9**. Relationships between observed and theoretical stable number-at-age estimates (by Leslie matrix modeling) of 38 northeast Atlantic fish stocks during 1946–2016. Dashed red lines indicate 1:1 lines. SA indicates subarea; SD indicates subdivision; D indicates division within the northeast Atlantic ecoregions defined by the International Council for the Exploration of the Sea (ICES).

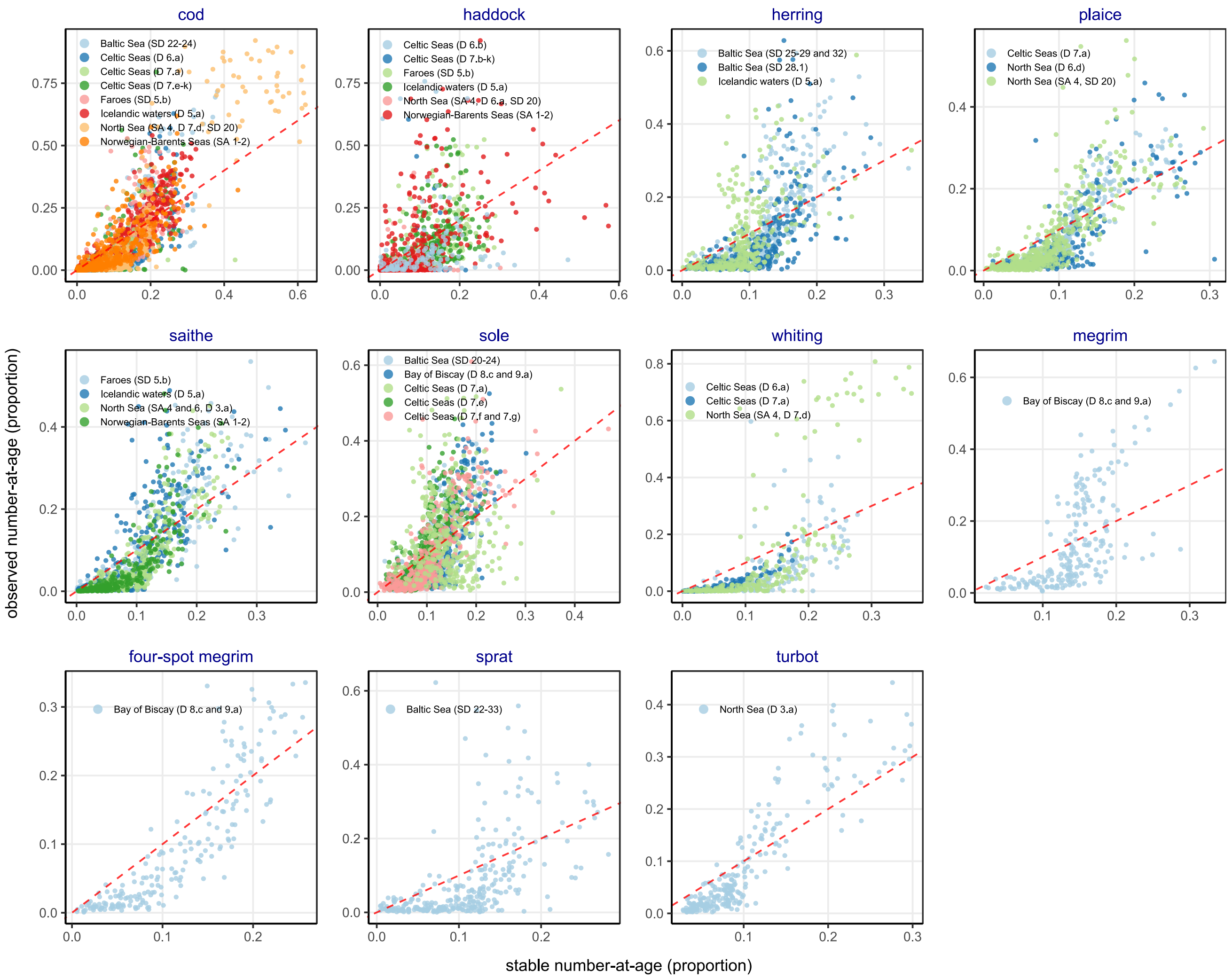

**Figure S10**. Age-specific fishing mortality rate estimates of 38 northeast Atlantic fish stocks during 1946–2016. The estimates are taken from the International Council for the Exploration of the Sea (ICES) 2017 stock assessments. Circles indicate annual mortality estimates. Solid lines indicate averages across years estimated by LOESS (locally estimated scatterplot smoothing). SA indicates subarea; SD indicates subdivision; D indicates division within the northeast Atlantic ecoregions defined by ICES.

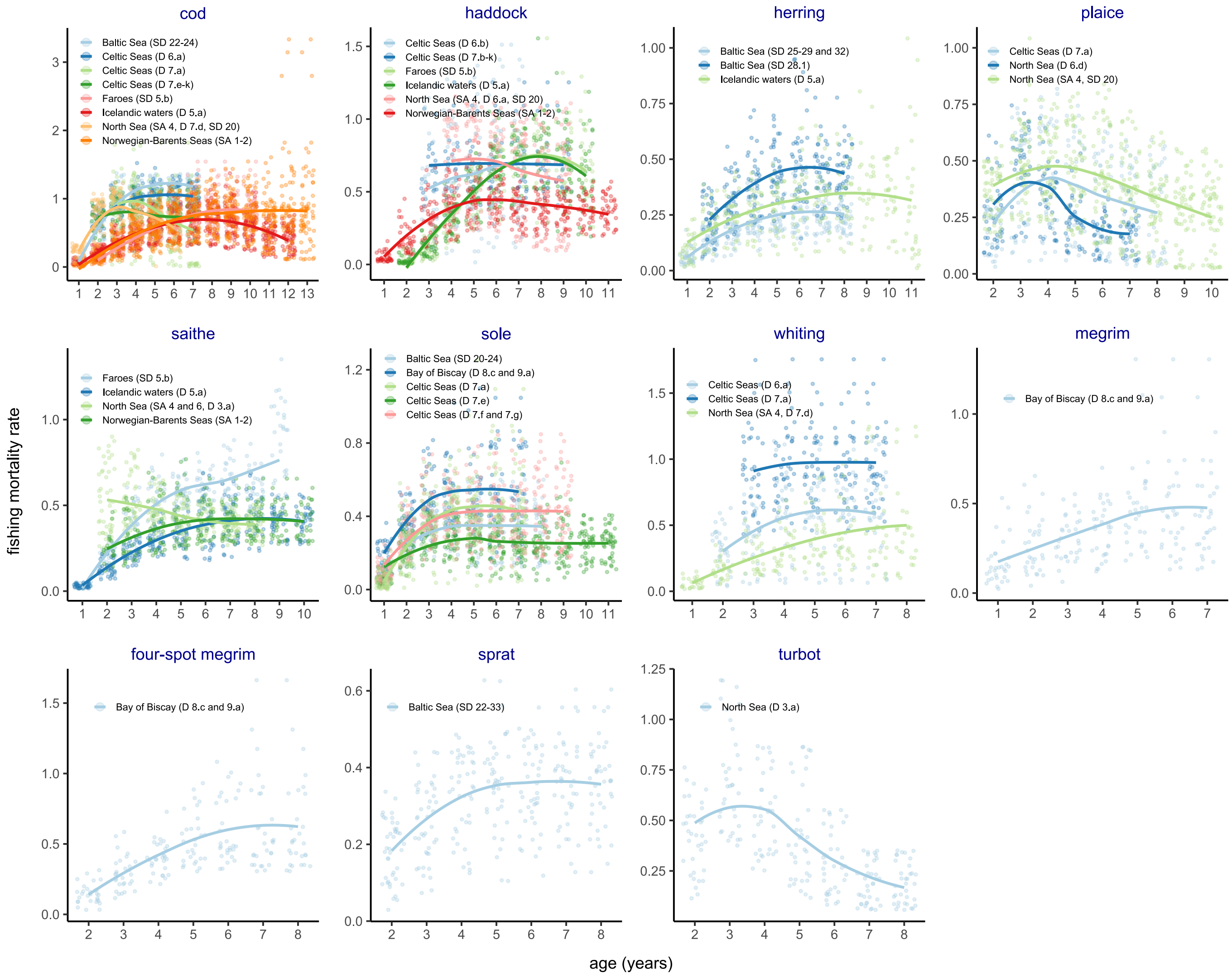

Figure S11. Ecoregion-scale trends in and covariates of per capita recruitment success (fecundity) of the northeast Atlantic fish stocks during 1946–2016. (a) Temporal trends. Solid lines and ribbons indicate temporal trends and 95% confidence intervals estimated by generalized additive mixed effects modeling with year as a fixed effect, and ecoregion, year within ecoregion, and stock as random effects. Circles indicate partial residuals estimated by the models. (b–e) Time-varying effects of spawner age structure, spawner biomass, fishing pressure. and climate condition. Covariates selected in the models are indicated in the y-axis labels. Color gradients and contour lines indicate the partial effects of covariates estimated (as tensor interaction terms) in the models. Warmer colors indicate higher values, and the ranges of the partial effects differ among the covariates and are indicated as values on the contour lines. Gray circles indicate back-transformed input data.

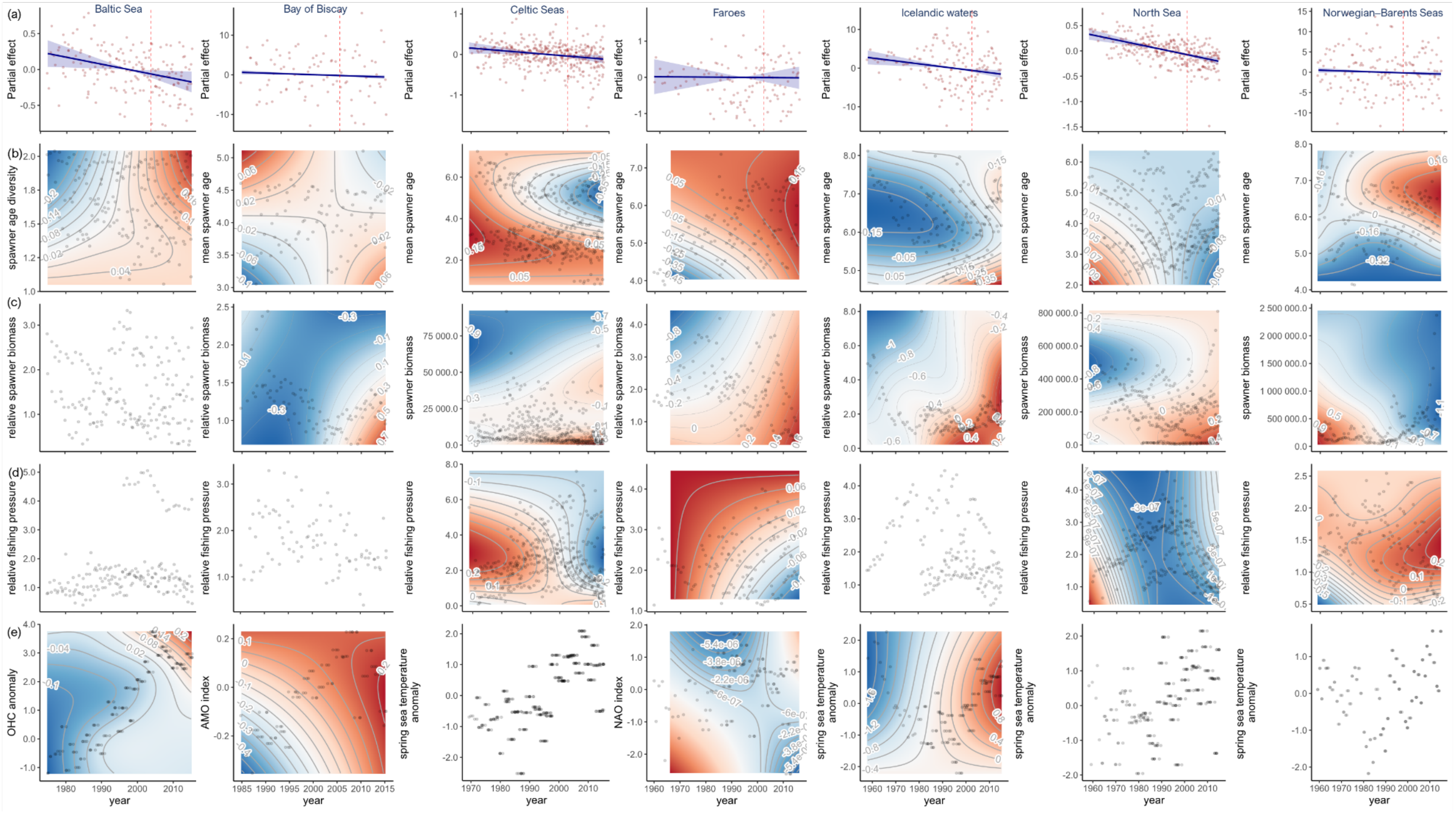

**Figure S12**. Correlations among regional and local climate indices for the northeast Atlantic Ocean used in the study. Warmer colors indicate negative correlations and cooler colors indicate positive correlations. The size of circles is in proportion to correlation coefficient.

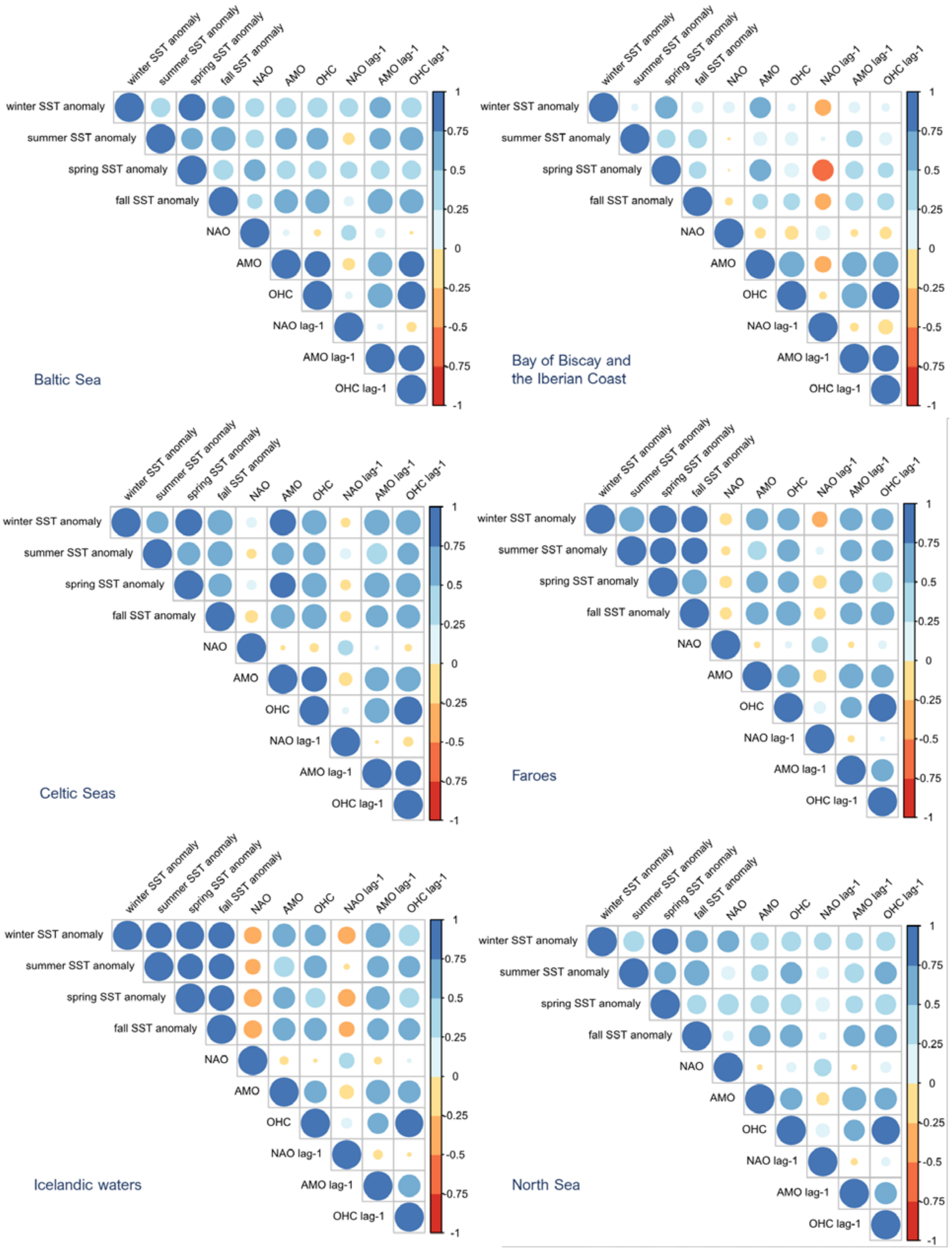

**Figure S12** (continued).

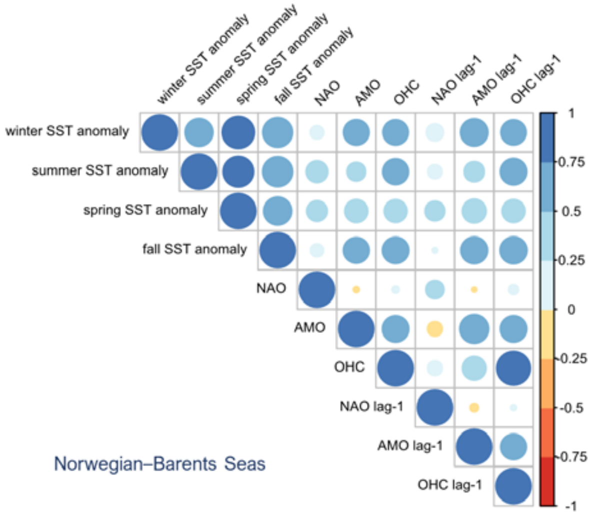

**Figure S13**. Regional trends and ecoregion-specific deviations in elasticities in (a) (post-recruitment) juvenile survival and (b) adult survival of the recovering and non-recovering northeast Atlantic fish stocks during 1946–2016. Solid lines and ribbons in regional trends indicate temporal trends and 95% confidence intervals estimated by generalized additive mixed effects modeling with year as a fixed effect, and ecoregion, year within ecoregion, and stock as random effects. Circles indicate partial residuals estimated from the models.

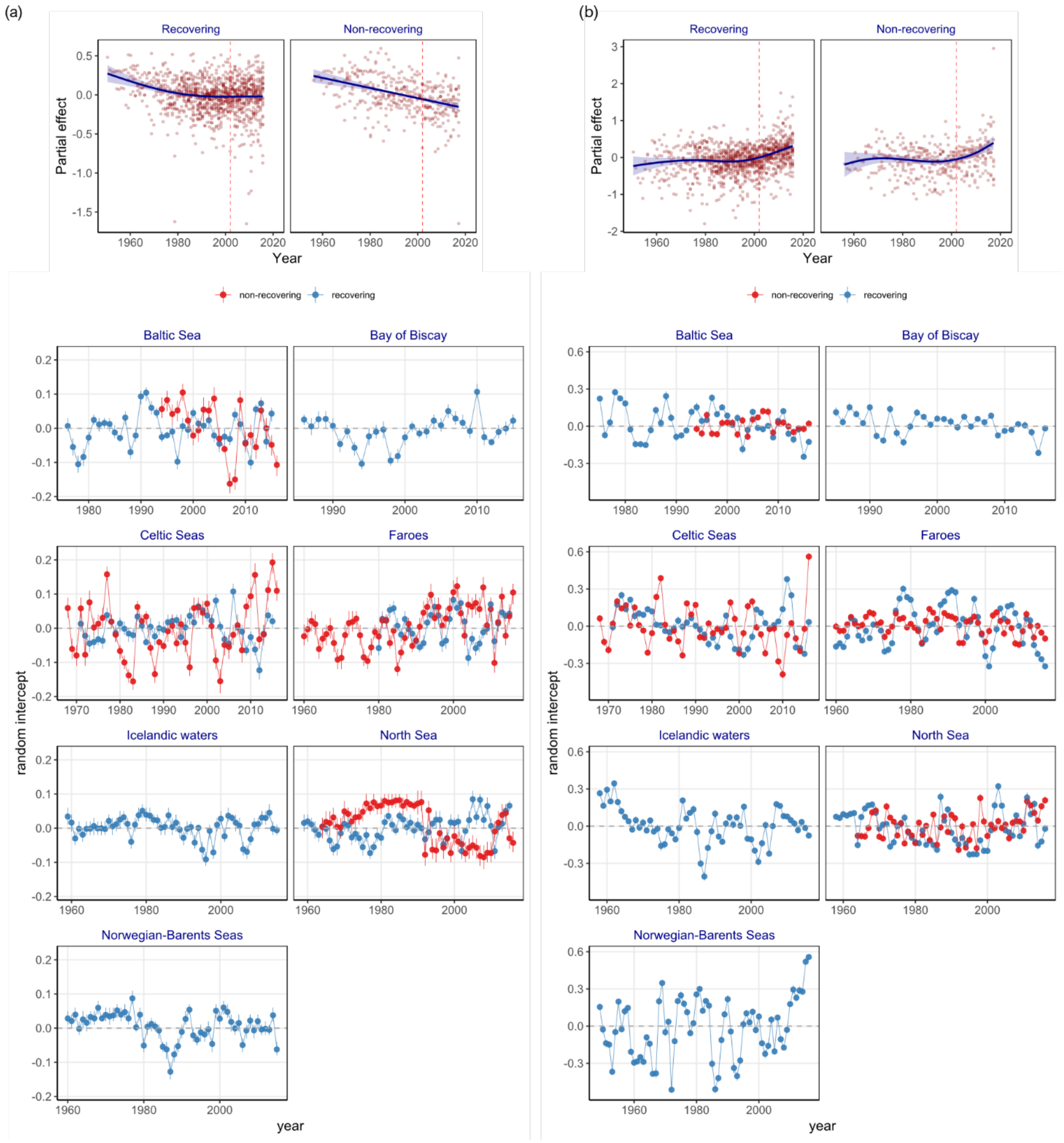

**Figure S14.** Ecoregion-scale trends in and covariates of elasticity to per capita recruitment success of the northeast Atlantic fish stocks during 1946–2016. (a) Temporal trends. Solid lines and ribbons indicate temporal trends and 95% confidence intervals estimated by generalized additive mixed effects modeling with year as a fixed effect, and ecoregion, year within ecoregion, and stock as random effects. Circles indicate partial residuals estimated by the models. (b–e) Time-varying effects of spawner age structure, spawner biomass, fishing pressure. and climate condition. Covariates selected in the models are indicated in the y-axis labels. Color gradients and contour lines indicate the partial effects of covariates estimated (as tensor interaction terms) in the models. Warmer colors indicate higher values, and the ranges of the partial effects differ among the covariates and are indicated as values on the contour lines. Gray circles indicate back-transformed input data.

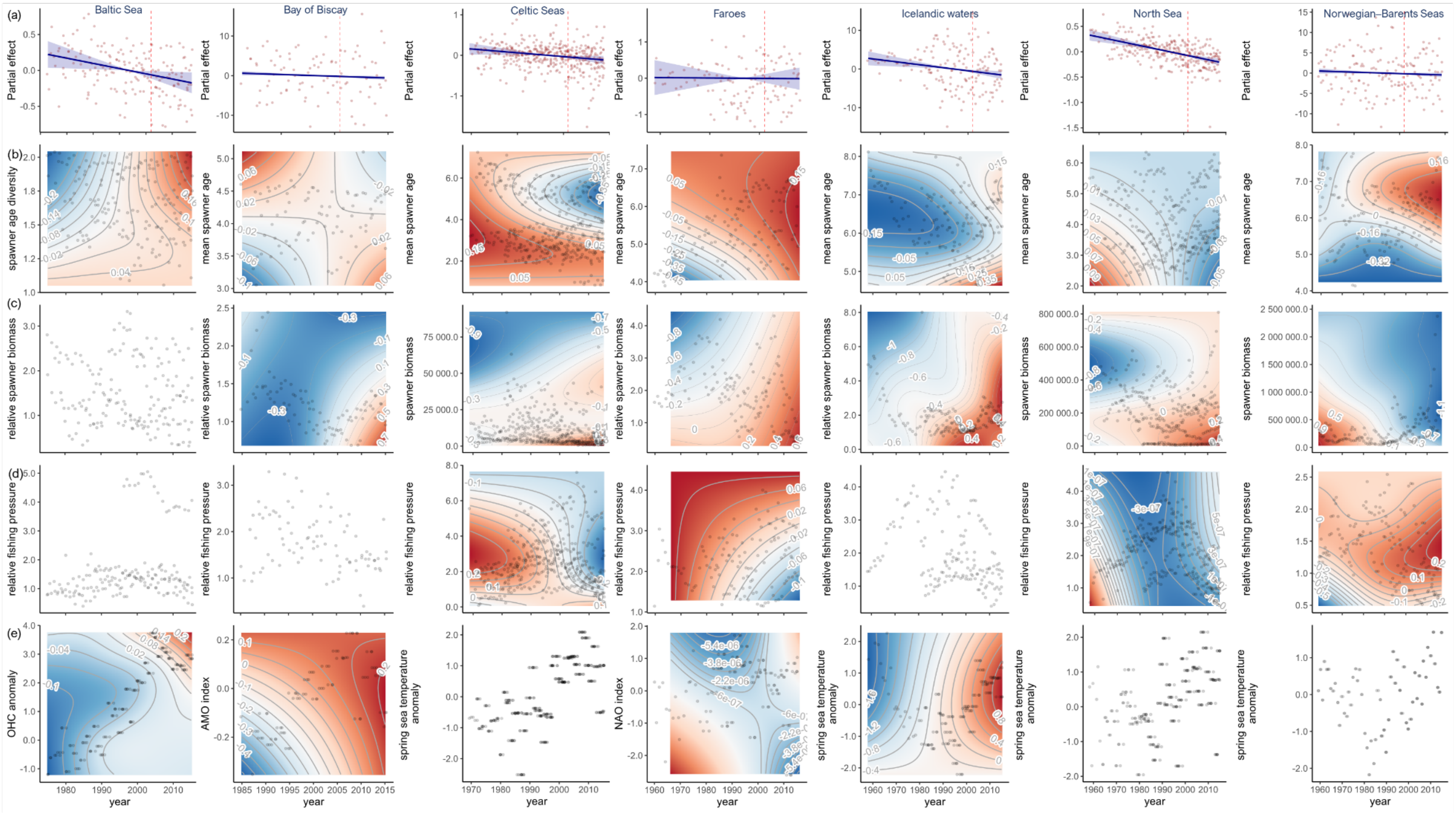
